# The Ran-binding protein Mog1 sustains Spt5 C-terminal phosphorylation by counteracting Spt4 in budding yeast

**DOI:** 10.64898/2026.11.13.766381

**Authors:** Joan Serrano-Quílez, Carme Nuño-Cabanes, Pablo Mulas-Saurí, Susana Rodríguez-Navarro

**Affiliations:** Instituto de Biomedicina de Valencia (IBV-CSIC), Valencia, Spain

## Abstract

Transcription elongation by RNA polymerase II (RNAPII) requires phosphorylation of the C-terminal repeat (CTR) of the elongation factor Spt5 by the Bur1-Bur2 kinase, yet how this modification is sustained remains unclear. Here we identify the conserved Ran cofactor Mog1 as a positive determinant of Spt5 CTR phosphorylation in budding yeast. Cells lacking Mog1 show reduced Spt5 phosphorylation and chromatin association, lower RNAPII CTD Ser2 phosphorylation and a previously unreported, guanine-reversible sensitivity to nucleotide depletion. These defects are mediated by Spt4, which accumulates in *mog1*Δ cells without a change in its nucleoplasmic distribution, and deleting *SPT4* restores Spt5 phosphorylation, H2B monoubiquitination and H3K4 trimethylation, and normal growth and morphology. Both Spt4-Spt5 subunits, but neither BUR kinase subunit, co-purify with Mog1. Deleting the demethylase *JHD2* restores H3K4me3 without rescuing growth, establishing the chromatin defects as consequences, not causes, of the elongation defect. Bur1 abundance and activation are unaffected in *mog1*Δ cells, and reducing BUR activity partially compensates rather than aggravates the loss of Mog1, indicating that Mog1 acts on the Spt4-Spt5 substrate; synthetic sickness with *ctk1*Δ places its contribution in early elongation. We propose that Spt4 imposes a limit on Spt5 CTR phosphorylation and that Mog1 counteracts this constraint.

**Graphical Abstract:** 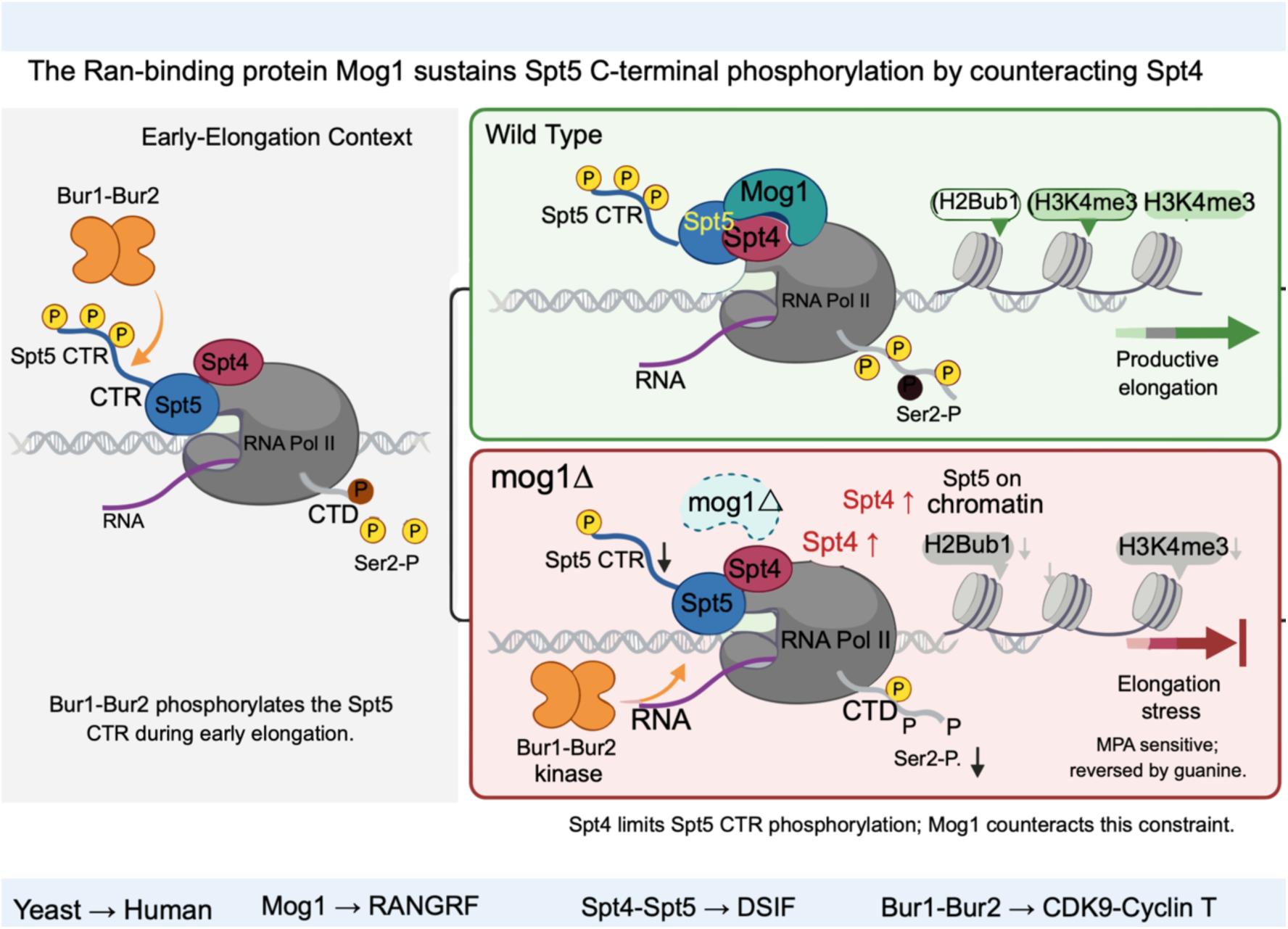

## INTRODUCTION

In eukaryotes, gene expression requires the precise coordination of transcription initiation, elongation and termination with co-transcriptional RNA processing and chromatin modification (1, 2). The elongation phase of RNA polymerase II (RNAPII) transcription is subject to extensive regulation that ensures productive polymerase progression across gene bodies while coupling mRNA synthesis to downstream events. A central control mechanism is the dynamic phosphorylation of the carboxy-terminal domain (CTD) of the largest RNAPII subunit, Rpb1, particularly at serine 2 (Ser2) of its heptapeptide repeats. Progressive Ser2 phosphorylation marks the transition to productive elongation and provides a platform for the recruitment of elongation factors, chromatin modifiers and RNA-processing machinery (3, 4).

Among the elongation factors, the conserved Spt4-Spt5 complex, known as DSIF in metazoans, plays a central role (5, 6). Spt5, the only elongation factor conserved across all three domains of life, associates directly with RNAPII shortly after initiation and travels with the polymerase throughout the gene body, controlling both the rate and the processivity of RNA synthesis (7, 8). Its partner Spt4 is non-essential, *spt4*Δ cells are viable but show altered transcriptional responses and sensitivity to elongation stress (6, 9, 10). A key feature of productive elongation is phosphorylation of the Spt5 C-terminal repeat (CTR) by the cyclin-dependent kinase Bur1 in complex with its cyclin Bur2 (11, 12). In metazoans, the equivalent modification by the Bur1 orthologue CDK9 triggers the switch from promoter-proximal pausing to productive elongation, converting Spt4-Spt5 from a pausing factor into a processivity factor (13, 14).

Bur1-dependent Spt5 CTR phosphorylation is directly coupled to the chromatin modifications that mark actively transcribed genes. It stimulates recruitment of the PAF complex (Paf1C) and, through activation of the Rad6/Bre1 ubiquitin-conjugating system, promotes histone H2B monoubiquitination (H2Bub1), which is in turn required for H3K4 trimethylation (H3K4me3) by COMPASS (15–17). Because these modifications are interdependent, disruption at any early step propagates downstream and reshapes chromatin states across gene bodies (18). Ser2 phosphorylation is subsequently reinforced during late elongation by the kinase Ctk1, which acts downstream of Bur1. Together, Bur1 and Ctk1 constitute the yeast functional equivalent of the metazoan CDK9-CDK12 axis, with Bur1 acting preferentially during early elongation and Ctk1 sustaining CTD modification thereafter (3, 19). At the 3’ end of genes, dephosphorylation of the Spt5 CTR by the PP1 phosphatase Glc7 releases Paf1C from RNAPII, a step required for efficient termination (20).

Mog1 was originally identified in *Saccharomyces cerevisiae* as a Ran-binding protein required for nuclear protein import (21, 22). It binds directly the GTP-bound form of the Ran homologue Gsp1, and the 1.9 A crystal structure of yeast Mog1 revealed a compact fold distinct from other Ran-binding proteins, with a surface cluster of conserved acidic residues mediating the Ran interaction (23). Mog1 is structurally and functionally conserved through evolution. Its human orthologue RANGRF supports nuclear import and additionally regulates trafficking of Nav1.5, the pore-forming subunit of the cardiac sodium channel, linking Mog1 function to human physiology (24, 25). In addition to transport, research conducted in our laboratory has demonstrated that *mog1Δ* cells exhibit deficiencies in global transcription and mRNA export. These cells also show a significant decrease in H2Bub1 and H3K4me3 levels, along with impaired chromatin recruitment of Rad6, Bre1, and Rtf1, which are essential components of the H2Bub1 pathway (26, 27). Consistent with a chromatin-linked role, Mog1 interacts with Shg1, a subunit of the COMPASS-associated Set1 complex, through a surface region of the protein (26, 28). These observations positioned Mog1 as an upstream regulator of transcription-coupled chromatin modification; however, the specific stage at which it functions within the elongation cycle remains undefined.

The loss of H2Bub1 in *mog1Δ* cells is particularly informative (26). Because H2Bub1 depends on the Bur1-Spt5-Paf1C axis, its reduction raises the possibility that Mog1 acts at the level of Spt5 phosphorylation or Bur1 activity rather than solely on the downstream chromatin-modifying enzymes. Here we test this hypothesis. We show that loss of Mog1 reduces Spt5 CTR phosphorylation, Spt5 association with transcribed chromatin and RNAPII Ser2 phosphorylation, and confers a previously unreported, guanine-reversible sensitivity to elongation stress. We further show that these defects require Spt4, and that preventing H3K4me3 removal by deleting the demethylase *JHD2* restores the mark but not growth, establishing the chromatin marks as a consequence rather than a cause of the elongation defect. Consistent with a robust rather than single-contact engagement, neither a surface nor a Ran-binding-face substitution of Mog1 abolishes its elongation function. The notion that Spt5 CTR phosphorylation is set not only by kinase activity but also by factors that control access to the substrate has recently emerged in metazoans, where LEDGF/p75 prevents phosphorylation of the SPT5 phosphorylation-site-rich domain and thereby promotes promoter-proximal pausing (29), and where phosphorylation of distinct SPT5 regions by CDK9 coordinately controls pause release, elongation speed and termination (30). The work presented here describes a yeast counterpart of this regulatory layer, in which Spt4 limits and Mog1 permits modification of the Spt5 CTR.

## MATERIALS AND METHODS

### Reagents

Cells were routinely grown at 30 °C with shaking at 160 rpm in YPD (1% yeast extract, 2% peptone, 2% glucose) or in synthetic complete medium (SC; 0.17% yeast nitrogen base, 0.5% ammonium sulphate, 2% glucose and the appropriate drop-out amino-acid mix). Solid media contained 2.4% agar. For antibiotic selection, YPD was supplemented with 200 µg/mL G418 or 300 µg/mL hygromycin B. Where indicated, plates contained 6-azauracil (6-AU) or mycophenolic acid (MPA) at the concentrations specified in the figure legends; parallel plates were supplemented with guanine (0.1 mg/mL) to confirm that MPA phenotypes reflected nucleotide depletion rather than off-target effects. Whole-cell extracts were prepared by trichloroacetic acid (TCA) precipitation. Proteins were resolved by SDS-PAGE on gels of 6-15% acrylamide and transferred to nitrocellulose membranes (0.45 µm; 0.2 µm for histone modifications). For the separation of phosphorylated species, Phos-tag SDS-PAGE was performed with resolving gels supplemented with 20 µM Phos-tag and 20 µM MnCl₂ (31), and gels were equilibrated in transfer buffer containing 1 mM EDTA to chelate Mn²⁺ before transfer to PVDF membranes. Membranes were blocked in 5% skimmed milk or 1% BSA in TBS-T and probed with the following primary antibodies: anti-Ser2P of the RNAPII CTD (rat; ChromoTek), anti-TAP (rabbit; Thermo Fisher), anti-HA (mouse; Roche), anti-GFP (mouse; Roche), anti-H2Bub (rabbit; Cell Signaling), anti-H3K4me3 (rabbit; Abcam), anti-histone H3 (rabbit; Abcam) and anti-Pgk1 (mouse; Invitrogen), the latter two serving as loading controls. Detection was performed with infrared-dye-conjugated secondary antibodies (IRDye 800CW anti-rabbit and IRDye 680CW anti-mouse; LI-COR) on an Odyssey Imaging System, with the exception of the rat anti-Ser2P antibody, which was detected with an HRP-conjugated anti-rat secondary antibody (Dako) and developed by chemiluminescence (ECL) on an Amersham Imager.

Mog1-TAP complexes were purified under native conditions in lysis buffer (50 mM Tris-HCl pH 7.5, 100 mM NaCl, 1.5 mM MgCl₂, 0.15% NP-40) supplemented with DTT, PMSF and protease-inhibitor cocktail. Clarified extracts were incubated with IgG-Sepharose; bound complexes were released by TEV cleavage, then bound to calmodulin affinity resin and eluted with EGTA. Final eluates were precipitated with TCA. ChIP enrichment was quantified by qPCR (TB Green, Takara Bio) on a QuantStudio Real-Time PCR System. Chromatin was sheared in a Bioruptor sonicator (Diagenode) to an average fragment size of ∼300 bp.

### Biological Resources

All *Saccharomyces cerevisiae* strains used in this study are derivatives of the BY4741/BY4742 or W303 backgrounds and are listed in Supplementary Table S1, together with the plasmids and oligonucleotides used (Supplementary Tables S2 and S3). Amino acids were supplemented according to auxotrophic requirements. Gene deletions and C-terminal epitope tagging were generated by homologous recombination using PCR-amplified cassettes (32, 33), introduced by a standard lithium-acetate/PEG transformation (34) and selected on the appropriate SC drop-out plates or, for dominant markers, on antibiotic-containing YPD. Correct integration was confirmed by diagnostic PCR and, where appropriate, by Sanger sequencing or immunoblotting.

Strains carrying multiple mutations were obtained by crossing haploids of opposite mating type on YPD, selecting diploids and inducing sporulation; tetrads were digested with Zymolyase-20T and dissected on YPD with an MSM 300 micromanipulator (Singer Instruments), and spores were genotyped by replica plating onto selective media. The *mog1-E65K* allele was introduced at the endogenous *MOG1* locus by CRISPR-Cas9 as described in (35), avoiding the overexpression artefacts of plasmid-based complementation. A locus-specific single-guide RNA, designed with CRISPOR (36), was expressed from a linearised pMEL10 plasmid and co-transformed into Cas9-expressing cells together with a 120-nt double-stranded donor carrying the E65K substitution. Edited clones were identified by PCR and Sanger sequencing of the targeted region.

For growth assays, overnight cultures were diluted to OD₆₀₀ = 0.1 and ten-fold serial dilutions were spotted onto the indicated media. Plates were incubated at the indicated temperatures and growth was scored after 2–4 days.

Whole-cell extracts were prepared from cultures grown in SC at 30 °C (or at the temperatures and conditions indicated). Cells harvested at the exponential phase were precipitated in 20% TCA, essentially as described (37), neutralised in 1 M Tris, resuspended in SDS loading buffer and lysed with glass beads in a Precellys Evolution homogeniser (Bertin Technologies), followed by boiling. Transfer efficiency was monitored by Ponceau staining. When quantified, band intensities were measured in ImageJ and normalised as indicated in the figure legends.

ChIP was performed from 50 mL cultures grown to OD₆₀₀ ≈ 0.7-0.8 and crosslinked with 1% formaldehyde for 20 min at room temperature; crosslinking was quenched with 125 mM glycine. Cells were washed in cold TBS, lysed with glass beads, and chromatin was sheared as described above. After clarification, an aliquot was retained as input and the remainder was used for immunoprecipitation with magnetic beads bound directly to IgG (for TAP-tagged proteins) or pre-coated with the appropriate antibody, for at least 2 h at 4 °C. Beads were washed sequentially with lysis buffer, high-salt lysis buffer, washing buffer and TE, and bound material was eluted. Input and immunoprecipitated samples were incubated overnight at 65 °C to reverse crosslinks, treated with proteinase K, and DNA was purified by phenol-chloroform extraction and ethanol precipitation. Enrichment was expressed as IP/Input and normalised to an intergenic region. Primer sequences are listed in Supplementary Table S3.

Mog1-TAP complexes were purified by a two-step TAP procedure (38, 39) with minor modifications. Cultures of 2-4 L were harvested at OD₆₀₀ ≈ 2.5, washed, and lysed in a BeadBeater (Fritsch) with glass beads. Final eluates were analysed by liquid chromatography-tandem mass spectrometry (LC-MS/MS) at the Servei Central de Suport a la Investigació Experimental (SCSIE), University of Valencia. Identified proteins, together with the sequence coverage obtained for the Mog1 bait, are listed in Table S4. Cells expressing genomically tagged Spt4-GFP, either alone or together with Net1-mCherry, Pus1-mCherry or Sec63-mCherry (expressed from centromeric plasmids), as well as cells imaged using brightfield microscopy alone, were grown in SC medium to mid-exponential phase (OD₆₀₀ ≈ 0.3-0.5). Cells were collected by brief centrifugation, resuspended in a small volume of SC medium, and immediately imaged on a Zeiss LSM 980 confocal microscope equipped with a 63× oil-immersion objective using ZEN Blue software. GFP was excited at 488 nm and mCherry at 561 nm; emission was collected in the appropriate spectral windows. Brightfield images were also acquired, either independently or alongside GFP fluorescence images. Z-stacks were acquired at 0.3 µm intervals spanning the full cell volume, and maximum-intensity projections were generated using ZEN Blue. Images were processed and assembled in Fiji (ImageJ).

To interpret the Mog1 point and deletion mutants, the corresponding residues were mapped onto the experimental crystal structure of yeast Mog1 (PDB 1EQ6) (23). Secondary structure and per-residue relative solvent accessibility were computed with Biotite (40), and side-chain burial was assessed to distinguish surface-exposed from core positions. Structures were visualised in PyMOL (41).

### Statistical Analyses

All analyses were performed in R (v4.4.3). Unless otherwise stated, experiments were carried out with at least three biological replicates and a normal distribution of the values was assumed. Means were compared by unpaired two-tailed Student’s t-test (two groups) or by one-way ANOVA followed by Tukey’s multiple-comparison test (three or more groups); Welch’s correction was applied when the compared groups were heteroscedastic. The specific test used is indicated in each figure legend, and P < 0.05 was considered statistically significant.

### Web Sites/Data Base Referencing

The following resources were used: PDB entry 1EQ6 (www.rcsb.org); CRISPOR sgRNA design tool (crispor.tefor.net); ImageJ/Fiji (imagej.nih.gov); R statistical computing environment (www.r-project.org).

Use of AI-assisted technologies. The authors used an AI assistant (Claude, Anthropic) for critical reading of the manuscript and its figures, for literature searching, and for language editing. All data, analyses and interpretations were generated and verified by the authors, who take full responsibility for the content of this work. The AI tool is not an author and made no independent contribution to the design, execution or interpretation of the experiments.

## RESULTS

### Mog1 sustains Spt5 phosphorylation and its chromatin engagement during elongation

We previously found that Mog1 co-purifies with the essential elongation factor Spt5 and that cells lacking Mog1 lose the elongation-associated chromatin marks H2B monoubiquitination (H2Bub1) and H3K4 trimethylation (H3K4me3) (26). These observations suggested that Mog1 contributes to RNA polymerase II (RNAPII) transcription elongation, but the underlying molecular basis remained undefined. We first asked whether Mog1 is required for growth when elongation is challenged. As already shown, *mog1Δ* cells grow slightly slower than wild-type on rich medium ((26), Figure 2B). Upon treatment with mycophenolic acid (MPA), an inhibitor of IMP dehydrogenase that limits GTP availability and imposes transcription elongation stress (42), *mog1Δ* cells displayed reduced growth, indicating that Mog1 becomes important for proliferation when nucleotide supply is limiting (Figure 1A). This sensitivity was fully reversed by guanine supplementation (Figure 1A), confirming that it reflects canonical GTP-depletion-mediated transcriptional stress rather than off-target effects.

**Figure 1.**
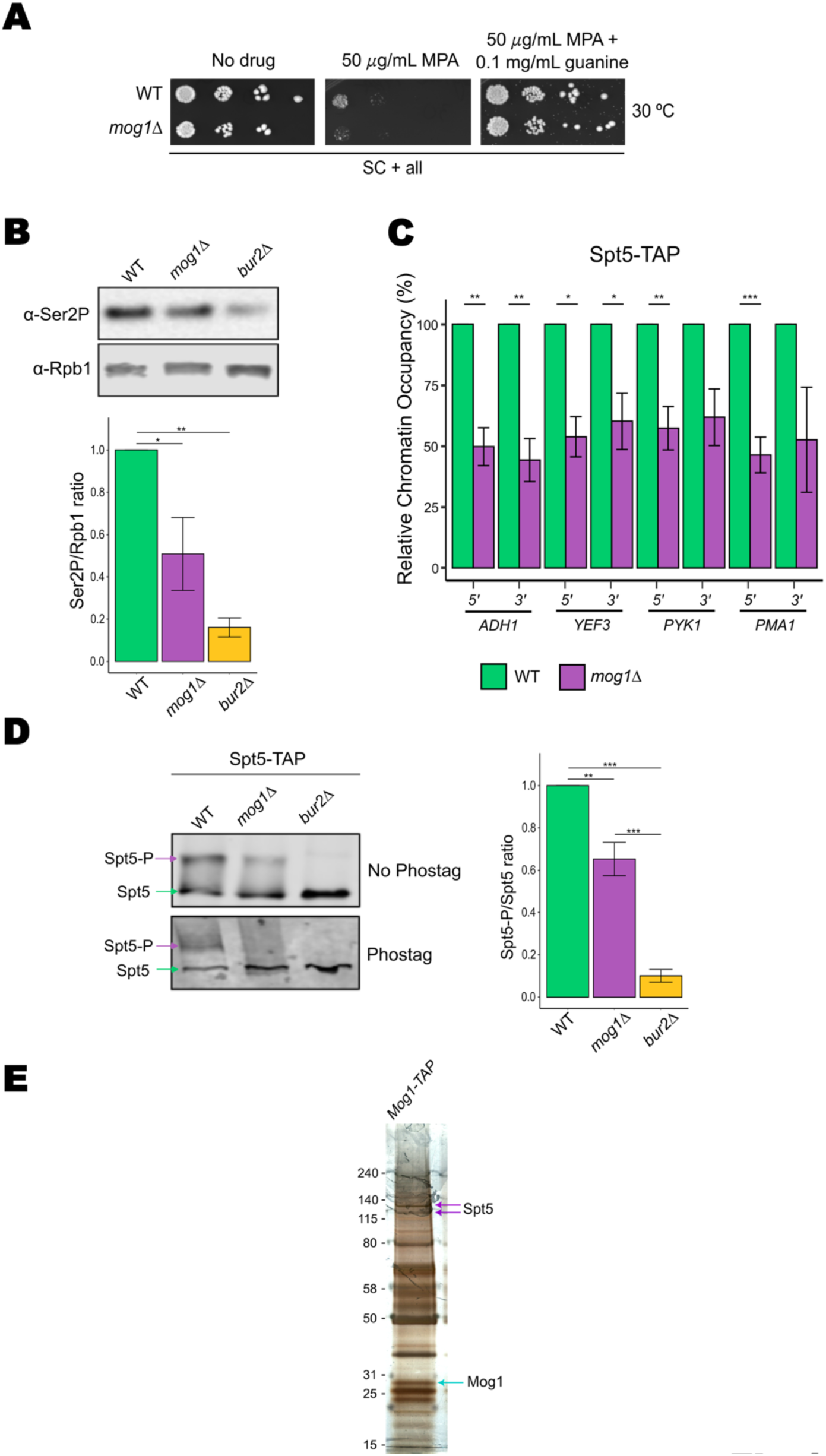
Loss of Mog1 impairs transcription elongation and reduces Spt5 phosphorylation, Spt5 chromatin association and RNAPII Ser2 phosphorylation, and Mog1 co-purifies with Spt5. (A) Ten-fold serial dilution spot assays of wild-type and *mog1Δ* cells on SC medium at 30 °C, without drug or containing 50 µg/mL mycophenolic acid (MPA); a parallel MPA plate was supplemented with 0.1 mg/mL guanine to confirm that the phenotype reflects GTP depletion. *mog1Δ* cells show a marked, guanine-reversible MPA sensitivity. (B) Western blot analysis of total phosphorylated RNAPII CTD at serine 2 (Ser2-P) in wild-type (WT), *mog1Δ* and *bur2Δ* strains. Rpb1 was used as loading control. Quantification of Ser2-P/Rpb1 signal intensity is shown on the right, normalized to WT. Data represent mean ± standard error from n = 3 biological replicates. Statistical significance was determined using one-way ANOVA followed by Tukey’s multiple comparison test. (C) ChIP-qPCR analysis of TAP-tagged Spt5 occupancy at the 5′ and 3′ coding regions of *ADH1, YEF3, PYK1* and *PMA1* in WT and *mog1Δ* strains. Relative enrichment was calculated as the IP/Input ratio and normalized to an intergenic region. Bar plots show mean ± standard error from n = 4 biological replicates. Statistical significance was assessed using unpaired two-tailed Student’s t-tests. (D) Western blot analysis of Spt5 phosphorylation in WT, *mog1Δ* and *bur2Δ* strains. Upper panel shows Spt5-TAP under standard SDS–PAGE; lower panel displays Phos-tag electrophoresis. Green arrows indicate unphosphorylated Spt5; purple arrows indicate the phosphorylated form (Spt5-P). Quantification (right) shows Spt5-P/Spt5 ratios normalized to WT. Values represent mean ± standard error from n = 3 biological replicates, quantified on two independent membranes. Statistical significance was determined using one-way ANOVA followed by Tukey’s multiple comparison test. The identity of the upper phosphorylated band was previously validated by λ-phosphatase treatment. For all panels, values are shown relative to WT (=1), with standard errors propagated using first-order Taylor expansion (delta method). *P = 0.01–0.05; **P < 0.01; ***P < 0.001. (E) Silver-stained SDS-PAGE of the Mog1-TAP eluate. Arrowheads mark the two species migrating above 100 kDa that were excised and identified as Spt5 by LC-MS/MS, and the band corresponding to the Mog1-TAP bait. The two Spt5 species are compatible with the unphosphorylated and phosphorylated forms detected in (D); the modification state of the slower-migrating species was not determined.

**Figure 2.**
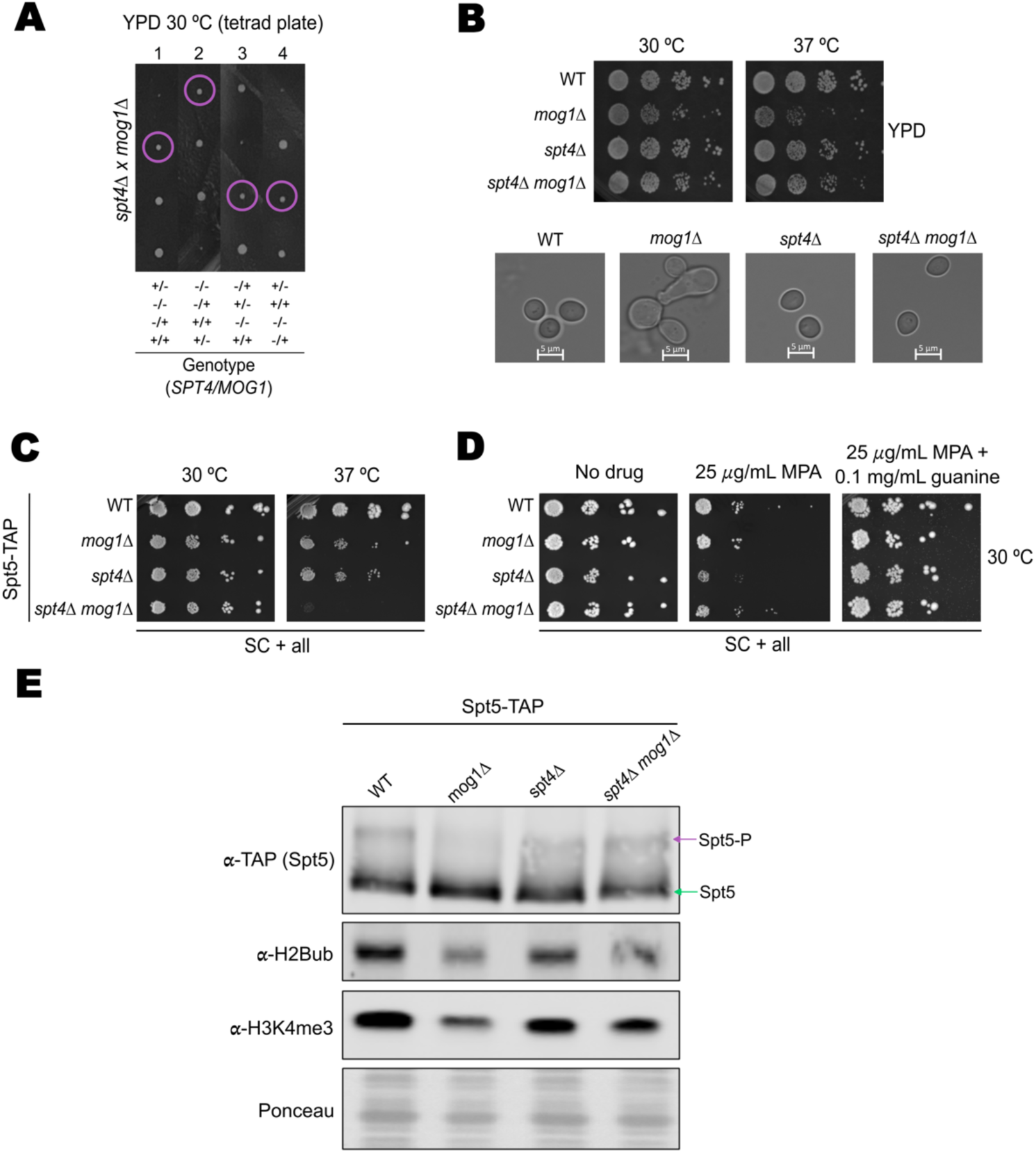
Deletion of SPT4 suppresses the growth and molecular defects of *mog1Δ*. (A) Tetrad dissection of the *spt4Δ × mog1Δ* cross. Each vertical column corresponds to a single tetrad, and each horizontal row represents one meiotic product. Colonies were grown on YPD at 30 °C for 3 days. Genotypes are indicated below using “+” for wild-type and “–” for deletion alleles as follows: +/+ = *SPT4 MOG1* (wild type), –/+ = *spt4Δ MOG1,* +/– = *SPT4 mog1Δ,* –/– = *spt4Δ mog1Δ.* This layout allows direct comparison of colony size among the four genotypic classes within each tetrad. (B) Tenfold serial dilution spot assays of the indicated strains. Exponentially growing cultures were spotted on YPD medium (30 °C and 37 °C) . Plates were incubated for 2–3 days before imaging. Brightfield microscopy images of wild type, *mog1Δ, spt4Δ,* and *spt4Δ mog1Δ* cells are shown below the spot assays. *mog1Δ* cells exhibit an altered morphology characterized by enlarged and elongated cells, whereas deletion of SPT4 restores a morphology resembling that of wild type cells. Scale bars, 5 µm. (C) Spot assays on SC medium in the Spt5-TAP background at 30 °C and 37 °C: *spt4Δmog1Δ* is synthetically lethal at 37 °C. (D) Spot assays on SC with MPA at 30 °C, with and without guanine, in the Spt5-TAP background: spt4Δ relieves the MPA sensitivity of mog1Δ. (E) Immunoblot of Spt5 phosphorylation and the elongation-associated chromatin marks H2Bub1 and H3K4me3 in wild-type, *mog1Δ, spt4Δ* and *spt4Δmog1Δ* cells (TCA extracts, SC, 30 °C, Spt5-TAP background). Deletion of *SPT4* restores Spt5-P and H3K4me3, and partially restores H2Bub1, in the *mog1Δ* background. Ponceau staining was used as loading control.

A key molecular feature of elongating RNAPII is the phosphorylation of its C-terminal domain (CTD) at serine 2. Using an α-Ser2P RNAPII CTD antibody, we found that global Ser2 phosphorylation was reduced to approximately half of wild-type levels in *mog1Δ* cells (0.51 ± 0.17, n = 3; Figure 1B). Mutant cells lacking *BUR2* were used as positive control.

A hallmark of early transcription elongation is the chromatin recruitment and phosphorylation of the transcription elongation factor Spt5 (7, 11). ChIP-qPCR analysis of TAP-tagged Spt5 revealed substantially reduced chromatin association at both the 5′ and 3′ regions of *ADH1, YEF3, PYK1* and *PMA1* in *mog1Δ* compared to wild type (Figure 1C). This reduction was comparable across all regions and genes analyzed. A reduction in Spt5 occupancy could in principle reflect the lower global transcription rate of *mog1*Δ cells (mRNA synthesis rate 0.75 of wild-type (26)) rather than a specific defect in Spt5 engagement. Two observations argue against this interpretation. First, in that study RNAPII occupancy measured as Rpb3 ChIP at the same regions of *ADH1, PMA1* and *YEF3* was reduced to a smaller extent than the reduction in Spt5 occupancy reported here. Second, at the *PMA1* 5′ region Rpb3 occupancy was unchanged in *mog1*Δ cells, whereas Spt5 occupancy was reduced by more than half. Although these measurements come from independent experiments and cannot be combined into a single ratio, the comparison indicates that reduced polymerase density alone cannot account for the loss of Spt5 from transcribed chromatin. Total Spt5-TAP levels were unchanged (Figures 1D and 2E), excluding reduced protein abundance as the explanation.

Next, to determine whether the reduced chromatin association was linked to defective Spt5 modification, we assessed global Spt5 phosphorylation status; wild-type and *bur2*Δ were used as internal controls. In standard western blots (Figure 1D, top), *mog1*Δ cells showed decreased intensity of a slower-migrating band likely corresponding to phosphorylated Spt5. This effect of deleting *MOG1* was further confirmed by Phos-tag electrophoresis, since the identity of the upper band as the phosphorylated form of Spt5 has been previously validated by λ-phosphatase treatment (43). As shown in Figure 1D (bottom), phosphorylation-dependent Spt5 bands were clearly detected in wild-type extracts but were markedly diminished in *mog1*Δ and absent in *bur2*Δ cells.

Finally, we asked whether the functional relationship between Mog1 and Spt5 is accompanied by a physical association. Tandem affinity purification of Mog1 followed by LC-MS/MS recovered Spt5 as the transcription-related protein with the highest sequence coverage (47 %), with Spt4 also among the most represented (30 %), whereas neither Bur1 nor Bur2 was detected; proteins also identified in a parallel purification from an untagged strain were excluded (Table S4). In addition, silver staining of the Mog1-TAP eluate resolved two discrete species migrating above 100 kDa, both of which were excised from the gel and identified as Spt5 by mass spectrometry (Figure 1E), consistent with its predicted mass of 117 kDa. Their different mobilities are compatible with the unphosphorylated and phosphorylated forms resolved in Figure 1D, although the modification state of the slower-migrating species was not determined. Co-purification does not establish a direct contact, but it places Mog1 in the vicinity of the Spt4-Spt5 module.

Collectively, these results indicate that Mog1 is required to sustain Spt5 phosphorylation and its association with chromatin during transcription elongation, in addition to maintaining RNAPII CTD Ser2 phosphorylation levels, and that Mog1 and Spt5 are found in the same purified complexes. These molecular effects are consistent with the elongation-associated phenotype of *mog1*Δ cells.

### Spt4 accumulates in *mog1*Δ cells and its deletion suppresses *mog1*Δ phenotypes restoring Spt5 phosphorylation

Having established that Mog1 is required to sustain Spt5 C-terminal phosphorylation and its chromatin association and that they copurify, we next sought the molecular route through which Mog1 acts. Because Spt5 is essential and cannot be deleted, we turned to its non-essential binding partner Spt4, which directly associates with Spt5 and modulates elongation dynamics (6, 9, 10). We reasoned that combining *MOG1* and *SPT4* deletions would reveal whether Spt4 mediates the elongation defect caused by loss of Mog1, and analyzed their genetic interactions under basal and transcription-challenging conditions. Tetrad dissection of the *spt4Δ mog1Δ* double mutant revealed clear suppression of the growth defect associated with *mog1Δ* following deletion of *SPT4.* At 30 °C on YPD, the *spt4Δ mog1Δ* double mutant formed visibly larger colonies than the *mog1Δ* single mutant, which displayed markedly reduced growth (Figure 2A). This suppression was also evident both by growth spot assays (Figure 2B, upper panel) and by morphologic visualization (Figure 2B, lower). Thus, on YPD medium, deletion of *SPT4* restored the growth of *mog1Δ* cells to the level of *spt4*Δ cells at both 30 °C and 37 °C, and restored normal cell morphology. Interestingly, in a Spt5-TAP background, the *spt4Δ mog1Δ* double mutant displayed synthetic lethality on SC at 37 °C (Figure 2C). Thus, the genetic interaction between *mog1Δ, spt4Δ* and *SPT5-TAP* is strongly context-dependent and sensitive to stress. Together, these data demonstrate that loss of Spt4 alleviates the basal growth defects caused by Mog1 deficiency, but it depends on fully functional Spt5.

Next, we checked whether this suppression extended to conditions of transcription stress. Deletion of *SPT4* also relieved the MPA sensitivity of *mog1*Δ cells at 30 °C on SC medium (Figure 2D), and, as expected, guanine supplementation reversed all MPA phenotypes (Figure 2D and Figure S1). Thus, removing Spt4 alleviates both the basal and the elongation-stress growth defects of *mog1*Δ.

To assess whether genetic suppression correlated with the restoration of elongation-associated molecular states, we analyzed Spt5 phosphorylation together with elongation-associated chromatin marks in SC-grown cells at 30 °C using TCA extracts prepared from strains in the Spt5-TAP background. Whereas *mog1Δ* cells show reduced Spt5 phosphorylation (Figure 1), the *spt4Δ mog1Δ* double mutant displayed increased Spt5 phosphorylation relative to *mog1Δ* alone (Figure 2E). This increase was accompanied by a clear recovery of H3K4 trimethylation and a partial recovery of H2BK123 monoubiquitination, as detected within the same immunoblot panel (Figure 2E). These results indicate that deletion of *SPT4* restores a coordinated set of elongation-associated phosphorylation and chromatin states that are disrupted in *mog1Δ cells.* We note that this analysis was performed in the Spt5-TAP background at 30 °C, a condition in which the *spt4*Δ *mog1*Δ double mutant grows normally; the synthetic lethality of this combination in the same background at 37 °C (Figure 2C) indicates that the suppression is conditional and sensitive to the functional status of Spt5.

### Spt4 accumulates in *mog1*Δ cells without changes in its subnuclear localization

To gain further mechanistic insight into how Spt4 suppresses the *mog1*Δ defects, we examined Spt4 protein levels in the absence of Mog1. Western blot analysis using an anti-GFP antibody in a strain expressing genomically tagged Spt4-GFP revealed that Spt4-GFP protein levels were consistently higher in *mog1*Δ cells than in wild-type cells (Figure 3A), indicating that Mog1 restrains Spt4 abundance. Since Mog1 is a factor that participates in protein transport (21, 22), we tested whether the elevated Spt4 signal reflected altered localization besides its increased protein levels, and examined Spt4-GFP by fluorescence microscopy. Initial inspection of the GFP signal revealed a heterogeneous intranuclear distribution, with a region of higher fluorescence intensity adjacent to a region of lower intensity within the same nucleus (Figure 3B). This pattern raised the possibility that Spt4 might preferentially accumulate in, or conversely be excluded from, a specific subnuclear compartment such as the nucleolus. To directly test this, strains co-expressing Spt4-GFP and Net1-mCherry, a protein associated with rDNA and involved in nucleolar organisation, were generated to assess Spt4 distribution relative to the nucleolus. To facilitate interpretation of the GFP signal, two additional nuclear markers were introduced via centromeric plasmids: Sec63-mCherry, an endoplasmic reticulum translocon component that outlines the nuclear envelope and defines the nuclear boundary, and Pus1-mCherry, a pseudouridine synthase displaying diffuse nucleoplasmic distribution that serves as a reference for the nuclear interior. Together, these markers allowed unambiguous discrimination between nucleoplasmic, nucleolar, and cytoplasmic localisation of Spt4-GFP. Critically, this nucleoplasmic distribution of Spt4-GFP was unchanged in *mog1*Δ cells relative to wild type (Figure 3C), indicating that the increased Spt4 abundance in the absence of Mog1 reflects a functional rather than a spatial effect.

**Figure 3.**
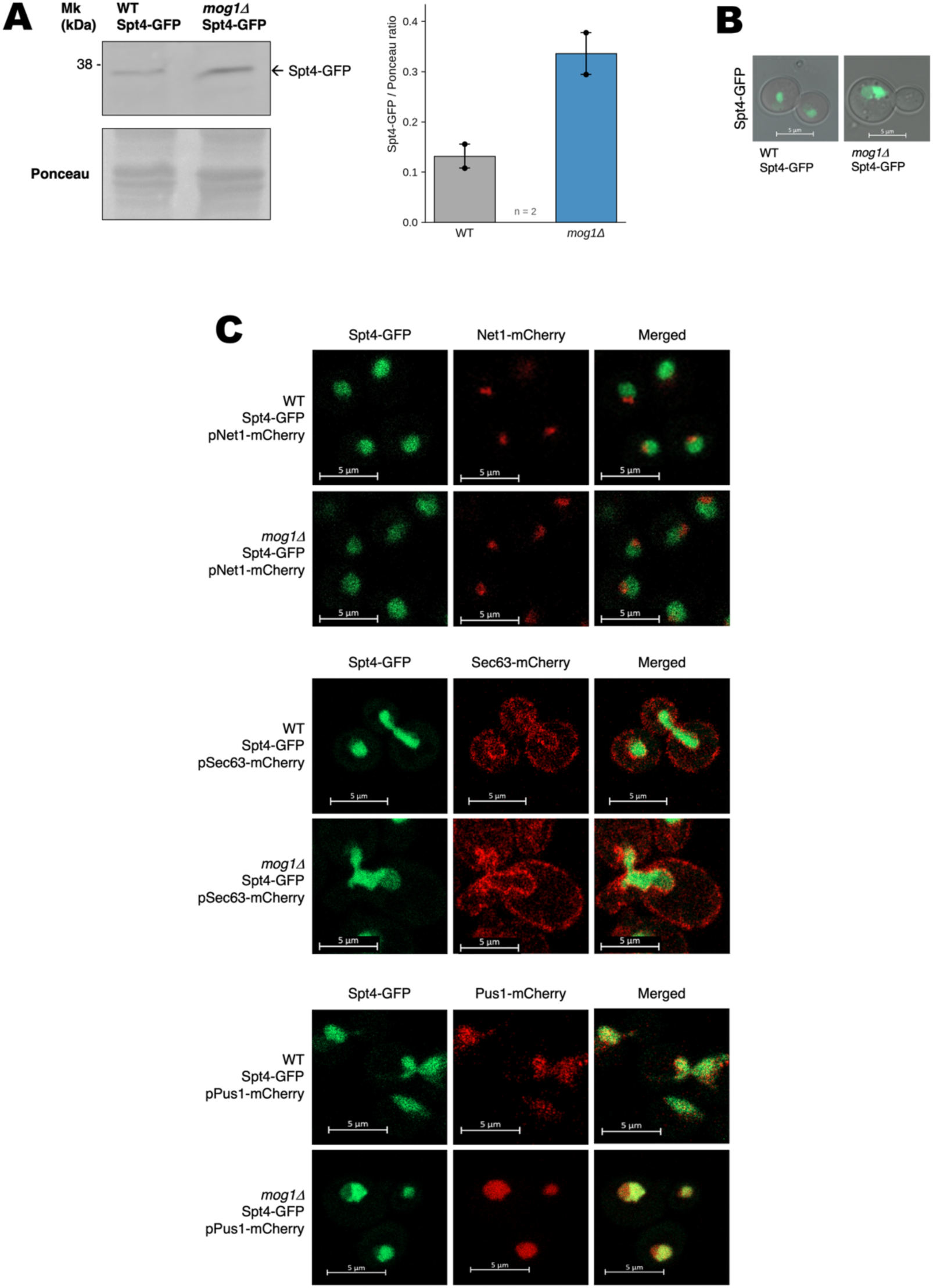
Spt4 accumulates in *mog1*Δ cells without a change in its nucleoplasmic localisation. (A) Immunoblot (anti-GFP) of genomically tagged Spt4-GFP in wild-type and *mog1*Δ cells. Quantification of Spt4-GFP signal intensity normalised to total protein (Ponceau staining). Data represent the mean ± SEM of two independent biological replicates (n=2). (B) Representative fluorescence microscopy images of Spt4-GFP in wild-type and *mog1*Δ cells, merged with brightfield images to visualise cell morphology. (C) Representative fluorescence microscopy images of Spt4-GFP relative to nuclear markers (Net1-mCherry, nucleolus; Sec63-mCherry, nuclear envelope; Pus1-mCherry, nucleoplasm) in wild-type and *mog1*Δ cells. Individual fluorescence channels and merged images are shown. The nucleoplasmic distribution of Spt4-GFP is unchanged in *mog1*Δ, indicating a functional rather than a spatial effect. Scale bars, 5 µm (B, C).

### *mog1*Δ interacts genetically with the BUR pathway mutants without altering Bur1 levels or phosphorylation

Together, our results identify Spt4 as the mediator of the *mog1*Δ elongation defect, in the absence of Mog1, Spt4 accumulates and its removal restores Spt5 phosphorylation and the downstream H2Bub1 and H3K4me3 marks. This suggests that Mog1 helps to maintain Spt4 within a range that permits Spt5 modification. Thus, the logical next step was to examine the functional interaction between Mog1 and the kinase pathway responsible for Spt5 phosphorylation.

The BUR complex, comprising the cyclin-dependent kinase Bur1 and its cyclin partner Bur2, phosphorylates the C-terminal repeats of Spt5 during early elongation (5, 11). We combined *MOG1* deletion with either *BUR2* deletion or the conditional *bur1-80* mutant (*BUR1* is essential) (11) and assessed growth under different conditions (Figure 4A-C). Under optimal temperatures (27 °C and 30 °C), deletion of *MOG1* slightly suppressed the severe growth defect of *bur2*Δ cells, whereas at 37 °C the *mog1*Δ *bur2*Δ double mutant was synthetically lethal (Figure 4A). In addition, a non-additive reduction of Ser2-P and Spt5-P is observed in the *mog1*Δ *bur2*Δ double mutant (Figure S2).

**Figure 4.**
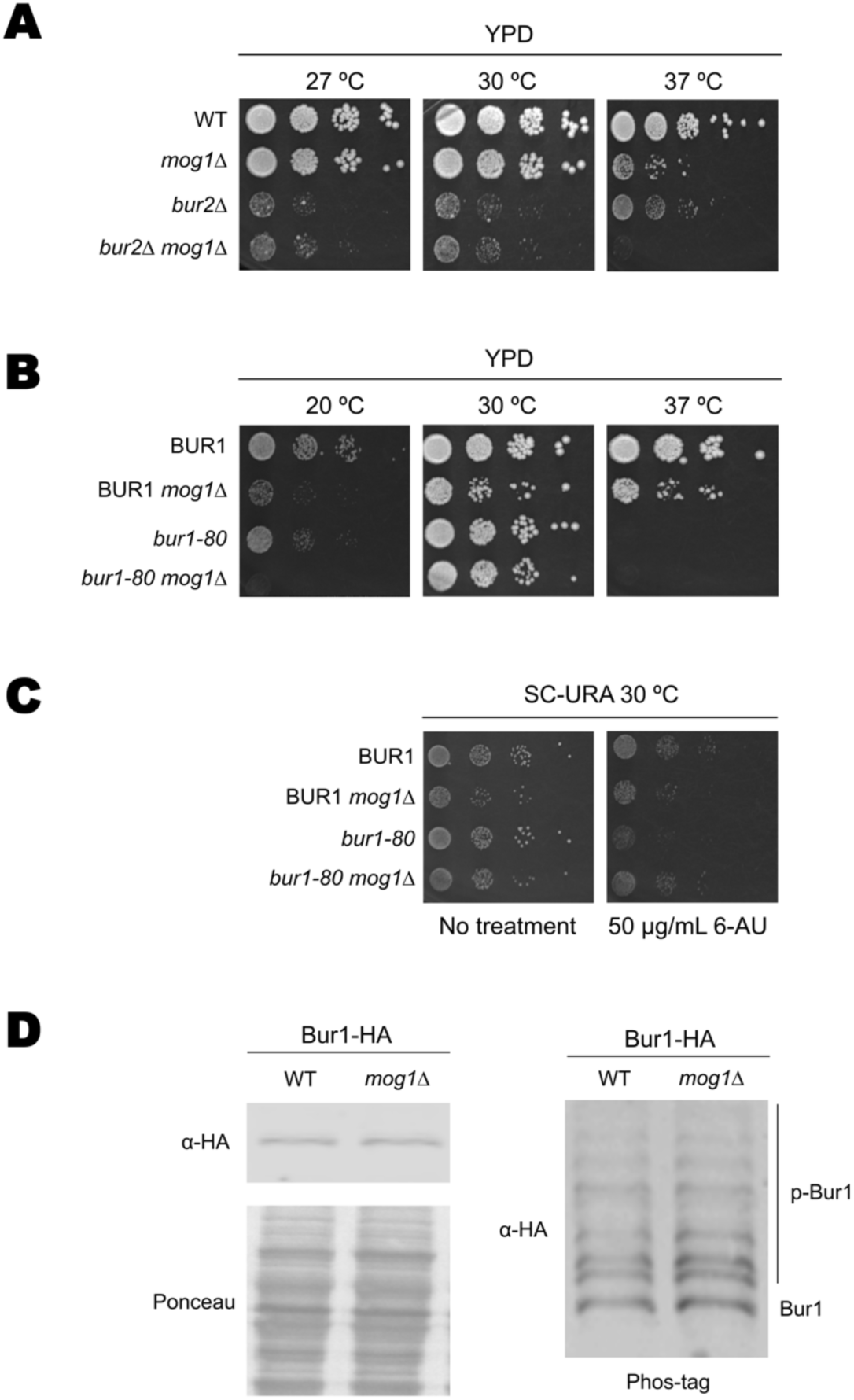
Mog1 interacts genetically with the BUR pathway without altering Bur1 abundance or activation. (A) Genetic interactions of *MOG1* with *BUR2*. Ten-fold serial dilution spot assays of *WT*, *mog1Δ*, *bur2Δ* and *bur2Δmog1Δ* strains grown on YPD plates at 27 °C, 30 °C and 37 °C (B) Spot assays of bur1-80 and bur1-80 *mog1*Δ at 20 C°, 30 C° and 37 °C. Deletion of *MOG1* restores growth at 30 °C; at 20 °C the double mutant is synthetically lethal, and at 37 °C both bur1-80 and the double mutant are inviable. (C) Spot assays on 6-azauracil (6-AU). Deletion of *MOG1* alleviates the 6-AU sensitivity of bur1-80, with bur1-80 *mog1*Δ growing better than either single mutant. (D) Bur1-HA analysed by standard SDS-PAGE and Phos-tag gel electrophoresis in wild-type and *mog1*Δ cells. Ponceau staining was used as a loading control. Bur1 abundance and overall phosphorylation are comparable, indicating that Mog1 does not affect Bur1 stability or activation.

At 30 °C, bur1-80 cells were not appreciably impaired, whereas *mog1*Δ cells showed their characteristic growth defect; the *bur1-80 mog1Δ* double mutant grew better than the *mog1Δ* single mutant (Figure 4B, middle panel), indicating that reducing Bur1 function partially suppresses the loss of Mog1. Because *bur1-80* cells proliferate slowly at both low and high temperatures, we assessed interactions under restrictive conditions, at 20 °C the *bur1-80 mog1Δ* double mutant displayed synthetic lethality (Figure 4B, left panel), whereas at 37 °C both *bur1-80* and the double mutant were non-viable (Figure 4B, right panel). Notably, the single *mog1*Δ strain also showed reduced growth at 20 °C. Furthermore, *bur1-80* cells are sensitive to 6-azauracil (6-AU), a compound that also impairs transcription elongation by limiting nucleotide availability, thereby exacerbating defects in RNA polymerase progression (11, 42). Remarkably, deletion of *MOG1* alleviated this sensitivity, with *bur1-80 mog1Δ* displaying improved growth compared with either single mutant (Figure 4C).

To evaluate whether these genetic effects could be explained by altered Bur1 stability or activation, we analyzed Bur1-HA levels using conventional SDS-PAGE and Phos-tag gel electrophoresis, which separates phosphorylated from non-phosphorylated species (31). Bur1 is activated through phosphorylation by the cyclin-dependent kinase Cak1 (44). As shown in Figure 4D, Bur1 protein abundance and overall phosphorylation were comparable between wild-type and *mog1*Δ cells. These observations indicate that Mog1 does not affect Bur1 stability or activation under the conditions tested.

Together, these genetic interactions define a functional relationship between Mog1 and the BUR pathway and constrain its nature. Bur1 abundance and overall phosphorylation were unchanged in *mog1*Δ cells (Figure 4D), arguing against an effect on kinase stability or on Cak1-dependent activation. Moreover, if Mog1 provided a positive input into BUR activity, loss of Mog1 would be expected to aggravate bur mutants. Instead, under permissive conditions the two perturbations partially compensated one another: deletion of *MOG1* improved the growth of *bur2*Δ cells at 27 °C and 30 °C (Figure 4A), the *bur1-80* allele improved the growth of *mog1*Δ cells at 30 °C (Figure 4B), and *bur1-80 mog1*Δ cells grew better than either single mutant on 6-AU (Figure 4C). At temperature extremes, by contrast, the double mutants were inviable (Figure 4A, B), indicating that both activities become jointly required when elongation is further challenged. Mutual suppression under permissive conditions, together with unaltered Bur1 levels and activation, indicates that Mog1 does not act by promoting BUR kinase activity. Taken with the Spt4 dependence of the *mog1Δ* phenotypes, these data place the action of Mog1 on the substrate side of the reaction, the Spt4-Spt5 complex, rather than on the kinase.

### Mog1 becomes critical for fitness when early transcription elongation is compromised

Because Mog1 sustains Spt5 phosphorylation by acting on the Spt4-Spt5 substrate rather than on the BUR kinase, we asked whether its contribution to elongation extends to other steps of the elongation cycle. We therefore examined genetic interactions with two additional, functionally distinct elongation factors, Ctk1, the kinase that reinforces RNAPII Ser2 phosphorylation during late elongation (45), and Dst1 (TFIIS), which promotes RNAPII recovery from backtracking (46).

Tetrad dissection (Figure 5A, left) revealed markedly impaired growth in the *ctk1*Δ *mog1*Δ double mutant, which formed extremely small colonies at 30 °C. After prolonged incubation, these colonies exhibited an abrupt shift toward rapid growth, suggesting a reversion event. Owing to this heterogeneous behaviour, spot assays could not be reliably performed on this strain. In contrast, the interaction with *DST1* was milder but consistent. The *dst1*Δ *mog1*Δ double mutant showed reduced colony size in tetrad analysis (Figure 5A, right) and exhibited slightly impaired growth in spot assays on YPD at both 30 °C and 37 °C (Figure 5B). This indicates a modest reduction in fitness when elongation-recovery pathways are compromised.

**Figure 5.**
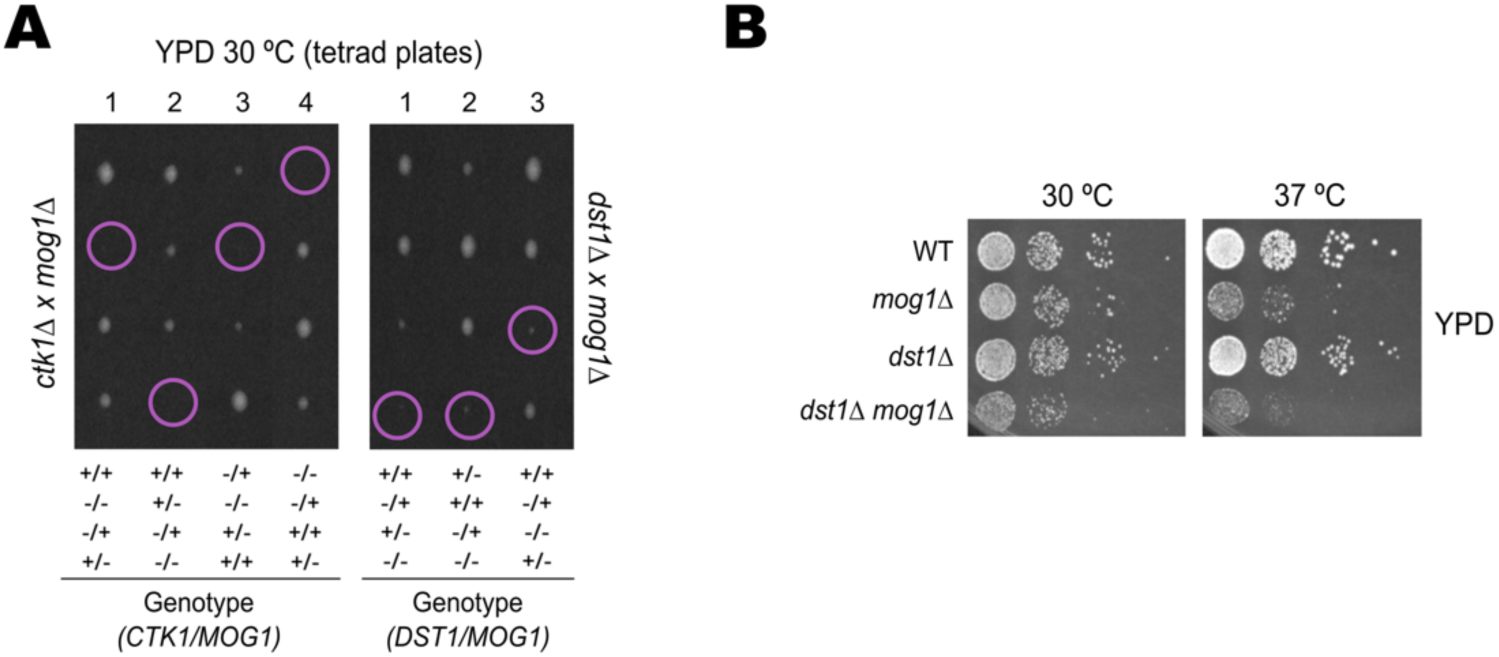
Genetic interactions of *MOG1* with *CTK1* and *DST1*. (A) Tetrad dissection of diploids resulting from crosses *ctk1Δ* × *mog1*Δ (left) and *dst1Δ* × *mog1Δ* (right). Each vertical column corresponds to a single tetrad, and each horizontal row to an individual meiotic product. Colonies were grown on YPD at 30 °C for 3 days. Genotypes are indicated below using “+” for wild type and “–” for gene deletion as follows: +/+ = corresponding double wild-type (*CTK1 MOG1* or *DST1 MOG1*), –/+ = *ctk1Δ MOG1* or *dst1Δ MOG1*, +/– *= CTK1 mog1Δ* or *DST1 mog1Δ*, –/– *= ctk1Δ mog1Δ* or *dst1Δ mog1Δ*. (B) Tenfold serial dilution spot assays of WT, d*st1Δ*, *mog1Δ* and *dst1Δ mog1Δ* strains. Cells were grown to exponential phase and spotted on YPD plates incubated at 30 °C or 37 °C for 2–3 days before imaging.

Together, these results indicate that Mog1 becomes critical for cellular fitness when transcription elongation is challenged, with strong synthetic defects when RNAPII CTD Ser2 reinforcement is impaired (*ctk1*Δ) and subtler effects when polymerase backtracking resolution is affected (*dst1*Δ). The graded severity, severe with Ctk1, mild with Dst1, suggests that Mog1 acts most closely with the Spt5/Ser2-phosphorylation arm of elongation rather than with backtracking recovery.

### Preventing H3K4me3 demethylation is not sufficient to compensate the loss of Mog1

The restoration of H2B monoubiquitination and H3K4 trimethylation in the *mog1*Δ *spt4*Δ double mutant (Figure 2E) indicated that these chromatin states are downstream consequences of the elongation defect rather than independent effects of Mog1 loss. This left open whether the loss of H3K4me3 itself contributes to the *mog1*Δ phenotypes. To address this, we uncoupled the mark from the elongation defect by deleting *JHD2*, which encodes the principal H3K4me3 demethylase in *S. cerevisiae* (47), and asked whether preventing H3K4me3 removal is sufficient to compensate for the loss of Mog1. Western blot analysis confirmed that this manipulation fully restored the mark. As expected, *jhd2*Δ cells displayed H3K4me3 levels comparable to wild-type, and additional deletion of *MOG1* did not reduce H3K4me3 in the *jhd2*Δ background (Figure 6A), in contrast to the ∼50% decrease observed in *mog1*Δ single-mutant cells (Figure 2E and Figure 6A). Thus, in the *jhd2*Δ *mog1*Δ double mutant global H3K4me3 is restored to *jhd2*Δ levels, i.e. at or above wild-type.

**Figure 6.**
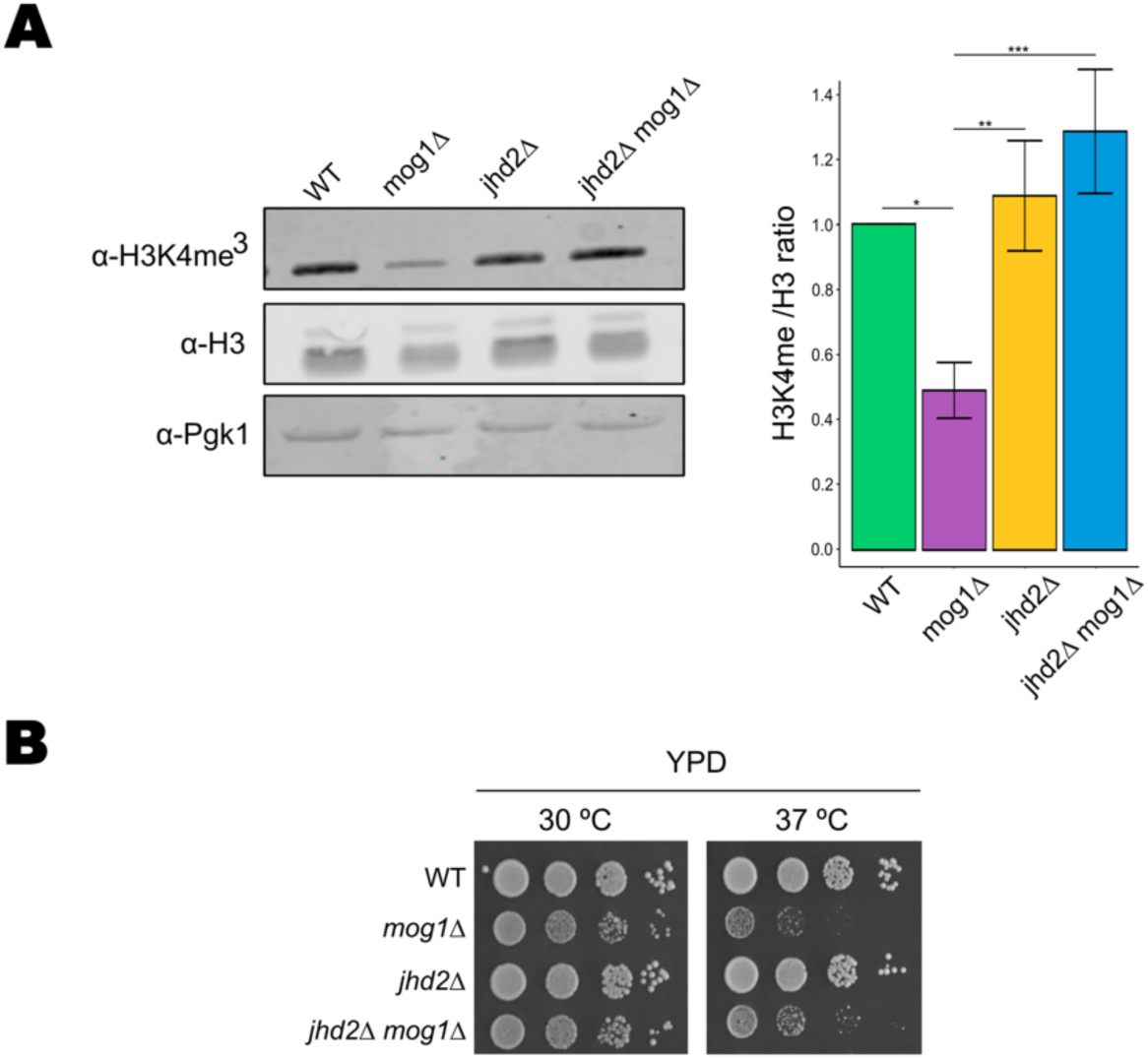
Restoring H3K4me3 by deletion of the demethylase *JHD2* does not suppress the growth defects of *mog1*Δ. (A) Western blot analysis of global H3K4me3 levels in WT, *mog1Δ, jhd2Δ* and *jhd2Δmog1Δ* strains. Acid-extracted histones were probed with anti-H3K4me3, and total histone H3 and Pgk1 were used as loading controls. H3K4me3 and H3 signals were first normalized to Pgk1 within each blot, and the H3K4me3/H3 ratio was then calculated for each biological replicate. The right panel shows quantification of H3K4me3/H3 ratios, normalized to WT, with standard errors propagated using first-order Taylor expansion (delta method). Bar charts represent mean ± SEM from n = 4 biological replicates. Statistical differences were evaluated using one-way ANOVA followed by Tukey’s post hoc test (*P = 0.01–0.05; **P < 0.01; ***P < 0.001). (B) Tenfold serial dilution spot assays of WT, mog1Δ, jhd2Δ and jhd2Δmog1Δ strains. Cells were grown to exponential phase and plated onto YPD medium. Plates were incubated for 2 days at 30 °C or 37 °C. At 37 °C both mog1Δ and jhd2Δmog1Δ give rise to colonies of variable size and frequency between experiments, and these spots were not scored quantitatively; in no experiment did deletion of JHD2 restore jhd2Δ- or wild-type-like growth to mog1Δ cells.

Despite this full restoration of the mark, growth was not restored. Serial-dilution assays showed that *jhd2*Δ *mog1*Δ cells grew essentially as *mog1*Δ cells and clearly worse than *jhd2*Δ or wild-type cells, both at 30 °C and at 37 °C (Figure 6B). At 30 °C the double mutant retained the reduced, heterogeneous colony growth characteristic of *mog1*Δ at the higher dilutions, and at 37 °C it remained strongly impaired, forming only sparse microcolonies in the first dilutions, whereas *jhd2*Δ was indistinguishable from wild-type. We note that at 37 °C both *mog1*Δ and *jhd2*Δ *mog1*Δ give rise to colonies of variable size and frequency between experiments, so these spots were not scored quantitatively, in no experiment did deletion of *JHD2* restore *jhd2*Δ- or wild-type-like growth to *mog1*Δ cells.

Together, these results show that the reduction of H3K4me3 in *mog1*Δ cells can be fully prevented by deletion of *JHD2*, but that preventing it is not sufficient to relieve the growth defect caused by the loss of Mog1.

### Mog1 mutants differentially affect its role in transcription elongation

We next mapped two residues of independent biological interest onto the crystal structure of yeast Mog1 (PDB 1EQ6). First, we studied the Mog1-W145 residue (Figure 7A), a surface-exposed tryptophan (side-chain relative accessibility ∼58% on the 1EQ6 crystal structure), which falls within the region of Mog1 (residues 114–154) identified as the Shg1-binding domain in our previous work (26, 28). Second, we studied E65 (Figure 7A), whose substitution (E65K) impairs Ran binding (21). Notably, W145 and the E65-proximal surface lie on opposite faces of the Mog1 fold (Cα–Cα separation >20 Å on the 1EQ6 crystal structure), allowing their functional contributions to be tested independently.

**Figure 7.**
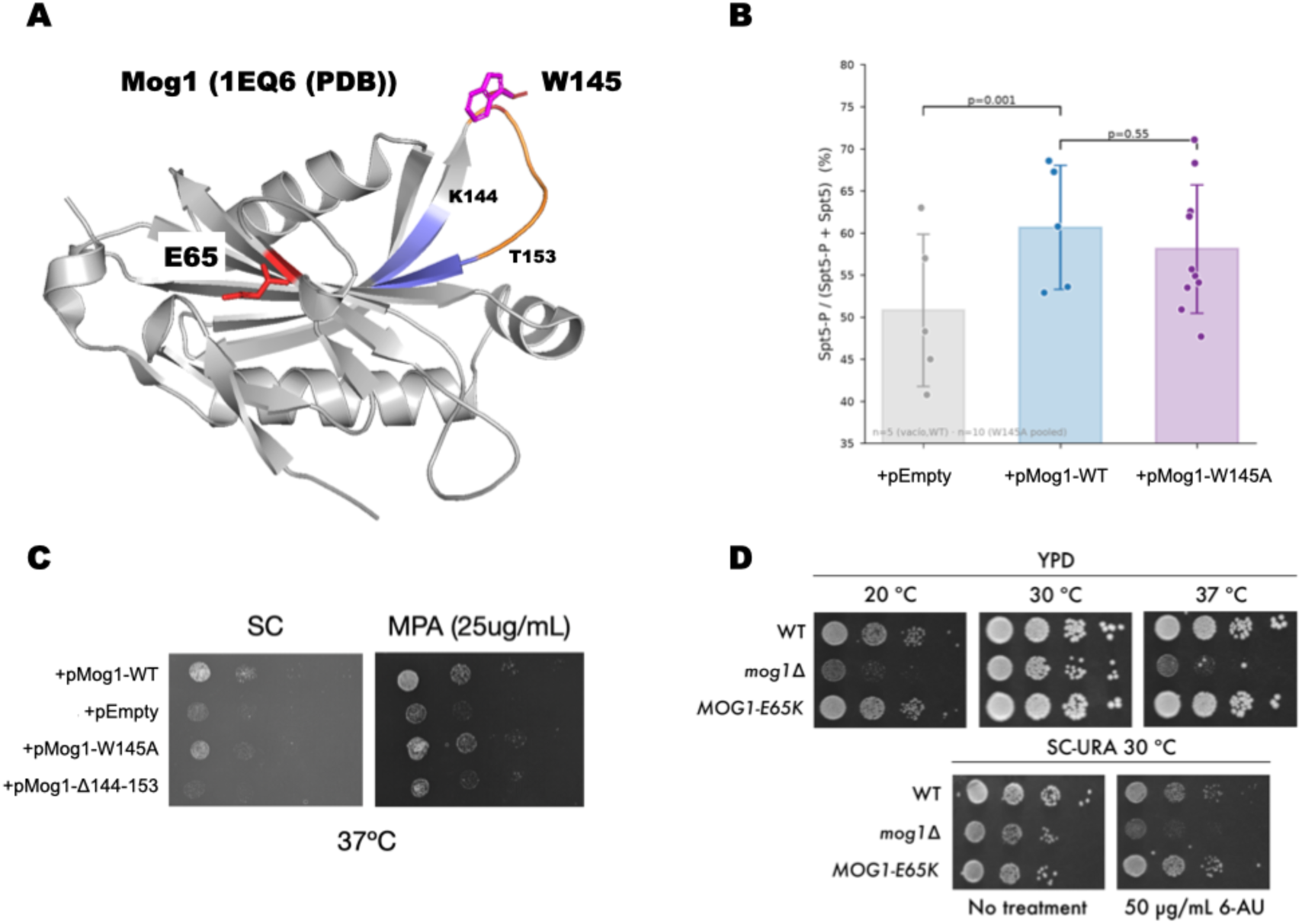
Point substitutions on either face of Mog1 leave its elongation function intact. (A) Structural mapping of the mutated residues onto the yeast Mog1 crystal structure (PDB 1EQ6, 1.9 Å resolution). The K144–T153 loop is shown in orange; flanking β-strands (β138–140 and β152–156) in blue. The W145 side chain (magenta) lies within the loop, surface-exposed (relative accessibility ∼58 %). The E65 side chain (red) marks the Ran-binding face on the opposite side of the fold (Cα–Cα separation >20 Å). Left: global view of the Mog1 fold. Right: close-up of the loop region showing K144, W145 and T153. (B) Spt5 phosphorylation in *mog1*Δ cells expressing empty vector, Mog1-WT or Mog1-W145A from a centromeric plasmid. Bar charts show the Spt5-P/(Spt5-P + Spt5) ratio (%). Data represent mean ± SD; individual data points are shown. mog1Δ + empty vector reduces the ratio relative to Mog1-WT (p = 0.001, paired t-test, n = 5); W145A is not significantly different from Mog1-WT (p = 0.55, n.s.). (C) Ten-fold serial dilution spot assays of mog1Δ cells expressing pMog1-WT, empty vector, pMog1-W145A or pMog1-ΔK144-T153 on SC and SC + MPA (25 µg/mL) at 37 °C. W145A complements *mog1*Δ growth; deletion of the entire K144–T153 loop does not. (D) Ten-fold serial dilution spot assays of wild-type, *mog1*Δ and genomic *mog1-E65K* strains on YPD at 20 °C, 30 °C and 37 °C, and on SC-URA at 30 °C with or without 50 µg/mL 6-azauracil (6-AU). *mog1-E65K* grows like wild-type under all conditions tested.

To probe these two Mog1 faces functionally, we assessed the corresponding mutants across independent functional read-outs. As a first output, we measured Spt5 phosphorylation. *mog1*Δ cells expressing an empty vector reduced the Spt5-P/Spt5 ratio to 83.4 ± 5.7% of Mog1-WT levels, confirming that Mog1 is required for full Spt5 phosphorylation and that the system recapitulates the *mog1*Δ phenotype (Figure 7B). The Mog1-W145A allele sustained Spt5 phosphorylation at wild-type levels (95.8 ± 5.5%, n.s.). Consistent with this, Mog1-W145A cells showed no growth phenotype on MPA and complements *MOG1* deletion in minimal medium at high temperature (Figure 7C). Thus W145, despite lying in the Shg1-interaction region, is dispensable for Mog1’s function in Spt5 phosphorylation.

Mapping the mutated residues onto the Mog1 crystal structure (PDB 1EQ6) shows that W145 lies within a solvent-exposed loop (residues 144–153) that connects two β-strands of the fold (β138–140 and β152–156; Figure 7A). Whereas the Mog1-W145A substitution removes a single surface side chain, deletion of the entire K144-T153 loop (Mog1-ΔK144-T153) removes a ten-residue structural connector between these β-strands. In contrast to the Mog1-W145A mutant, Mog1-ΔK144-T153 does not restore *mog1*Δ growth on SC or MPA (Figure 7C).

Lastly, extending our previous conclusion that the chromatin-related function of Mog1 does not require Ran binding (26), we introduced the E65K mutation directly into the genomic *MOG1* locus by CRISPR-Cas9, avoiding the overexpression inherent to plasmid-based complementation. Unlike *mog1Δ,* and similar to Mog1-W145A, the genomic Mog1-E65K mutant grew like wild-type under all conditions tested, including 20 °C, 30 °C, 37 °C and 6-AU (Figure 7D). In the plasmid-based Spt5-P assay, Mog1-E65K was likewise comparable to Mog1-WT (Figure S3). Taken together, neither the Mog1-W145A nor the Mog1-E65K substitution experimentally abolished the role of Mog1 in Spt5 phosphorylation or elongation-linked growth.

## DISCUSSION

Transcription elongation requires the coordinated action of RNA polymerase II (RNAPII), elongation factors and the chromatin modifications that accompany them. Mog1 was characterised as a Ran-binding protein required for nuclear protein import, and our previous work connected it to H2B monoubiquitination and H3K4 trimethylation without identifying the step at which it acts (26). The results presented here identify that step. Mog1 is required to sustain phosphorylation of the Spt5 C-terminal repeat (CTR) and, with it, RNAPII CTD Ser2 phosphorylation and Spt5 association with transcribed chromatin; the defects of *mog1*Δ cells require Spt4; and the chromatin changes previously described are a consequence of the elongation defect rather than its cause. Two independent lines of evidence place the action of Mog1 on the Spt4-Spt5 substrate rather than on the kinase that modifies it. Genetically, Bur1 abundance and overall phosphorylation are unchanged in *mog1*Δ cells, and reducing BUR activity does not aggravate the loss of Mog1 but partially compensates it under permissive conditions, which is the opposite of what a positive input into kinase activity would predict. Biochemically, Spt5 is not merely detected in Mog1 preparations but is abundant enough to be visible by silver staining, and Spt4 is likewise among the best-represented partners, whereas neither Bur1 nor Bur2 is recovered at all. Co-purification does not establish a direct contact, and the association may be indirect or transient within elongation complexes, but the contrast is internal to the same preparations and therefore independent of how the experiment is normalised. Defining a physical interface will require approaches capable of detecting transient contacts, such as cross-linking mass spectrometry or in vitro reconstitution.

What Spt4 contributes to this arrangement is constrained by the epistasis. Deletion of *SPT4* in the *mog1*Δ background restores Spt5 phosphorylation, the associated chromatin marks, and normal growth and cell morphology, establishing that Spt4 is required for the *mog1*Δ phenotypes to manifest. Spt4 levels were also higher in *mog1*Δ cells, with no detectable change in nucleoplasmic distribution; whether this excess contributes to the defect or follows from it has not been tested. We propose that what limits Spt5 CTR phosphorylation is not the absolute level of kinase activity but the balance between kinase output and the accessibility of the CTR within the Spt4-Spt5 complex, with Mog1 acting on the substrate side of that balance. The model accounts for the full set of genetic interactions: lowering either side alone is deleterious, whereas lowering both partially restores the balance, which is why *spt4*Δ rescues *mog1*Δ, why reduced BUR activity does so under permissive conditions, and why the same combinations become inviable when temperature places an additional demand on elongation. It is also consistent with the established role of Spt4 in shaping Spt5 conformation within the elongation complex (6) and with work placing the Spt4-Spt5 axis at the nucleosomal boundary that governs early elongation competence (48).

The output of this system is sensitive to the integrity of Spt5 itself. In the Spt5-TAP background the suppression of *mog1*Δ by *spt4*Δ is lost at 37 °C, where the double mutant becomes inviable, even though it grows normally at 30 °C in the same background, the condition under which the molecular analysis was performed. A modest perturbation introduced by C-terminal tagging is therefore enough to change the sign of the interaction, which supports the view that Mog1 operates within a finely tuned system and, equally, sets a limit on how far the suppression should be generalised.

The consequences of reduced Spt5 CTR phosphorylation extend beyond elongation itself. CTR phosphorylation contributes to the association of Paf1C with elongating RNAPII, and dephosphorylation of the CTR by Glc7/PP1 dislodges Paf1C near the cleavage and polyadenylation site (20); because H2Bub1 and H3K4me3 lie downstream of Paf1C, a defect in CTR phosphorylation offers a direct route to the chromatin phenotype we reported previously (26). The requirement for the CTR in Paf1C recruitment has nonetheless been debated, since Bur1 and Kin28 can promote Paf1C association independently of it (16). Our data do not resolve that question, but the coordinate loss and recovery of Spt5-P together with H2Bub1 and H3K4me3 is most readily accommodated by a CTR-phosphorylation-dependent contribution to Paf1C function. Whether the elongation defect of *mog1*Δ cells also affects Glc7-dependent Paf1C dissociation and transcription termination remains open.

That the chromatin marks are downstream is established directly rather than inferred. Deletion of *JHD2* fully restored H3K4me3 in *mog1*Δ cells without restoring growth, whereas deletion of *SPT4* restored both the mark and growth together with Spt5 phosphorylation. Because *jhd2*Δ restores H3K4me3 without restoring the upstream H2Bub1 signal, whereas *spt4*Δ restores both together with elongation competence, the comparison separates the mark from the process and indicates that it is the elongation defect, and not the chromatin mark per se, that limits the growth of *mog1*Δ cells. The synthetic sickness of *mog1*Δ *set1*Δ cells reported previously (26) is not in conflict with this conclusion, since deletion of *SET1* abolishes all H3K4 methylation states whereas deletion of *JHD2* restores only the trimethylated form. The chromatin defect is moreover selective: H3K79me3, H3K36me3 and H3K4me2 levels are unaffected while H2Bub1 and H3K4me3 are reduced (26), a pattern expected from a defect in the Spt5-CTR/Paf1C axis rather than from a general disruption of chromatin.

The genetic interactions with *CTK1* and *DST1* delineate the window in which Mog1 operates. Ctk1 reinforces RNAPII CTD Ser2 phosphorylation during late elongation, downstream of and partially independent of Bur1. That *mog1*Δ and *ctk1*Δ are each viable while the double mutant forms only minute colonies that acquire suppressors on prolonged incubation indicates that the two activities provide partially redundant input into Ser2-P across the gene body: in *mog1*Δ cells the reduced early Ser2-P is buffered by late-stage Ctk1 activity, and removing Ctk1 eliminates that compensation. This places Mog1 in the early elongation window, upstream of or parallel to Ctk1-dependent reinforcement, consistent with the reduced but not abolished Ser2-P of *mog1*Δ single mutants. The milder interaction with *dst1*Δ is complementary: Dst1/TFIIS promotes recovery of backtracked RNAPII, a function that becomes more important when elongation efficiency is already compromised. The graded severity, severe with Ctk1, mild with Dst1, places Mog1 closer to the Spt5/Ser2-phosphorylation arm of elongation than to backtracking recovery.

The transcriptional function of Mog1 does not depend on any single surface of the protein tested here. Neither a substitution in a solvent-exposed surface loop (W145A) nor one on the Ran-binding face (E65K) abolished Spt5 phosphorylation or elongation-linked growth, whereas complete loss of Mog1 did. Mapping these residues onto the yeast Mog1 crystal structure (PDB 1EQ6) rationalises the contrast within the W145 region. W145 is a surface-exposed residue whose individual substitution is tolerated, whereas the K144-T153 segment forms a surface loop connecting two β-strands, so that deleting it removes an architectural connector and produces a growth defect on SC and MPA. The W145 result also bears on the COMPASS connection. W145 lies within the region of Mog1 identified as the Shg1-binding domain by two-hybrid screening (26), and its substitution leaves the elongation function intact, indicating that this contact is not required for the function described here. Finally, introducing E65K at the endogenous locus extends our earlier conclusion that the chromatin-related function of Mog1 is independent of Ran binding (26) to the elongation read-outs, while avoiding the overexpression inherent to plasmid-based complementation. We note that the Ran-binding defect of this substitution was established biochemically (21) and was not re-assessed here, and that the growth defect at 37 °C reported for *MOG1-E65K* in that work was not observed for the genomic allele.

These findings have a counterpart outside yeast. The idea that Spt5 CTR phosphorylation is set not only by kinase activity but also by factors controlling access to the substrate has emerged recently in metazoans, where LEDGF/p75 prevents phosphorylation of the SPT5 phosphorylation-site-rich domain and thereby promotes promoter-proximal pausing (29), and where phosphorylation of distinct SPT5 regions by CDK9 coordinately regulates pause release, elongation speed and termination (30). Within the broader parallel between the yeast Bur1-Ctk1 pair and the metazoan CDK9-CDK12 axis (3, 19), Mog1 represents the opposite sign of the LEDGF/p75 arrangement: a non-kinase factor that permits, rather than prevents, modification of the Spt5 CTR. Whether the human orthologue RANGRF retains such a role is untested, and the functional pleiotropy of Mog1 in mammalian systems (49) suggests that its contribution there may be correspondingly more complex.

Several limitations should be noted. Our readouts of elongation are global and measured at steady state: we did not determine here RNAPII elongation rate or processivity directly. However, in our previous work (26), we measured by GRO analyses that transcription synthesis is affected upon Mog1 loss. In addition, the contribution of Mog1 to elongation is inferred from Spt5 phosphorylation and occupancy, CTD Ser2 phosphorylation, co-transcriptional histone marks and sensitivity to nucleotide depletion. The epistasis establishes that Spt4 is required for the *mog1*Δ phenotypes but not that its elevated level is their proximal cause and co-purification does not demonstrate a direct contact between Mog1 and Spt4-Spt5.

Together, these results identify Mog1 as a modulatory node of early transcription elongation. By sustaining Spt5 CTR phosphorylation through an action on the Spt4-Spt5 substrate, Mog1 supports the elongation state that licenses Paf1C-dependent chromatin modification, which provides a mechanistic framework for the H2Bub1 and H3K4me3 defects we described previously (26). More generally, the work defines a constraint on Spt5 modification that is imposed by Spt4 and relieved by Mog1, and so extends the functional repertoire of a conserved Ran cofactor beyond nuclear transport.

## Acknowledgment

We thank all members of the Gene Expression laboratory for discussion and support, and Dr Queralt for critical feedback on the manuscript. We also thank Sebastián Chávez and the members of his laboratory at the Instituto de Biomedicina de Sevilla (IBiS, CSIC/Universidad de Sevilla/Hospital Universitario Virgen del Rocío), and in particular Carmen Ruger, for providing some of the mycophenolic acid plates used in this study.

## Funding

Ministerio de Ciencia, Innovación y Universidades/Agencia Estatal de Investigación, Spain [PID2024-162329NB-I00, PID2021-127734NB-I00 and PGC2018-099872-B-I00 to S.R.-N.], funded by MICIU/AEI/10.13039/501100011033 and by the European Union (FEDER/ERDF). Funding for open access charge: Ministerio de Ciencia, Innovación y Universidades/Agencia Estatal de Investigación [PID2024-162329NB-I00]

## AUTHOR CONTRIBUTIONS

J.S.-Q.: Conceptualization, Data curation, Formal analysis, Investigation, Methodology, Validation, Visualization, Writing; original draft, Writing; review and editing. C.N.-C.: Conceptualization, Formal analysis, Investigation, Methodology, Validation, Visualization, Writing; original draft, Writing; review and editing. P.M.-S.: Investigation, Methodology, Validation. S.R.-N.: Conceptualization, Funding acquisition, Project administration, Resources, Supervision, Writing; review and editing. All authors read and approved the final version of the manuscript.

## CONFLICT OF INTEREST STATEMENT

None declared

## DATA AVAILABILITY

Reviewer access details

Log in to the PRIDE website using the following details:

**Project accession:** PXD086926

**Token:** xkJvAHUaBNye

All data supporting the findings of this study are available within the article and its Supplementary Data.

## SUPPLEMENTARY FIGURES

**Figure S1.**
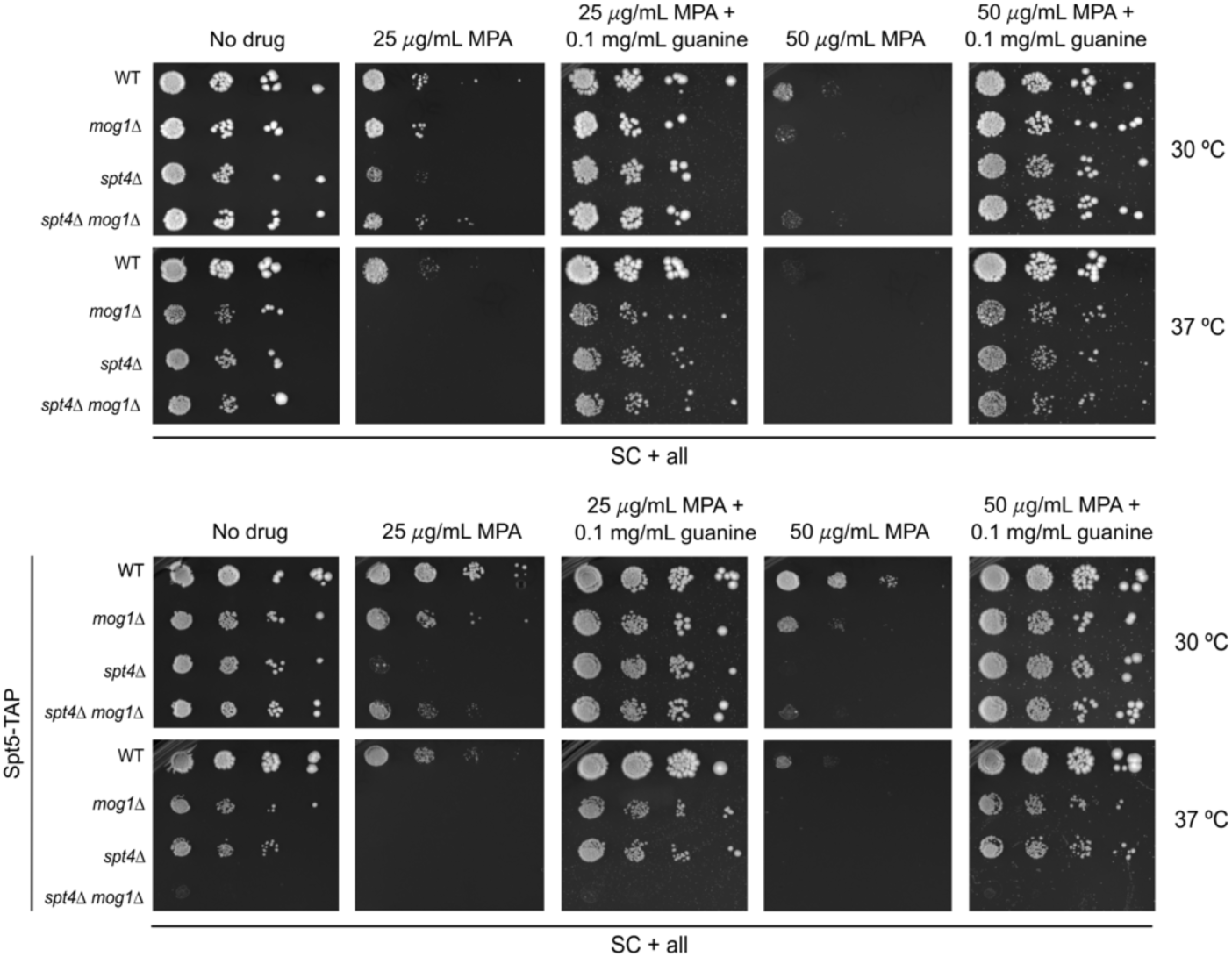
Guanine supplementation reverses the MPA sensitivity of mog1Δ cells in both Spt5-TAP and untagged backgrounds. Tenfold serial dilution spot assays of wild-type, *mog1*Δ, *spt4*Δ and *spt4*Δ *mog1*Δ strains in the *SPT5-TAP* background (upper panel) and in the untagged background (lower panel). Exponentially growing cultures were spotted on SC medium without drug, or with 25 or 50 µg/ml mycophenolic acid (MPA), alone or in combination with 0.1 mg/ml guanine. Plates were incubated at 30 °C and 37 °C for 2–3 days before imaging. The spot assays shown in Figure 2C and 2D are taken from these plates.

**Figure S2.**
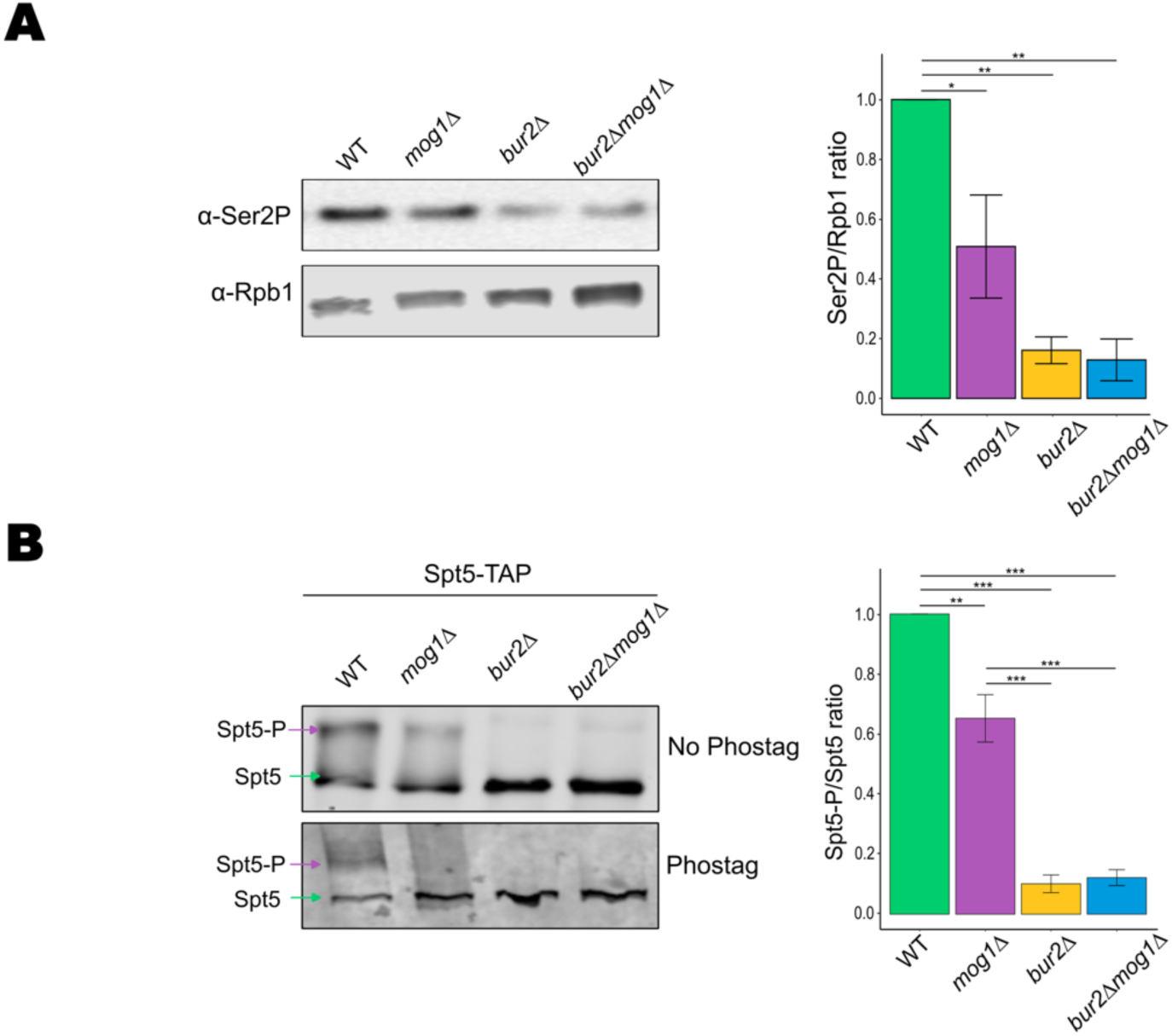
Ser2-P and Spt5 phosphorylation in *mog1*Δ, *bur2*Δ and *bur2*Δ *mog1*Δ cells. (A) Western blot analysis of total phosphorylated RNAPII CTD at serine 2 (Ser2-P) in wild-type (WT), *mog1*Δ, *bur2*Δ and *bur2*Δ *mog1*Δ strains. Rpb1 was used as loading control. Quantification of Ser2-P/Rpb1 signal intensity is shown on the right, normalized to WT. Data represent mean ± standard error from n = 3 biological replicates. Statistical significance was determined using one-way ANOVA followed by Tukey’s multiple comparison test. (B) Western blot analysis of Spt5 phosphorylation in WT, *mog1*Δ, *bur2*Δ and *bur2*Δ *mog1*Δ strains in the *SPT5-TAP* background. The upper panel shows Spt5-TAP under standard SDS–PAGE, and the lower panel shows Phos-tag electrophoresis. Green arrows indicate unphosphorylated Spt5, and purple arrows indicate the phosphorylated form (Spt5-P). Quantification (right) shows Spt5-P/Spt5 ratios normalized to WT. Values represent mean ± standard error from n = 3 biological replicates, quantified on two independent membranes. Statistical significance was determined using one-way ANOVA followed by Tukey’s multiple comparison test. For all panels, values are shown relative to WT (=1), with standard errors propagated using first-order Taylor expansion (delta method). The data shown in Figure 1B and 1D are a subset of the data in this figure. *P = 0.01–0.05; **P < 0.01; ***P < 0.001.

**Figure S3.**
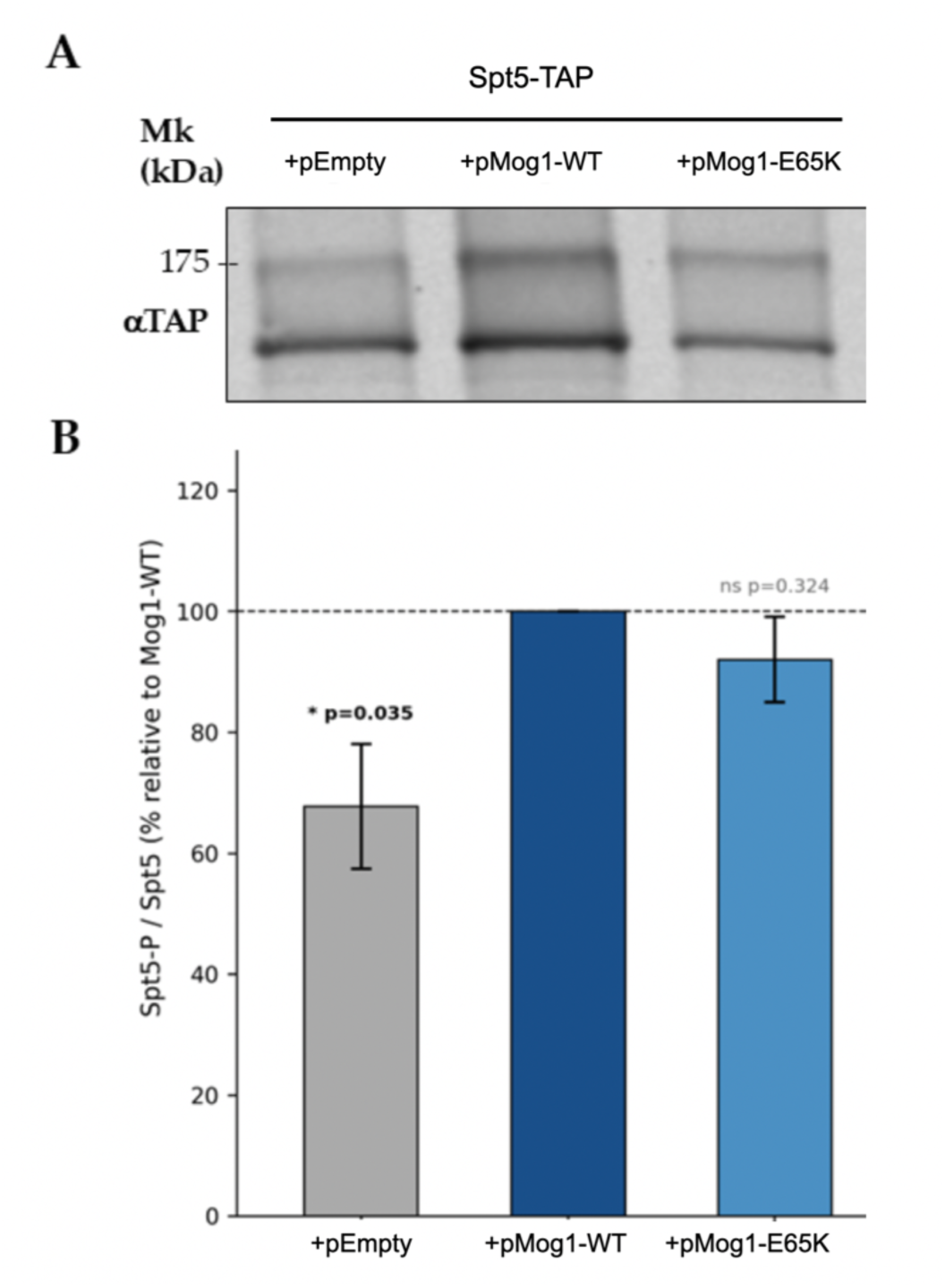
Mog1-E65K point substitution does not affect Spt5 phosphorylation. (A) Representative western blot of Spt5 phosphorylation in *mog1*Δ cells expressing empty vector, Mog1-WT or Mog1-E65K from a centromeric plasmid in a Spt5-TAP strain. The slower-migrating phosphorylated form (Spt5P, ∼134 kDa) and the unphosphorylated form (Spt5, ∼134 kDa). (B) Bar charts show the Spt5-P/(Spt5-P + Spt5) ratio (%). Data represent mean ± SEM of n = 3 independent biological replicates. Statistical comparisons vs. WT: two-tailed Student’s t-test; *, p < 0.05; ns, not significant.

## List of SUPPLEMENTARY Tables

| Table | Title | M&M section |
|---|---|---|
| Supplementary Table S1 | *Saccharomyces cerevisiae strains used in this study* | Yeast strains, plasmids and growth conditions |
| Supplementary Table S2 | *Plasmids used in this study* | Yeast strains, plasmids and growth conditions |
| Supplementary Table S3 | *Oligonucleotides used in this study (strain construction, sequencing and qPCR primers)* | Yeast strains…; y ChIP-qPCR |
| Supplementary Table S4 | *Proteins identified by tandem affinity purification and mass spectrometry* | Tandem affinity purification (TAP) and mass spectrometry |

## Notes

### Competing Interest Statement

The authors have declared no competing interest.

## REFERENCES

1. Cramer, P. (2019) Organization and regulation of gene transcription. Nature, 573, 45–54.

2. Hahn, S. and Young, E.T. (2011) Transcriptional Regulation in Saccharomyces cerevisiae: Transcription Factor Regulation and Function, Mechanisms of Initiation, and Roles of Activators and Coactivators. Genetics, 189, 705–736.

3. Harlen, K.M. and Churchman, L.S. (2017) The code and beyond: transcription regulation by the RNA polymerase II carboxy-terminal domain. Nat Rev Mol Cell Biol, 18, 263–273.

4. Jeronimo, C., Collin, P. and Robert, F. (2016) The RNA Polymerase II CTD: The Increasing Complexity of a Low-Complexity Protein Domain. Journal of Molecular Biology, 428, 2607–2622.

5. Chen, F.X., Smith, E.R. and Shilatifard, A. (2018) Born to run: control of transcription elongation by RNA polymerase II. Nature Reviews Molecular Cell Biology, 19, 464–478.

6. Decker, T.-M. (2021) Mechanisms of Transcription Elongation Factor DSIF (Spt4– Spt5). Journal of Molecular Biology, 433, 166657.

7. Aoi, Y., Takahashi, Y.-H., Shah, A.P., et al. (2021) SPT5 stabilization of promoter-proximal RNA polymerase II. Mol Cell, 81, 4413–4424.e5.

8. Viktorovskaya, O.V., Appling, F.D. and Schneider, D.A. (2011) Yeast Transcription Elongation Factor Spt5 Associates with RNA Polymerase I and RNA Polymerase II Directly. Journal of Biological Chemistry, 286, 18825–18833.

9. Booth, G.T., Wang, I.X., Cheung, V.G., et al. (2016) Divergence of a conserved elongation factor and transcription regulation in budding and fission yeast. Genome Res, 26, 799–811.

10. Uzun, Ü., Brown, T., Fischl, H., et al. (2021) Spt4 facilitates the movement of RNA polymerase II through the +2 nucleosomal barrier. Cell Rep, 36, 109755.

11. Keogh, M.-C., Podolny, V. and Buratowski, S. (2003) Bur1 Kinase Is Required for Efficient Transcription Elongation by RNA Polymerase II. Molecular and Cellular Biology, 23, 7005–7018.

12. Liu, Y., Warfield, L., Zhang, C., et al. (2009) Phosphorylation of the transcription elongation factor Spt5 by yeast Bur1 kinase stimulates recruitment of the PAF complex. Mol Cell Biol, 29, 4852–4863.

13. Wada, T., Takagi, T., Yamaguchi, Y., et al. (1998) DSIF, a novel transcription elongation factor that regulates RNA polymerase II processivity, is composed of human Spt4 and Spt5 homologs. Genes Dev, 12, 343–356.

14. Yamada, T., Yamaguchi, Y., Inukai, N., et al. (2006) P-TEFb-mediated phosphorylation of hSpt5 C-terminal repeats is critical for processive transcription elongation. Mol Cell, 21, 227–237.

15. Laribee, R.N., Krogan, N.J., Xiao, T., et al. (2005) BUR Kinase Selectively Regulates H3 K4 Trimethylation and H2B Ubiquitylation through Recruitment of the PAF Elongation Complex. Current Biology, 15, 1487–1493.

16. Qiu, H., Hu, C., Gaur, N.A., et al. (2012) Pol II CTD kinases Bur1 and Kin28 promote Spt5 CTR-independent recruitment of Paf1 complex. The EMBO Journal, 31, 3494–3505.

17. Wood, A., Schneider, J., Dover, J., et al. (2005) The Bur1/Bur2 complex is required for histone H2B monoubiquitination by Rad6/Bre1 and histone methylation by COMPASS. Mol Cell, 20, 589–599.

18. Fetian, T., Grover, A. and Arndt, K.M. (2024) Histone H2B ubiquitylation: Connections to transcription and effects on chromatin structure. Biochimica et Biophysica Acta (BBA) - Gene Regulatory Mechanisms, 1867, 195018.

19. Bowman, E.A. and Kelly, W.G. (2014) RNA polymerase II transcription elongation and Pol II CTD Ser2 phosphorylation: A tail of two kinases. Nucleus, 5, 224–236.

20. Namjilsuren, S. and Arndt, K.M. (2026) Glc7/PP1 triggers Paf1 complex dissociation from RNA polymerase II to enable transcription termination. Genes Dev, 40, 699–716.

21. Baker, R.P., Harreman, M.T., Eccleston, J.F., et al. (2001) Interaction between Ran and Mog1 Is Required for Efficient Nuclear Protein Import. Journal of Biological Chemistry, 276, 41255–41262.

22. Oki, M. and Nishimoto, T. (1998) A protein required for nuclear-protein import, Mo g1p, directly interacts with GTP-Gsp1p, the Saccharomyces cerevisiae Ran homologue. Proceedings of the National Academy of Sciences of the United States of America, 95, 15388–15393.

23. Stewart, M. and Baker, R.P. (2000) 1.9 A resolution crystal structure of the Saccharomyces cerevisiae Ran-binding protein Mog1p. J Mol Biol, 299, 213–223.

24. Chakrabarti, S., Wu, X., Yang, Z., et al. (2013) MOG1 rescues defective trafficking of Na(v)1.5 mutations in Brugada syndrome and sick sinus syndrome. Circ Arrhythm Electrophysiol, 6, 392–401.

25. Wu, L., Yong, S.L., Fan, C., et al. (2008) Identification of a new co-factor, MOG1, required for the full function of cardiac sodium channel Nav 1.5. J Biol Chem, 283, 6968–6978.

26. Oliete-Calvo, P., Serrano-Quílez, J., Nuño-Cabanes, C., et al. (2018) A role for Mog1 in H2Bub1 and H3K4me3 regulation affecting RNAPII transcription and mRNA export. EMBO Rep, 19.

27. Serrano-Quílez, J. and Rodriguez-Navarro, S. (2025) Unraveling gene expression: a beginner’s guide from chromatin modifications to mRNA export in Saccharomyces cerevisiae. Nucleus, 16, 2516909.

28. Luciano, P., Park, K., Audebert, S., et al. (2026) A systematic interactome of *Saccharomyces cerevisiae* SET1C expands its functional landscape and identifies candidate regulatory connections. eLife, 15, RP109886.

29. Guo, C., Si, S., Fang, H., et al. (2025) LEDGF/p75 promotes transcriptional pausing through preventing SPT5 phosphorylation. Sci Adv, 11, eadr2131.

30. Sun, R. and Fisher, R.P. (2025) Tripartite phosphorylation of SPT5 by CDK9 times pause release and tunes elongation rate of RNA polymerase II. Mol Cell, 85, 1743–1759.e5.

31. Kinoshita, E., Kinoshita-Kikuta, E. and Koike, T. (2009) Separation and detection of large phosphoproteins using Phos-tag SDS-PAGE. Nature Protocols, 4, 1513–1521.

32. Janke, C., Magiera, M.M., Rathfelder, N., et al. (2004) A versatile toolbox for PCR-based tagging of yeast genes: new fluorescent proteins, more markers and promoter substitution cassettes. Yeast, 21, 947–962.

33. Longtine, M.S., McKenzie, A. 3rd, Demarini, D.J., et al. (1998) Additional modules for versatile and economical PCR-based gene deletion and modification in Saccharomyces cerevisiae. Yeast, **14**, 953–961.

34. Gietz, R.D. and Schiestl, R.H. (2007) High-efficiency yeast transformation using the LiAc/SS carrier DNA/PEG method. Nat Protoc, 2, 31–34.

35. Mans, R., Wijsman, M., Daran-Lapujade, P., et al. (2018) A protocol for introduction of multiple genetic modifications in Saccharomyces cerevisiae using CRISPR/Cas9. FEMS Yeast Res, 18.

36. Concordet, J.-P. and Häussler, M. (2018) CRISPOR: intuitive guide selection for CRISPR/Cas9 genome editing experiments and screens. Nucleic Acids Res, 46, W242–W245.

37. Keogh, M.-C., Kim, J.-A., Downey, M., et al. (2006) A phosphatase complex that dephosphorylates gammaH2AX regulates DNA damage checkpoint recovery. Nature, 439, 497–501.

38. Puig, O., Caspary, F., Rigaut, G., et al. (2001) The tandem affinity purification (TAP) method: a general procedure of protein complex purification. Methods, 24, 218–229.

39. Rigaut, G., Shevchenko, A., Rutz, B., et al. (1999) A generic protein purification method for protein complex characterization and proteome exploration. Nat Biotechnol, 17, 1030–1032.

40. Kunzmann, P. and Hamacher, K. (2018) Biotite: a unifying open source computational biology framework in Python. BMC Bioinformatics, 19, 346.

41. Schrödinger, LLC (2015) The AxPyMOL Molecular Graphics Plugin for Microsoft PowerPoint, Version 1.8.

42. Riles, L., Shaw, R.J., Johnston, M., et al. (2004) Large-scale screening of yeast mutants for sensitivity to the IMP dehydrogenase inhibitor 6-azauracil. Yeast, 21, 241–248.

43. Ding, B., LeJeune, D. and Li, S. (2010) The C-terminal Repeat Domain of Spt5 Plays an Important Role in Suppression of Rad26-independent Transcription Coupled Repair*. Journal of Biological Chemistry, 285, 5317–5326.

44. Yao, S. and Prelich, G. (2002) Activation of the Bur1-Bur2 cyclin-dependent kinase complex by Cak1. Mol Cell Biol, 22, 6750–6758.

45. Cho, E.J., Kobor, M.S., Kim, M., et al. (2001) Opposing effects of Ctk1 kinase and Fcp1 phosphatase at Ser 2 of the RNA polymerase II C-terminal domain. Genes Dev, 15, 3319–3329.

46. Sigurdsson, S., Dirac-Svejstrup, A.B. and Svejstrup, J.Q. (2010) Evidence that transcript cleavage is essential for RNA polymerase II transcription and cell viability. Mol Cell, 38, 202–210.

47. Separovich, R.J. and Wilkins, M.R. (2021) Ready, SET, Go: Post-translational regulation of the histone lysine methylation network in budding yeast. Journal of Biological Chemistry, 297, 100939.

48. Badjatia, N., Rossi, M.J., Bataille, A.R., et al. (2021) Acute stress drives global repression through two independent RNA polymerase II stalling events in Saccharomyces. Cell Reports, 34, 108640.

49. Bai, X., Wang, Z., Xiong, H., et al. (2025) Molecular mechanism for the interaction of MOG1 with the intracellular loop II of cardiac sodium channel Na(v)1.5 and its role in arrhythmias. J Mol Cell Cardiol, 205, 68–83.

