## Supplementary material for "The Ran-binding protein Mog1 sustains Spt5 C-terminal phosphorylation by counteracting Spt4 in budding yeast": Supplemntal Tables S1-4

### Table S1. Saccharomyces cerevisiae strains used in this study

| Strain | Genotype | Source |
|---|---|---|
| **BY4741** | MATa his3Δ1 leu2Δ0 met15Δ0 ura3Δ0 | Euroscarf |
| **BY4742** | MATα his3Δ1 leu2Δ0 lys2Δ0 ura3Δ0 | Euroscarf |
| **mog1Δ** | MATa his3Δ1 leu2Δ0 met15Δ0 ura3Δ0 mog1::HIS3 | Oliete-Calvo et al. (2018) |
| **mog1Δ** | MATa his3Δ1 leu2Δ0 met15Δ0 ura3Δ0 mog1::LEU2 | This study |
| **mog1Δ** | MATa his3Δ1 leu2Δ0 met15Δ0 ura3Δ0 mog1::HPHMXNT1 | This study |
| **mog1Δ** | MATα his3Δ1 leu2Δ0 lys2Δ0 ura3Δ0 mog1::HIS3 | This study |
| **MOG1-E65K** | MATa his3Δ1 leu2Δ0 met15Δ0 ura3Δ0 MOG1-E65K | This study |
| **Mog1-TAP** | MATa his3Δ1 leu2Δ0 met15Δ0 ura3Δ0 MOG1-TAP::URA3 | Oliete-Calvo et al. (2018) |
| **Mog1-E65K-TAP** | MATa his3Δ1 leu2Δ0 met15Δ0 ura3Δ0 MOG1-E65K-TAP::URA3 | This study |
| **jhd2Δ** | MATa his3Δ1 leu2Δ0 met15Δ0 ura3Δ0 jhd2::KANMX4 | Euroscarf |
| **jhd2Δ mog1Δ** | MATa his3Δ1 leu2Δ0 met15Δ0 ura3Δ0 jhd2::KANMX4 mog1::LEU2 | This study |
| **bur2Δ** | MATa his3Δ1 leu2Δ0 met15Δ0 ura3Δ0 bur2::KANMX4 | This study |
| **bur2Δ mog1Δ** | MATα his3Δ1 leu2Δ0 ura3Δ0 bur2::KANMX4 mog1::LEU2 | This study |
| **Bur1-HA** | MATa his3Δ200 leu2Δ1 trp1Δ63 ura3-52 lys2Δ202 BUR1-3xHA::TRP1 | Stephen Buratowski's laboratory |
| **Bur1-HA mog1Δ** | MATa his3Δ200 leu2Δ1 trp1Δ63 ura3-52 lys2Δ202 BUR1-3xHA::TRP1 mog1::KANMX4 | This study |
| **bur1-80-HA** | MATa his3Δ200 leu2Δ1 trp1Δ63 ura3-52 lys2Δ202 bur1::HIS3 [pRS315-bur1-80-3xHA] | Stephen Buratowski's laboratory |
| **bur1-80-HA mog1Δ** | MATa his3Δ200 leu2Δ1 trp1Δ63 ura3-52 lys2Δ202 bur1::HIS3 mog1::KANMX4 [pRS315-bur1-80-3xHA] | This study |
| **Spt5-TAP** | MATa his3Δ1 leu2Δ0 ura3Δ0 met15Δ0 SPT5-TAP::HIS3 | This study |
| **Spt5-TAP** | MATα his3Δ1 leu2Δ0 ura3Δ0 lys2Δ0 SPT5-TAP::HIS3 | This study |
| **Spt5-TAP mog1Δ** | MATa his3Δ1 leu2Δ0 ura3Δ0 SPT5-TAP::HIS3 mog1::LEU2 | This study |
| **Spt5-TAP mog1Δ** | MATα his3Δ1 leu2Δ0 ura3Δ0 SPT5-TAP::HIS3 mog1::LEU2 | This study |
| **Spt5-TAP bur2Δ** | MATα his3Δ1 leu2Δ0 ura3Δ0 SPT5-TAP::HIS3 bur2::KANMX4 | This study |
| **Spt5-TAP bur2Δ** | MATa his3Δ1 leu2Δ0 ura3Δ0 SPT5-TAP::HIS3 bur2::KANMX4 | This study |
| **Spt5-TAP bur2Δ mog1Δ** | MATα his3Δ1 leu2Δ0 ura3Δ0 SPT5-TAP::HIS3 bur2::KANMX4 mog1::LEU2 | This study |
| **Spt5-TAP bur2Δ mog1Δ** | MATa his3Δ1 leu2Δ0 ura3Δ0 SPT5-TAP::HIS3 bur2::KANMX4 mog1::LEU2 | This study |
| **spt4Δ** | MATα his3Δ1 leu2Δ0 ura3Δ0 SPT5-TAP::HIS3 bur2::KANMX4 mog1::LEU2 | This study |
| **spt4Δ** | MATα his3Δ1 leu2Δ0 ura3Δ0 spt4::HPHMXNT1 | This study |
| **mog1Δ spt4Δ** | MATa his3Δ1 leu2Δ0 met15Δ0 ura3Δ0 spt4::HPHMXNT1 mog1::HIS3 | This study |
| **mog1Δ spt4Δ** | MATα his3Δ1 leu2Δ0 ura3Δ0 spt4::HPHMXNT1 mog1::HIS3 | This study |
| **Spt4-GFP** | MATa his3Δ1 leu2Δ0 ura3Δ0 SPT4-EGFP::HIS3 | This study |
| **Spt4-GFP mog1Δ** | MATa his3Δ1 leu2Δ0 ura3Δ0 SPT4-EGFP::HIS3 mog1::URA3 | This study |
| **Spt5-TAP spt4Δ** | MATa his3Δ1 leu2Δ0 ura3Δ0 SPT5-TAP::HIS3 spt4::HPHMXNT1 | This study |
| **Spt5-TAP spt4Δ** | MATα his3Δ1 leu2Δ0 ura3Δ0 SPT5-TAP::HIS3 spt4::HPHMXNT1 | This study |
| **Spt5-TAP spt4Δ mog1Δ** | MATa his3Δ1 leu2Δ0 ura3Δ0 SPT5-TAP::HIS3 mog1::LEU2 spt4::HPHMXNT1 | This study |
| **Spt5-TAP spt4Δ mog1Δ** | MATα his3Δ1 leu2Δ0 ura3Δ0 SPT5-TAP::HIS3 mog1::LEU2 spt4::HPHMXNT1 | This study |
| **ctk1Δ** | MATa his3Δ1 leu2Δ0 met15Δ0 ura3Δ0 ctk1::KANMX4 | Euroscarf |
| **ctk1Δ** | MATα his3Δ1 leu2Δ0 ura3Δ0 ctk1::KANMX4 | This study |

| Strain | Genotype | Source |
|---|---|---|
| **ctk1Δ mog1Δ** | MATa his3Δ1 leu2Δ0 ura3Δ0 ctk1::KANMX4 mog1::HIS3 | This study |
| **ctk1Δ mog1Δ** | MATα his3Δ1 leu2Δ0 ura3Δ0 ctk1::KANMX4 mog1::HIS3 | This study |
| **dst1Δ** | MATa his3Δ1 leu2Δ0 ura3Δ0 dst1::KANMX4 | This study |
| **dst1Δ** | MATα his3Δ1 leu2Δ0 ura3Δ0 lys2Δ0 dst1::KANMX4 | Sebastián Chávez's laboratory |
| **dst1Δ mog1Δ** | MATa his3Δ1 leu2Δ0 ura3Δ0 dst1::KANMX4 mog1::HIS3 | This study |
| **dst1Δ mog1Δ** | MATα his3Δ1 leu2Δ0 ura3Δ0 dst1::KANMX4 mog1::HIS3 | This study |

Genotypes are shown for experimentally established genetic markers. LYS2 and MET15 status was not determined for tetrad-derived strains and is therefore not indicated.

#### Table S2. Plasmids used in this study

| Plasmid | Description | Source |
|---|---|---|
| **CRISPR-Cas9 genome editing** | | |
| **pMEL10** | sgRNA expression plasmid (KlURA3 marker); linearised and co-transformed with the sgRNA fragment for in vivo assembly | Mans et al. (2015) |
| **pJH001** | Cas9 expression plasmid (LEU2 marker) | Laughery et al. (2015) |
| **Mog1 expression plasmids (centromeric)** | | |
| **pRS315** | Empty centromeric vector (LEU2), used as control | Sikorski & Hieter (1989) |
| **pMOG1** | pRS315-based plasmid expressing wild-type Mog1 | Lu et al. (2004) |
| **pMOG1-E65K** | pRS315-based plasmid expressing Mog1-E65K | This study |
| **pMOG1-W145A** | pRS315-based plasmid expressing Mog1-W145A | This study |
| **pMOG1-ΔK144-T153** | pRS315-based plasmid expressing Mog1 lacking residues K144–T153 | This study |
| **BUR1** | | |
| **pRS315-BUR1-3xHA** | Centromeric (LEU2) pRS315-based plasmid expressing wild-type BUR1 with a C-terminal 3xHA tag | Stephen Buratowski’s laboratory |
| **pRS315-bur1-80-3xHA** | Centromeric (LEU2) pRS315-based plasmid expressing the conditional bur1-80 allele with a C-terminal 3xHA tag | Stephen Buratowski’s laboratory |
| **Subcellular localisation markers (centromeric)** | | |
| **pSec63-mCherry** | Centromeric (LEU2) YCplac111-based plasmid expressing Sec63-mCherry (nuclear envelope / ER marker). | Ethel Queralt’s laboratory |
| **pPus1-mCherry** | Centromeric (HIS3) pRS313-based plasmid expressing Pus1-mCherry (nucleoplasmic marker). | Ethel Queralt’s laboratory |
| **Template plasmids for PCR-based gene deletion and tagging** | | |
| **pFA6-KANMX4** | Template for KANMX4 deletion cassettes | Wach et al. (1994) |
| **pFA6-hphNT1** | Template for hphNT1 deletion cassettes | Janke et al. (2004) |
| **pRS403** | Template for HIS3 deletion cassettes | Sikorski & Hieter (1989) |
| **pRS405** | Template for LEU2 deletion cassettes | Sikorski & Hieter (1989) |
| **pRS406** | Template for URA3 deletion cassettes | Sikorski & Hieter (1989) |
| **pBS1539** | Template for C-terminal TAP (CBP-TEV-Protein A) tagging with URA3 marker | Puig et al. (2001) |
| **pGFP-HIS3** | Template for C-terminal GFP tagging with HIS3 marker | Ethel Queralt’s laboratory |
| **pNet1-mCherry** | Template for C-terminal mCherry tagging with URA3 marker | Ethel Queralt’s laboratory |

#### Table S3. Oligonucleotides used in this study (strain construction, sequencing and qPCR primers)

| Name | Target | Region / type | Sequence (5′–3′) |
|---|---|---|---|
| **Primers used for qPCR** | | | |
| **Intergenic-FW** | ChrV intergenic | Forward | TGTTCCTTTAAGAGGTGATGGTGA |
| **Intergenic-RV** | ChrV intergenic | Reverse | GTGCGCAGTACTTGTGAAAACC |
| **ADH1-FW** | ADH1 | 5′ ORF | ACGAATCCCACGGTAAGTTG |
| **ADH1-RV** | ADH1 | 5′ ORF | AAGCGTGCAAGTCAGTGTGA |
| **ADH1-FW** | ADH1 | 3′ ORF | CTTTGGACTTCTTCGCCAGA |
| **ADH1-RV** | ADH1 | 3′ ORF | CAACGATTTGACCCTTTTCC |
| **PMA1-FW** | PMA1 | 5′ ORF | CAGCTCATCAGCCAACTCAA |
| **PMA1-RV** | PMA1 | 5′ ORF | TCGTCGACACCGTGATTAGA |
| **PMA1-FW** | PMA1 | 3′ ORF | TTACTGTCGTCCGTGTCTGG |
| **PMA1-RV** | PMA1 | 3′ ORF | CATTGGCTTACCGTTCATCA |
| **YEF3-FW** | YEF3 | 5′ ORF | CACTGCTGACAACAGACACG |
| **YEF3-RV** | YEF3 | 5′ ORF | TTGATACCCTTGGCCAATTC |
| **YEF3-FW** | YEF3 | 3′ ORF | CAAGGTGCTGGTCCAAGAAT |
| **YEF3-RV** | YEF3 | 3′ ORF | TCTCAATTCCGCAGAAGACA |
| **PYK1-FW** | PYK1 | 5′ ORF | ACCAAGGGTCCAGAAATCAG |
| **PYK1-RV** | PYK1 | 5′ ORF | TGTCATCGGTGGTGAAGAT |
| **PYK1-FW** | PYK1 | 3′ ORF | AACGCCGTTACCACTATGGC |
| **PYK1-RV** | PYK1 | 3′ ORF | TAACCAAGATGATTGGACAGTTTGG |
| **Oligonucleotides used for CRISPR mutagenesis** | | | |
| **CRISPR plasmid backbone** | CRISPR plasmid | Forward | GATCATTTATCTTTCACTGCGGAGAAG |
| **CRISPR plasmid backbone** | CRISPR plasmid | Reverse | GTTTTAGAGCTAGAAATAGCAAGTTAAAATAAGGCTAGTC |
| **Mog1-E65K CRISPR crRNA** | MOG1 | Forward | TGCGCATGTTTCGGCGTTCGAAACTTCTCCGCAGTGAAAGATAAATGATCGACGTACACCTCCTGTGTATGTTTTAGAGCTAGAAATAGCAAGTTAAAATAAG |
| **Mog1-E65K CRISPR crRNA** | MOG1 | Reverse | GTTGATAACGGACTAGCCTTATTTTAACTTGCTATTTCTAGCTCTAAAACATACACAGGAGGTGTACGTCGATCATTTATCTTTCACTGCGGAGAAGTTTCGAACGCCGAAACATGCGCA |
| **Mog1-E65K CRISPR donor** | MOG1 | Donor | ACTCTAAGAGAAGTCCCCGATACACAGAAGGTGTACGTCAATTCCCGTCGCGATGAAGAA |
| **Mog1-E65K CRISPR donor** | MOG1 | Forward | ACGGCGGCGCCATCACTACTGTCGTCCCACCTGGGTTTATAGACGCTTCTACTCTAAGAG |
| **Mog1-E65K CRISPR donor** | MOG1 | Reverse | AAGAGATCCACGATAATGCTTTCATTGGTGGCTAGTCCATCTTCGAATTCTTCTTCATCG |

#### Table S4. Proteins identified by tandem affinity purification and mass spectrometry

Selected columns are shown (gene, UniProt ID, description, length, coverage, peptide and spectral counts). Complete data, including protein probabilities, intensities and modifications, are provided in the accompanying Excel file.

##### Sheet 1. Pellet Mog1-TAP (808 proteins)

| Gene | Protein ID | Protein description | Length (aa) | Coverage (%) | Total peptides | Unique peptides | Total spectral count |
|---|---|---|---|---|---|---|---|
| ECM33 | A6ZL22 | Cell wall protein ECM33 | 429 | 3.50 | 1 | 0 | 2 |
| RPS1B | A6ZM02 | Small ribosomal subunit protein eS1B | 255 | 67.84 | 22 | 0 | 116 |
| TIF34 | A6ZMK5 | Eukaryotic translation initiation factor 3 subunit I | 347 | 32.56 | 7 | 0 | 21 |
| NIP1 | A6ZN26 | Eukaryotic translation initiation factor 3 subunit C | 812 | 2.34 | 2 | 0 | 3 |
| DBP5 | A6ZNQ1 | ATP-dependent RNA helicase DBP5 | 482 | 11.62 | 5 | 0 | 19 |
| DED1 | A6ZP47 | ATP-dependent RNA helicase DED1 | 604 | 66.06 | 31 | 0 | 160 |
| NOP58 | A6ZPE5 | Nucleolar protein 58 | 511 | 15.46 | 6 | 0 | 17 |
| PRT1 | A6ZPJ1 | Eukaryotic translation initiation factor 3 subunit B | 765 | 28.24 | 20 | 0 | 41 |
| IML2 | A6ZPP2 | Inclusion body clearance protein IML2 | 731 | 3.42 | 2 | 0 | 2 |
| TIF1 | A6ZQJ1 | ATP-dependent RNA helicase eIF4A | 395 | 52.15 | 17 | 0 | 49 |
| MDV1 | A6ZQL5 | Mitochondrial division protein 1 | 714 | 2.24 | 1 | 0 | 1 |
| ADE12 | A6ZRM0 | Adenylosuccinate synthetase | 433 | 12.01 | 4 | 0 | 9 |
| TOM70 | A6ZRW3 | Mitochondrial import receptor subunit TOM70 | 617 | 5.83 | 2 | 0 | 6 |
| DCP2 | A6ZRW5 | mRNA-decapping enzyme subunit 2 | 970 | 1.55 | 1 | 0 | 1 |
| DBP2 | A6ZRX0 | ATP-dependent RNA helicase DBP2 | 546 | 43.96 | 21 | 0 | 70 |
| CRP1 | A6ZT54 | Cruciform DNA-recognizing protein 1 | 465 | 3.23 | 1 | 0 | 1 |
| TDA11 | A6ZT67 | Topoisomerase I damage affected protein 11 | 507 | 9.86 | 4 | 0 | 9 |
| EGD2 | A6ZT99 | Nascent polypeptide-associated complex subunit alpha | 174 | 44.83 | 6 | 0 | 20 |
| RPS0A | A6ZUM5 | Small ribosomal subunit protein uS2A | 252 | 61.11 | 11 | 0 | 79 |
| EGD1 | A6ZWL1 | Nascent polypeptide-associated complex subunit beta-1 | 157 | 25.48 | 2 | 0 | 7 |
| DHH1 | A6ZXG9 | ATP-dependent RNA helicase DHH1 | 506 | 11.07 | 5 | 0 | 11 |
| SUB2 | A6ZXP4 | ATP-dependent RNA helicase SUB2 | 446 | 9.87 | 3 | 0 | 9 |
| ARO1 | A6ZY89 | Pentafunctional AROM polypeptide | 1588 | 1.57 | 2 | 0 | 1 |
| ADK1 | A6ZYI0 | Adenylate kinase | 222 | 32.88 | 7 | 0 | 22 |
| MIC60 | A6ZZY0 | MICOS complex subunit MIC60 | 539 | 2.78 | 1 | 0 | 2 |
| RPS0B | A7A0V3 | Small ribosomal subunit protein uS2B | 252 | 73.41 | 13 | 0 | 82 |
| HRI1 | A7A1I2 | Protein HRI1 | 244 | 19.67 | 4 | 0 | 7 |
| MRPL4 | A7A1V9 | Large ribosomal subunit protein uL29m | 319 | 2.82 | 1 | 0 | 2 |
| RPS1A | A7A1W1 | Small ribosomal subunit protein eS1A | 255 | 67.84 | 22 | 0 | 109 |
| EIS1 | C7GL88 | Eisosome protein 1 | 843 | 18.39 | 14 | 0 | 30 |
| RPS9A | O13516 | Small ribosomal subunit protein uS4A | 197 | 52.28 | 14 | 0 | 40 |
| RPL36B | O14455 | Large ribosomal subunit protein eL36B | 100 | 40.00 | 5 | 0 | 16 |
| MBF1 | O14467 | Multiprotein-bridging factor 1 | 151 | 24.50 | 3 | 0 | 5 |

| Gene | Protein ID | Protein description | Length (aa) | Coverage (%) | Total peptides | Unique peptides | Total spectral count |
|---|---|---|---|---|---|---|---|
| KRT86 | O43790 | Keratin, type II cuticular Hb6 | 486 | 5.56 | 2 | 0 | 3 |
| TIM9 | O74700 | Mitochondrial import inner membrane translocase subunit TIM9 | 87 | 40.23 | 3 | 0 | 10 |
| KRT33A | O76009 | Keratin, type I cuticular Ha3-I | 404 | 11.63 | 5 | 0 | 9 |
| KRT34 | O76011 | Keratin, type I cuticular Ha4 | 436 | 10.55 | 5 | 0 | 8 |
| KRT15 | O77727 | Keratin, type I cytoskeletal 15 | 453 | 20.97 | 12 | 7 | 67 |
| ADH1 | P00330 | Alcohol dehydrogenase 1 | 348 | 54.60 | 18 | 0 | 89 |
| TDH2 | P00358 | Glyceraldehyde-3-phosphate dehydrogenase 2 | 332 | 64.76 | 24 | 0 | 298 |
| TDH3 | P00359 | Glyceraldehyde-3-phosphate dehydrogenase 3 | 332 | 73.19 | 28 | 0 | 301 |
| TDH1 | P00360 | Glyceraldehyde-3-phosphate dehydrogenase 1 | 332 | 62.35 | 19 | 0 | 140 |
| SOD1 | P00445 | Superoxide dismutase [Cu-Zn] | 154 | 65.58 | 9 | 0 | 31 |
| SOD2 | P00447 | Superoxide dismutase [Mn], mitochondrial | 233 | 12.02 | 2 | 0 | 4 |
| CDC19 | P00549 | Pyruvate kinase 1 | 500 | 80.20 | 44 | 0 | 215 |
| PGK1 | P00560 | Phosphoglycerate kinase | 416 | 76.20 | 28 | 0 | 115 |
| LALBA | P00711 | Alpha-lactalbumin | 142 | 35.21 | 5 | 5 | 15 |
| | P00761 | Trypsin | 231 | 37.66 | 6 | 6 | 343 |
| HIS4 | P00815 | Histidine biosynthesis trifunctional protein | 799 | 16.52 | 11 | 0 | 32 |
| IPP1 | P00817 | Inorganic pyrophosphatase | 287 | 35.89 | 9 | 0 | 29 |
| ATP2 | P00830 | ATP synthase subunit beta, mitochondrial | 511 | 23.68 | 9 | 0 | 19 |
| CIT1 | P00890 | Citrate synthase, mitochondrial | 479 | 6.47 | 2 | 0 | 4 |
| ENO1 | P00924 | Enolase 1 | 437 | 73.00 | 25 | 0 | 100 |
| ENO2 | P00925 | Enolase 2 | 437 | 81.01 | 29 | 0 | 234 |
| TRP5 | P00931 | Tryptophan synthase | 707 | 16.69 | 8 | 0 | 15 |
| TPI1 | P00942 | Triosephosphate isomerase | 248 | 58.06 | 12 | 0 | 50 |
| GPM1 | P00950 | Phosphoglycerate mutase 1 | 247 | 51.82 | 12 | 0 | 80 |
| MES1 | P00958 | Methionine--tRNA ligase, cytoplasmic | 751 | 2.66 | 1 | 0 | 2 |
| SERPINB14 | P01012 | Ovalbumin | 386 | 10.62 | 3 | 3 | 9 |
| HTB1 | P02293 | Histone H2B.1 | 131 | 14.50 | 3 | 0 | 10 |
| HHF1 | P02309 | Histone H4 | 103 | 33.01 | 3 | 0 | 11 |
| RPP2B | P02400 | Large ribosomal subunit protein P2B | 110 | 53.64 | 4 | 0 | 16 |
| RPL28 | P02406 | Large ribosomal subunit protein uL15 | 149 | 36.24 | 7 | 0 | 25 |
| RPS17A | P02407 | Small ribosomal subunit protein eS17A | 136 | 54.41 | 8 | 0 | 29 |
| TUB2 | P02557 | Tubulin beta chain | 457 | 5.47 | 2 | 0 | 14 |
| CSN1S1 | P02662 | Alpha-S1-casein | 214 | 31.31 | 5 | 5 | 24 |
| CSN1S2 | P02663 | Alpha-S2-casein | 222 | 25.23 | 6 | 6 | 15 |
| CSN2 | P02666 | Beta-casein | 224 | 21.88 | 4 | 4 | 36 |
| CSN3 | P02668 | Kappa-casein | 190 | 29.47 | 4 | 4 | 37 |
| ALB | P02768 | Albumin | 609 | 48.28 | 27 | 22 | 62 |
| ALB | P02769 | Albumin | 607 | 35.09 | 20 | 15 | 51 |

| Gene | Protein ID | Protein description | Length (aa) | Coverage (%) | Total peptides | Unique peptides | Total spectral count |
|---|---|---|---|---|---|---|---|
| LTF | P02788 | Lactotransferrin | 710 | 50.14 | 29 | 29 | 93 |
| HSP82 | P02829 | ATP-dependent molecular chaperone HSP82 | 709 | 41.61 | 31 | 0 | 79 |
| TUF1 | P02992 | Elongation factor Tu, mitochondrial | 437 | 5.49 | 2 | 0 | 2 |
| TEF1 | P02994 | Elongation factor 1-alpha | 458 | 55.24 | 21 | 0 | 182 |
| REP2 | P03872 | Partitioning protein REP2 | 296 | 3.38 | 1 | 0 | 1 |
| CPA2 | P03965 | Carbamoyl-phosphate synthase arginine-specific large chain | 1118 | 1.43 | 1 | 0 | 1 |
| CAT | P04040 | Catalase | 527 | 13.28 | 6 | 6 | 12 |
| RPO31 | P04051 | DNA-directed RNA polymerase III subunit RPC1 | 1460 | 0.68 | 1 | 0 | 1 |
| PAB1 | P04147 | Polyadenylate-binding protein, cytoplasmic and nuclear | 577 | 65.34 | 35 | 0 | 188 |
| KRT1 | P04264 | Keratin, type II cytoskeletal 1 | 644 | 72.36 | 58 | 54 | 841 |
| RPL24A | P04449 | Large ribosomal subunit protein eL24A | 155 | 33.55 | 10 | 0 | 47 |
| RPL25 | P04456 | Large ribosomal subunit protein uL23 | 142 | 42.96 | 6 | 0 | 24 |
| THS1 | P04801 | Threonine--tRNA ligase, cytoplasmic | 734 | 19.21 | 12 | 0 | 26 |
| DPS1 | P04802 | Aspartate--tRNA ligase, cytoplasmic | 557 | 30.88 | 13 | 0 | 38 |
| HXK1 | P04806 | Hexokinase-1 | 485 | 25.98 | 10 | 0 | 25 |
| HXK2 | P04807 | Hexokinase-2 | 486 | 31.07 | 11 | 0 | 38 |
| CDC25 | P04821 | Cell division control protein 25 | 1589 | 1.45 | 1 | 0 | 1 |
| POR1 | P04840 | Mitochondrial outer membrane protein porin 1 | 283 | 29.33 | 7 | 0 | 18 |
| PMA1 | P05030 | Plasma membrane ATPase 1 | 918 | 18.74 | 14 | 0 | 35 |
| RPP0 | P05317 | Large ribosomal subunit protein uL10 | 312 | 36.22 | 11 | 0 | 28 |
| RPP2A | P05319 | Large ribosomal subunit protein P2A | 106 | 52.83 | 4 | 0 | 16 |
| SUP35 | P05453 | Eukaryotic peptide chain release factor GTP-binding subunit | 685 | 38.69 | 23 | 0 | 83 |
| MET6 | P05694 | 5-methyltetrahydropteroyltriglutamate--homocysteine methyltransferase | 767 | 10.56 | 6 | 0 | 11 |
| RPL7A | P05737 | Large ribosomal subunit protein uL30A | 244 | 46.31 | 12 | 0 | 28 |
| RPL9A | P05738 | Large ribosomal subunit protein uL6A | 191 | 31.41 | 7 | 0 | 17 |
| RPL6B | P05739 | Large ribosomal subunit protein eL6B | 176 | 53.41 | 9 | 0 | 21 |
| RPL17A | P05740 | Large ribosomal subunit protein uL22A | 184 | 47.83 | 8 | 0 | 40 |
| RPL26A | P05743 | Large ribosomal subunit protein uL24A | 127 | 49.61 | 11 | 0 | 31 |
| RPL33A | P05744 | Large ribosomal subunit protein eL33A | 107 | 39.25 | 5 | 0 | 12 |
| RPL15A | P05748 | Large ribosomal subunit protein eL15A | 204 | 32.84 | 6 | 0 | 20 |
| RPL22A | P05749 | Large ribosomal subunit protein eL22A | 121 | 61.16 | 4 | 0 | 89 |
| RPS3 | P05750 | Small ribosomal subunit protein uS3 | 240 | 63.75 | 16 | 0 | 49 |
| RPS9B | P05755 | Small ribosomal subunit protein uS4B | 195 | 51.79 | 15 | 0 | 44 |
| RPS13 | P05756 | Small ribosomal subunit protein uS15 | 151 | 58.28 | 9 | 0 | 39 |
| RPS31 | P05759 | Ubiquitin-ribosomal protein eS31 fusion protein | 152 | 44.74 | 6 | 0 | 17 |
| SCP160 | P06105 | Protein SCP160 | 1222 | 16.78 | 17 | 0 | 36 |
| ILV5 | P06168 | Ketol-acid reductoisomerase, mitochondrial | 395 | 49.37 | 14 | 0 | 67 |
| PDC1 | P06169 | Pyruvate decarboxylase isozyme 1 | 563 | 51.51 | 23 | 0 | 224 |

| Gene | Protein ID | Protein description | Length (aa) | Coverage (%) | Total peptides | Unique peptides | Total spectral count |
|---|---|---|---|---|---|---|---|
| LEU4 | P06208 | 2-isopropylmalate synthase | 619 | 2.91 | 1 | 0 | 1 |
| RPS14A | P06367 | Small ribosomal subunit protein uS11A | 137 | 70.80 | 12 | 0 | 31 |
| CDC31 | P06704 | Cell division control protein 31 | 161 | 18.01 | 2 | 0 | 5 |
| RHO1 | P06780 | GTP-binding protein RHO1 | 209 | 13.88 | 3 | 0 | 4 |
| CMD1 | P06787 | Calmodulin | 147 | 53.74 | 7 | 0 | 37 |
| FAS1 | P07149 | Fatty acid synthase subunit beta | 2051 | 13.90 | 20 | 0 | 56 |
| ADE57 | P07244 | Bifunctional purine biosynthetic protein ADE5,7 | 802 | 10.60 | 7 | 0 | 12 |
| ADE3 | P07245 | C-1-tetrahydrofolate synthase, cytoplasmic | 946 | 9.73 | 8 | 0 | 9 |
| ADH3 | P07246 | Alcohol dehydrogenase 3, mitochondrial | 375 | 4.00 | 2 | 0 | 6 |
| ATP1 | P07251 | ATP synthase subunit alpha, mitochondrial | 545 | 19.08 | 9 | 0 | 20 |
| URA2 | P07259 | Protein URA2 | 2214 | 11.74 | 21 | 0 | 49 |
| CDC33 | P07260 | Eukaryotic translation initiation factor 4E | 213 | 27.70 | 5 | 0 | 10 |
| GDH1 | P07262 | NADP-specific glutamate dehydrogenase 1 | 454 | 11.01 | 3 | 0 | 3 |
| HTS1 | P07263 | Histidine--tRNA ligase, mitochondrial | 546 | 7.33 | 3 | 0 | 5 |
| LEU1 | P07264 | 3-isopropylmalate dehydratase | 779 | 3.34 | 2 | 0 | 4 |
| PEP4 | P07267 | Saccharopepsin | 405 | 19.26 | 5 | 0 | 11 |
| PFY1 | P07274 | Profilin | 126 | 19.05 | 2 | 0 | 2 |
| RAD2 | P07276 | DNA repair protein RAD2 | 1031 | 13.87 | 11 | 0 | 34 |
| RPS19A | P07280 | Small ribosomal subunit protein eS19A | 144 | 59.72 | 10 | 0 | 50 |
| SEC53 | P07283 | Phosphomannomutase | 254 | 18.90 | 4 | 0 | 4 |
| SES1 | P07284 | Serine--tRNA ligase, cytoplasmic | 462 | 12.34 | 5 | 0 | 7 |
| CTSD | P07339 | Cathepsin D | 412 | 31.07 | 9 | 9 | 39 |
| ILV2 | P07342 | Acetolactate synthase catalytic subunit, mitochondrial | 687 | 44.54 | 23 | 0 | 71 |
| RPC40 | P07703 | DNA-directed RNA polymerases I and III subunit RPAC1 | 335 | 39.70 | 11 | 0 | 28 |
| VAS1 | P07806 | Valine--tRNA ligase, mitochondrial | 1104 | 5.89 | 5 | 0 | 12 |
| CAR2 | P07991 | Ornithine aminotransferase | 424 | 3.30 | 1 | 0 | 1 |
| RPB2 | P08518 | DNA-directed RNA polymerase II subunit RPB2 | 1224 | 0.82 | 1 | 0 | 1 |
| ERG20 | P08524 | Farnesyl pyrophosphate synthase | 352 | 9.66 | 3 | 0 | 14 |
| CYR1 | P08678 | Adenylate cyclase | 2026 | 1.53 | 2 | 0 | 2 |
| SUI3 | P09064 | Eukaryotic translation initiation factor 2 subunit beta | 285 | 34.74 | 7 | 0 | 18 |
| PRB1 | P09232 | Cerevisin | 635 | 7.24 | 4 | 0 | 5 |
| ILS1 | P09436 | Isoleucine--tRNA ligase, cytoplasmic | 1072 | 6.06 | 5 | 0 | 10 |
| MIS1 | P09440 | C-1-tetrahydrofolate synthase, mitochondrial | 975 | 18.67 | 15 | 0 | 31 |
| LPD1 | P09624 | Dihydrolipoyl dehydrogenase, mitochondrial | 499 | 2.00 | 1 | 0 | 1 |
| TUB1 | P09733 | Tubulin alpha-1 chain | 447 | 18.57 | 6 | 0 | 27 |
| TUB3 | P09734 | Tubulin alpha-3 chain | 445 | 8.99 | 3 | 0 | 22 |
| CDC16 | P09798 | Anaphase-promoting complex subunit CDC16 | 840 | 3.33 | 2 | 0 | 2 |
| RNR2 | P09938 | Ribonucleoside-diphosphate reductase small chain 1 | 399 | 3.01 | 1 | 0 | 3 |

| Gene | Protein ID | Protein description | Length (aa) | Coverage (%) | Total peptides | Unique peptides | Total spectral count |
|---|---|---|---|---|---|---|---|
| RPS25B | P0C0T4 | Small ribosomal subunit protein eS25B | 108 | 21.30 | 3 | 0 | 4 |
| RPS21A | P0C0V8 | Small ribosomal subunit protein eS21A | 87 | 68.97 | 6 | 0 | 25 |
| RPS22A | P0C0W1 | Small ribosomal subunit protein uS8A | 130 | 63.85 | 7 | 0 | 19 |
| RPL11A | P0C0W9 | Large ribosomal subunit protein uL5A | 174 | 47.13 | 8 | 0 | 25 |
| RPS28B | P0C0X0 | Small ribosomal subunit protein eS28B | 67 | 41.79 | 3 | 0 | 9 |
| RPL27A | P0C2H6 | Large ribosomal subunit protein eL27A | 136 | 22.06 | 3 | 0 | 6 |
| RPL31A | P0C2H8 | Large ribosomal subunit protein eL31A | 113 | 69.91 | 9 | 0 | 36 |
| SSC1 | P0CS90 | Import motor subunit, mitochondrial | 654 | 40.98 | 24 | 0 | 67 |
| RPL20A | P0CX23 | Large ribosomal subunit protein eL20A | 172 | 43.02 | 7 | 0 | 31 |
| RPL43A | P0CX25 | Large ribosomal subunit protein eL43A | 92 | 50.00 | 4 | 0 | 9 |
| RPL42A | P0CX27 | Large ribosomal subunit protein eL42A | 106 | 7.55 | 1 | 0 | 1 |
| RPS23A | P0CX29 | Small ribosomal subunit protein uS12A | 145 | 50.34 | 7 | 0 | 20 |
| RPS24A | P0CX31 | Small ribosomal subunit protein eS24A | 135 | 53.33 | 10 | 0 | 31 |
| RPS30A | P0CX33 | Small ribosomal subunit protein eS30A | 63 | 30.16 | 3 | 0 | 4 |
| RPS4A | P0CX35 | Small ribosomal subunit protein eS4A | 261 | 53.26 | 15 | 0 | 74 |
| RPS6A | P0CX37 | Small ribosomal subunit protein eS6A | 236 | 45.34 | 11 | 0 | 39 |
| RPS8A | P0CX39 | Small ribosomal subunit protein eS8A | 200 | 46.50 | 9 | 0 | 21 |
| RPL23A | P0CX41 | Large ribosomal subunit protein uL14A | 137 | 50.36 | 6 | 0 | 42 |
| RPL1A | P0CX43 | Large ribosomal subunit protein uL1A | 217 | 38.25 | 6 | 0 | 18 |
| RPL2A | P0CX45 | Large ribosomal subunit protein uL2A | 254 | 39.76 | 10 | 0 | 43 |
| RPS11A | P0CX47 | Small ribosomal subunit protein uS17A | 156 | 55.13 | 10 | 0 | 21 |
| RPL18A | P0CX49 | Large ribosomal subunit protein eL18A | 186 | 26.34 | 5 | 0 | 15 |
| RPS16A | P0CX51 | Small ribosomal subunit protein uS9A | 143 | 53.15 | 8 | 0 | 25 |
| RPL12A | P0CX53 | Large ribosomal subunit protein uL11A | 165 | 69.09 | 12 | 0 | 46 |
| RPS18A | P0CX55 | Small ribosomal subunit protein uS13A | 146 | 58.90 | 7 | 0 | 28 |
| RPL19A | P0CX82 | Large ribosomal subunit protein eL19A | 189 | 34.39 | 8 | 0 | 33 |
| RPL35A | P0CX84 | Large ribosomal subunit protein uL29A | 120 | 36.67 | 7 | 0 | 24 |
| AMY1B | P0DTE7 | Alpha-amylase 1B | 511 | 3.13 | 1 | 0 | 1 |
| SSA1 | P10591 | Heat shock protein SSA1 | 642 | 74.92 | 41 | 0 | 277 |
| SSA2 | P10592 | Heat shock protein SSA2 | 639 | 74.96 | 44 | 0 | 296 |
| TXN | P10599 | Thioredoxin | 105 | 40.00 | 4 | 4 | 10 |
| SAM1 | P10659 | S-adenosylmethionine synthase 1 | 382 | 50.00 | 14 | 0 | 25 |
| RPL4A | P10664 | Large ribosomal subunit protein uL4A | 362 | 57.18 | 17 | 0 | 79 |
| RPA190 | P10964 | DNA-directed RNA polymerase I subunit RPA190 | 1664 | 0.66 | 1 | 0 | 1 |
| ARF1 | P11076 | ADP-ribosylation factor 1 | 181 | 13.81 | 2 | 0 | 6 |
| PYC1 | P11154 | Pyruvate carboxylase 1 | 1178 | 5.52 | 5 | 0 | 10 |
| SSB1 | P11484 | Ribosome-associated molecular chaperone SSB1 | 613 | 61.99 | 33 | 0 | 261 |
| SUP45 | P12385 | Eukaryotic peptide chain release factor subunit 1 | 437 | 52.86 | 23 | 0 | 60 |

| Gene | Protein ID | Protein description | Length (aa) | Coverage (%) | Total peptides | Unique peptides | Total spectral count |
|---|---|---|---|---|---|---|---|
| TCP1 | P12612 | T-complex protein 1 subunit alpha | 559 | 9.66 | 5 | 0 | 7 |
| MRP13 | P12686 | Small ribosomal subunit protein mS44 | 339 | 2.65 | 1 | 0 | 3 |
| YPK1 | P12688 | Serine/threonine-protein kinase YPK1 | 680 | 3.97 | 2 | 0 | 3 |
| LAT1 | P12695 | Dihydrolipoyllysine-residue acetyltransferase component of pyruvate dehydrogenase complex, mitochondrial | 482 | 37.34 | 14 | 0 | 65 |
| PGI1 | P12709 | Glucose-6-phosphate isomerase | 554 | 38.99 | 15 | 0 | 34 |
| KRT10 | P13645 | Keratin, type I cytoskeletal 10 | 584 | 67.64 | 40 | 34 | 607 |
| HOM2 | P13663 | Aspartate-semialdehyde dehydrogenase | 365 | 13.97 | 3 | 0 | 7 |
| DPM1 | P14020 | Dolichol-phosphate mannosyltransferase | 267 | 4.87 | 1 | 0 | 5 |
| RPL30 | P14120 | Large ribosomal subunit protein eL30 | 105 | 56.19 | 4 | 0 | 13 |
| RPL3 | P14126 | Large ribosomal subunit protein uL3 | 387 | 39.79 | 16 | 0 | 72 |
| FBA1 | P14540 | Fructose-bisphosphate aldolase | 359 | 45.68 | 12 | 0 | 93 |
| GFA1 | P14742 | Glutamine--fructose-6-phosphate aminotransferase [isomerizing] | 717 | 4.60 | 2 | 0 | 5 |
| NMT1 | P14743 | Glycylpeptide N-tetradecanoyltransferase | 455 | 2.42 | 1 | 0 | 4 |
| CMP2 | P14747 | Serine/threonine-protein phosphatase 2B catalytic subunit A2 | 604 | 13.41 | 7 | 0 | 22 |
| CPR1 | P14832 | Peptidyl-prolyl cis-trans isomerase | 162 | 56.17 | 8 | 0 | 47 |
| APE1 | P14904 | Vacuolar aminopeptidase 1 | 514 | 7.78 | 3 | 0 | 8 |
| TAL1 | P15019 | Transaldolase | 335 | 48.66 | 14 | 0 | 31 |
| HSC82 | P15108 | ATP-dependent molecular chaperone HSC82 | 705 | 41.99 | 31 | 0 | 85 |
| KRS1 | P15180 | Lysine--tRNA ligase, cytoplasmic | 591 | 3.72 | 2 | 0 | 6 |
| | P15252 | Rubber elongation factor protein | 138 | 15.94 | 2 | 2 | 3 |
| AMD1 | P15274 | AMP deaminase | 810 | 2.35 | 2 | 0 | 2 |
| GUK1 | P15454 | Guanylate kinase | 187 | 35.29 | 6 | 0 | 12 |
| TRM1 | P15565 | tRNA (guanine(26)-N(2))-dimethyltransferase, mitochondrial | 570 | 5.79 | 3 | 0 | 4 |
| FRS1 | P15624 | Phenylalanine--tRNA ligase beta subunit | 595 | 12.94 | 6 | 0 | 10 |
| FRS2 | P15625 | Phenylalanine--tRNA ligase alpha subunit | 503 | 5.37 | 2 | 0 | 3 |
| NOP1 | P15646 | rRNA 2'-O-methyltransferase fibrillarin | 327 | 20.18 | 5 | 0 | 11 |
| BGL2 | P15703 | Glucan 1,3-beta-glucosidase | 313 | 16.61 | 4 | 0 | 6 |
| STI1 | P15705 | Heat shock protein STI1 | 589 | 17.32 | 9 | 0 | 14 |
| ABP1 | P15891 | Actin-binding protein | 592 | 4.73 | 2 | 0 | 4 |
| HSP26 | P15992 | Heat shock protein 26 | 214 | 47.20 | 8 | 0 | 24 |
| THR4 | P16120 | Threonine synthase | 514 | 18.09 | 8 | 0 | 11 |
| VMA2 | P16140 | V-type proton ATPase subunit B | 517 | 48.74 | 20 | 0 | 53 |
| PDA1 | P16387 | Pyruvate dehydrogenase E1 component subunit alpha, mitochondrial | 420 | 38.57 | 15 | 0 | 42 |
| KAR2 | P16474 | Endoplasmic reticulum chaperone BiP | 682 | 29.62 | 19 | 0 | 42 |
| YEF3 | P16521 | Elongation factor 3A | 1044 | 31.51 | 29 | 0 | 79 |
| CDC23 | P16522 | Anaphase-promoting complex subunit CDC23 | 626 | 2.24 | 1 | 0 | 1 |
| OM45 | P16547 | Mitochondrial outer membrane protein OM45 | 393 | 16.28 | 6 | 0 | 13 |

| Gene | Protein ID | Protein description | Length (aa) | Coverage (%) | Total peptides | Unique peptides | Total spectral count |
|---|---|---|---|---|---|---|---|
| APA1 | P16550 | Protein APA1 | 321 | 9.03 | 3 | 0 | 3 |
| NCP1 | P16603 | NADPH--cytochrome P450 reductase | 691 | 7.24 | 4 | 0 | 7 |
| PFK1 | P16861 | ATP-dependent 6-phosphofructokinase subunit alpha | 987 | 27.05 | 19 | 0 | 43 |
| VMA1 | P17255 | V-type proton ATPase catalytic subunit A | 1071 | 24.84 | 23 | 0 | 52 |
| THR1 | P17423 | Homoserine kinase | 357 | 10.64 | 3 | 0 | 9 |
| MDH1 | P17505 | Malate dehydrogenase, mitochondrial | 334 | 33.23 | 6 | 0 | 22 |
| TPM1 | P17536 | Tropomyosin-1 | 199 | 13.57 | 3 | 0 | 5 |
| SRV2 | P17555 | Adenylyl cyclase-associated protein | 526 | 29.66 | 10 | 0 | 27 |
| GRX2 | P17695 | Glutaredoxin-2 | 143 | 16.78 | 2 | 0 | 6 |
| GLK1 | P17709 | Glucokinase-1 | 500 | 14.60 | 6 | 0 | 15 |
| CLC1 | P17891 | Clathrin light chain | 233 | 44.21 | 10 | 0 | 21 |
| PDI1 | P17967 | Protein disulfide-isomerase | 522 | 26.44 | 10 | 0 | 28 |
| PET9 | P18239 | ADP,ATP carrier protein 2 | 318 | 19.50 | 6 | 0 | 18 |
| SNF5 | P18480 | SWI/SNF chromatin-remodeling complex subunit SNF5 | 905 | 1.44 | 1 | 0 | 1 |
| FUR1 | P18562 | Uracil phosphoribosyltransferase | 216 | 3.70 | 1 | 0 | 1 |
| COQ1 | P18900 | Hexaprenyl pyrophosphate synthase, mitochondrial | 473 | 7.61 | 3 | 0 | 4 |
| FAS2 | P19097 | Fatty acid synthase subunit alpha | 1887 | 15.53 | 23 | 0 | 51 |
| ARF2 | P19146 | ADP-ribosylation factor 2 | 181 | 13.81 | 2 | 0 | 6 |
| ACO1 | P19414 | Aconitate hydratase, mitochondrial | 778 | 14.65 | 10 | 0 | 15 |
| MYO2 | P19524 | Myosin-2 | 1574 | 4.76 | 5 | 0 | 9 |
| HSP60 | P19882 | Heat shock protein 60, mitochondrial | 572 | 59.44 | 33 | 0 | 120 |
| FPR1 | P20081 | FK506-binding protein 1 | 114 | 48.25 | 5 | 0 | 13 |
| SUI2 | P20459 | Eukaryotic translation initiation factor 2 subunit alpha | 304 | 5.26 | 1 | 0 | 3 |
| SAR1 | P20606 | Small COPII coat GTPase SAR1 | 190 | 24.74 | 4 | 0 | 7 |
| ADE2 | P21264 | Phosphoribosylaminoimidazole carboxylase | 571 | 3.68 | 2 | 0 | 2 |
| ATP15 | P21306 | ATP synthase subunit epsilon, mitochondrial | 62 | 38.71 | 2 | 0 | 5 |
| RNR1 | P21524 | Ribonucleoside-diphosphate reductase large chain 1 | 888 | 4.50 | 3 | 0 | 3 |
| REB1 | P21538 | DNA-binding protein REB1 | 810 | 11.73 | 7 | 0 | 16 |
| VPS1 | P21576 | Vacuolar protein sorting-associated protein 1 | 704 | 4.40 | 3 | 0 | 8 |
| IDP1 | P21954 | Isocitrate dehydrogenase [NADP], mitochondrial | 428 | 14.49 | 3 | 0 | 6 |
| MCK1 | P21965 | Protein kinase MCK1 | 375 | 16.80 | 6 | 0 | 13 |
| SNF2 | P22082 | Transcription regulatory protein SNF2 | 1703 | 2.00 | 3 | 0 | 3 |
| CHC1 | P22137 | Clathrin heavy chain | 1653 | 4.72 | 7 | 0 | 12 |
| SSA4 | P22202 | Heat shock protein SSA4 | 642 | 25.70 | 17 | 0 | 99 |
| VMA4 | P22203 | V-type proton ATPase subunit E | 233 | 68.24 | 16 | 0 | 74 |
| TRX1 | P22217 | Thioredoxin-1 | 103 | 66.02 | 6 | 0 | 11 |
| SAN1 | P22470 | Protein SAN1 | 610 | 10.66 | 4 | 0 | 19 |
| UBA1 | P22515 | Ubiquitin-activating enzyme E1 1 | 1024 | 4.30 | 4 | 0 | 6 |

| Gene | Protein ID | Protein description | Length (aa) | Coverage (%) | Total peptides | Unique peptides | Total spectral count |
|---|---|---|---|---|---|---|---|
| CMK2 | P22517 | Calcium/calmodulin-dependent protein kinase II | 447 | 5.82 | 2 | 0 | 3 |
| SIN3 | P22579 | Transcriptional regulatory protein SIN3 | 1536 | 0.91 | 1 | 0 | 1 |
| TRX2 | P22803 | Thioredoxin-2 | 104 | 66.35 | 6 | 0 | 22 |
| HSP12 | P22943 | 12 kDa heat shock protein | 109 | 21.10 | 2 | 0 | 4 |
| TKL1 | P23254 | Transketolase 1 | 680 | 25.15 | 14 | 0 | 27 |
| CNA1 | P23287 | Serine/threonine-protein phosphatase 2B catalytic subunit A1 | 553 | 11.57 | 6 | 0 | 13 |
| HYP2 | P23301 | Eukaryotic translation initiation factor 5A-1 | 157 | 11.46 | 1 | 0 | 1 |
| GSY1 | P23337 | Glycogen [starch] synthase isoform 1 | 708 | 2.40 | 2 | 0 | 3 |
| RPL24B | P24000 | Large ribosomal subunit protein eL24B | 155 | 33.55 | 10 | 0 | 45 |
| PHO3 | P24031 | Constitutive acid phosphatase | 467 | 3.43 | 1 | 0 | 1 |
| SSD1 | P24276 | Protein SSD1 | 1250 | 34.16 | 30 | 0 | 112 |
| ERG6 | P25087 | Sterol 24-C-methyltransferase ERG6 | 383 | 14.36 | 4 | 0 | 7 |
| NAP1 | P25293 | Nucleosome assembly protein | 417 | 10.79 | 3 | 0 | 8 |
| SIS1 | P25294 | Protein SIS1 | 352 | 12.22 | 3 | 0 | 11 |
| CNB1 | P25296 | Calcineurin subunit B | 175 | 26.86 | 3 | 0 | 4 |
| CDC10 | P25342 | Cell division control protein 10 | 322 | 21.12 | 5 | 0 | 17 |
| RVS161 | P25343 | Reduced viability upon starvation protein 161 | 265 | 33.96 | 8 | 0 | 29 |
| HCM1 | P25364 | Forkhead transcription factor HCM1 | 564 | 3.19 | 1 | 0 | 1 |
| GRX1 | P25373 | Glutaredoxin-1 | 110 | 14.55 | 1 | 0 | 1 |
| RPS2 | P25443 | Small ribosomal subunit protein uS5 | 254 | 38.19 | 10 | 0 | 37 |
| YDJ1 | P25491 | Mitochondrial protein import protein MAS5 | 409 | 10.02 | 3 | 0 | 11 |
| BUD3 | P25558 | Bud site selection protein 3 | 1636 | 1.53 | 2 | 0 | 2 |
| SRO9 | P25567 | RNA-binding protein SRO9 | 434 | 42.40 | 12 | 0 | 42 |
| YCL049C | P25577 | Uncharacterized protein YCL049C | 312 | 2.88 | 1 | 0 | 2 |
| IMG1 | P25626 | Large ribosomal subunit protein bL19m | 169 | 13.02 | 2 | 0 | 5 |
| PAT1 | P25644 | DNA topoisomerase 2-associated protein PAT1 | 796 | 2.89 | 2 | 0 | 2 |
| CDC48 | P25694 | Cell division control protein 48 | 835 | 13.17 | 8 | 0 | 24 |
| CPR3 | P25719 | Peptidyl-prolyl cis-trans isomerase C, mitochondrial | 182 | 7.14 | 1 | 0 | 2 |
| RPL5 | P26321 | Large ribosomal subunit protein uL18 | 297 | 33.00 | 6 | 0 | 30 |
| CDC60 | P26637 | Leucine--tRNA ligase, cytoplasmic | 1090 | 16.79 | 14 | 0 | 44 |
| RPS5 | P26783 | Small ribosomal subunit protein uS7 | 225 | 42.67 | 10 | 0 | 34 |
| RPL16A | P26784 | Large ribosomal subunit protein uL13A | 199 | 31.66 | 6 | 0 | 12 |
| RPL16B | P26785 | Large ribosomal subunit protein uL13B | 198 | 32.83 | 7 | 0 | 16 |
| RPS7A | P26786 | Small ribosomal subunit protein eS7A | 190 | 67.89 | 13 | 0 | 72 |
| CMK1 | P27466 | Calcium/calmodulin-dependent protein kinase I | 446 | 10.99 | 4 | 0 | 11 |
| NSR1 | P27476 | Nuclear localization sequence-binding protein | 414 | 9.66 | 3 | 0 | 3 |
| ADE1 | P27616 | Phosphoribosylaminoimidazole-succinocarboxamide synthase | 306 | 34.64 | 8 | 0 | 25 |
| SPT5 | P27692 | Transcription elongation factor SPT5 | 1063 | 47.32 | 40 | 0 | 138 |

| Gene | Protein ID | Protein description | Length (aa) | Coverage (%) | Total peptides | Unique peptides | Total spectral count |
|---|---|---|---|---|---|---|---|
| KRE2 | P27809 | Glycolipid 2-alpha-mannosyltransferase | 442 | 14.93 | 5 | 0 | 15 |
| IDH2 | P28241 | Isocitrate dehydrogenase [NAD] subunit 2, mitochondrial | 369 | 23.31 | 6 | 0 | 17 |
| OXP1 | P28273 | 5-oxoprolinase | 1286 | 3.42 | 3 | 0 | 4 |
| URA7 | P28274 | CTP synthase 1 | 579 | 4.84 | 2 | 0 | 2 |
| CAP1 | P28495 | F-actin-capping protein subunit alpha | 268 | 10.82 | 3 | 0 | 5 |
| ARO2 | P28777 | Chorismate synthase | 376 | 7.18 | 2 | 0 | 5 |
| IDH1 | P28834 | Isocitrate dehydrogenase [NAD] subunit 1, mitochondrial | 360 | 32.50 | 9 | 0 | 43 |
| BMH1 | P29311 | Protein BMH1 | 267 | 44.94 | 10 | 0 | 54 |
| RPL8B | P29453 | Large ribosomal subunit protein eL8B | 256 | 59.77 | 18 | 0 | 35 |
| SEC65 | P29478 | Signal recognition particle subunit SEC65 | 273 | 6.23 | 2 | 0 | 6 |
| TRR1 | P29509 | Thioredoxin reductase 1 | 319 | 40.44 | 7 | 0 | 11 |
| CAM1 | P29547 | Elongation factor 1-gamma 1 | 415 | 7.47 | 3 | 0 | 10 |
| PMI40 | P29952 | Mannose-6-phosphate isomerase | 429 | 6.29 | 2 | 0 | 3 |
| CRM1 | P30822 | Exportin-1 | 1084 | 2.12 | 2 | 0 | 4 |
| ATP7 | P30902 | ATP synthase subunit d, mitochondrial | 174 | 12.07 | 2 | 0 | 2 |
| HOM6 | P31116 | Homoserine dehydrogenase | 359 | 33.70 | 9 | 0 | 27 |
| CYS3 | P31373 | Cystathionine gamma-lyase | 394 | 19.29 | 6 | 0 | 18 |
| VMA5 | P31412 | V-type proton ATPase subunit C | 392 | 14.54 | 4 | 0 | 8 |
| HSP104 | P31539 | Heat shock protein 104 | 908 | 17.51 | 13 | 0 | 21 |
| TPS2 | P31688 | Trehalose-phosphatase | 896 | 18.53 | 12 | 0 | 32 |
| ACB1 | P31787 | Acyl-CoA-binding protein | 87 | 48.28 | 3 | 0 | 10 |
| SEC21 | P32074 | Coatomer subunit gamma | 935 | 3.96 | 3 | 0 | 5 |
| GLN1 | P32288 | Glutamine synthetase | 370 | 13.51 | 5 | 0 | 9 |
| EFT1 | P32324 | Elongation factor 2 | 842 | 50.36 | 35 | 0 | 104 |
| PSE1 | P32337 | Importin subunit beta-3 | 1089 | 1.01 | 1 | 0 | 1 |
| NTH1 | P32356 | Cytosolic neutral trehalase | 751 | 4.79 | 3 | 0 | 6 |
| MVD1 | P32377 | Diphosphomevalonate decarboxylase | 396 | 12.88 | 3 | 0 | 5 |
| MDH3 | P32419 | Malate dehydrogenase, peroxisomal | 343 | 14.58 | 3 | 0 | 6 |
| RIM1 | P32445 | Single-stranded DNA-binding protein RIM1, mitochondrial | 135 | 28.89 | 3 | 0 | 5 |
| ARO4 | P32449 | Phospho-2-dehydro-3-deoxyheptonate aldolase, tyrosine-inhibited | 370 | 17.57 | 4 | 0 | 8 |
| CDC3 | P32457 | Cell division control protein 3 | 520 | 24.81 | 12 | 0 | 25 |
| CDC12 | P32468 | Cell division control protein 12 | 407 | 23.34 | 7 | 0 | 18 |
| EFB1 | P32471 | Elongation factor 1-beta | 206 | 33.50 | 5 | 0 | 14 |
| PDB1 | P32473 | Pyruvate dehydrogenase E1 component subunit beta, mitochondrial | 366 | 33.33 | 8 | 0 | 27 |
| GCD11 | P32481 | Eukaryotic translation initiation factor 2 subunit gamma | 527 | 15.75 | 6 | 0 | 14 |
| NHP2 | P32495 | H/ACA ribonucleoprotein complex subunit NHP2 | 156 | 9.62 | 2 | 0 | 2 |
| NUP2 | P32499 | Nucleoporin NUP2 | 720 | 39.44 | 20 | 0 | 52 |
| gag | P32503 | Major capsid protein | 680 | 16.32 | 9 | 8 | 42 |

| Gene | Protein ID | Protein description | Length (aa) | Coverage (%) | Total peptides | Unique peptides | Total spectral count |
|---|---|---|---|---|---|---|---|
| PAN1 | P32521 | Actin cytoskeleton-regulatory complex protein PAN1 | 1480 | 6.15 | 7 | 0 | 20 |
| ZUO1 | P32527 | Zuotin | 433 | 14.78 | 5 | 0 | 12 |
| RPN2 | P32565 | 26S proteasome regulatory subunit RPN2 | 945 | 1.16 | 1 | 0 | 1 |
| CYS4 | P32582 | Cystathionine beta-synthase | 507 | 71.79 | 28 | 0 | 92 |
| SSE1 | P32589 | Heat shock protein homolog SSE1 | 693 | 56.28 | 35 | 0 | 159 |
| SWI3 | P32591 | SWI/SNF complex subunit SWI3 | 825 | 11.39 | 8 | 0 | 19 |
| GLC7 | P32598 | Serine/threonine-protein phosphatase PP1-2 | 312 | 9.29 | 2 | 0 | 6 |
| SAC6 | P32599 | Fimbrin | 642 | 2.02 | 1 | 0 | 2 |
| RTG2 | P32608 | Retrograde regulation protein 2 | 588 | 3.74 | 2 | 0 | 2 |
| VMA8 | P32610 | V-type proton ATPase subunit D | 256 | 3.91 | 1 | 0 | 1 |
| FRD1 | P32614 | Fumarate reductase 1 | 470 | 7.45 | 3 | 0 | 8 |
| NRP1 | P32770 | Putative RNA-binding protein involved in heterochromatin assembly | 719 | 1.67 | 1 | 0 | 3 |
| GSP1 | P32835 | GTP-binding nuclear protein GSP1/CNR1 | 219 | 36.53 | 7 | 0 | 13 |
| UGP1 | P32861 | UTP--glucose-1-phosphate uridylyltransferase | 499 | 17.03 | 7 | 0 | 13 |
| PRS1 | P32895 | Ribose-phosphate pyrophosphokinase 1 | 427 | 10.30 | 3 | 0 | 9 |
| IMP3 | P32899 | U3 small nucleolar ribonucleoprotein protein IMP3 | 183 | 8.20 | 1 | 0 | 1 |
| SPT4 | P32914 | Transcription elongation factor SPT4 | 102 | 30.39 | 2 | 0 | 6 |
| RPT5 | P33297 | 26S proteasome regulatory subunit 6A | 434 | 12.90 | 4 | 0 | 9 |
| RPT3 | P33298 | 26S proteasome regulatory subunit 6B homolog | 428 | 10.51 | 3 | 0 | 13 |
| RPT1 | P33299 | 26S proteasome regulatory subunit 7 homolog | 467 | 4.28 | 2 | 0 | 2 |
| SER1 | P33330 | Phosphoserine aminotransferase | 395 | 5.06 | 1 | 0 | 8 |
| NTF2 | P33331 | Nuclear transport factor 2 | 125 | 25.60 | 3 | 0 | 12 |
| PGM1 | P33401 | Phosphoglucomutase 1 | 570 | 6.49 | 3 | 0 | 8 |
| GCN1 | P33892 | eIF-2-alpha kinase activator GCN1 | 2672 | 1.53 | 3 | 0 | 3 |
| BMH2 | P34730 | Protein BMH2 | 273 | 40.29 | 9 | 0 | 48 |
| FAB1 | P34756 | 1-phosphatidylinositol 3-phosphate 5-kinase FAB1 | 2278 | 0.35 | 1 | 0 | 1 |
| TSA1 | P34760 | Peroxiredoxin TSA1 | 196 | 61.73 | 11 | 0 | 50 |
| RRP1 | P35178 | Ribosomal RNA-processing protein 1 | 278 | 14.03 | 3 | 0 | 7 |
| SQT1 | P35184 | Ribosome assembly protein SQT1 | 431 | 20.65 | 7 | 0 | 30 |
| KRT9 | P35527 | Keratin, type I cytoskeletal 9 | 623 | 64.53 | 34 | 33 | 538 |
| TMA19 | P35691 | Translationally-controlled tumor protein homolog | 167 | 38.92 | 7 | 0 | 21 |
| DEF1 | P35732 | RNA polymerase II degradation factor 1 | 738 | 21.54 | 12 | 0 | 37 |
| KES1 | P35844 | Oxysterol-binding protein homolog 4 | 434 | 14.75 | 5 | 0 | 18 |
| SWH1 | P35845 | Oxysterol-binding protein homolog 1 | 1188 | 5.72 | 5 | 0 | 9 |
| KRT2 | P35908 | Keratin, type II cytoskeletal 2 epidermal | 639 | 88.26 | 57 | 51 | 531 |
| RPS27A | P35997 | Small ribosomal subunit protein eS27A | 82 | 37.80 | 3 | 0 | 10 |
| TEF4 | P36008 | Elongation factor 1-gamma 2 | 412 | 33.98 | 13 | 0 | 33 |
| YNK1 | P36010 | Nucleoside diphosphate kinase | 153 | 18.95 | 2 | 0 | 5 |

| Gene | Protein ID | Protein description | Length (aa) | Coverage (%) | Total peptides | Unique peptides | Total spectral count |
|---|---|---|---|---|---|---|---|
| MAE1 | P36013 | NAD-dependent malic enzyme, mitochondrial | 669 | 4.78 | 2 | 0 | 3 |
| EAP1 | P36041 | Protein EAP1 | 632 | 11.08 | 5 | 0 | 19 |
| RPL14A | P36105 | Large ribosomal subunit protein eL14A | 138 | 39.13 | 5 | 0 | 8 |
| SPC34 | P36131 | DASH complex subunit SPC34 | 295 | 3.05 | 1 | 0 | 4 |
| PXL1 | P36166 | Paxillin-like protein 1 | 706 | 1.70 | 1 | 0 | 1 |
| JNM1 | P36224 | Nuclear migration protein JNM1 | 373 | 10.19 | 3 | 0 | 7 |
| TYS1 | P36421 | Tyrosine--tRNA ligase, cytoplasmic | 394 | 2.03 | 1 | 0 | 1 |
| MRPL3 | P36516 | Large ribosomal subunit protein mL44 | 390 | 3.59 | 1 | 0 | 1 |
| MRPL11 | P36521 | Large ribosomal subunit protein uL10m | 249 | 8.84 | 2 | 0 | 5 |
| PIM1 | P36775 | Lon protease homolog, mitochondrial | 1133 | 3.80 | 3 | 0 | 4 |
| SHM2 | P37291 | Serine hydroxymethyltransferase, cytosolic | 469 | 38.38 | 17 | 0 | 36 |
| SHM1 | P37292 | Serine hydroxymethyltransferase, mitochondrial | 490 | 10.41 | 5 | 0 | 7 |
| APE3 | P37302 | Aminopeptidase Y | 537 | 6.33 | 2 | 0 | 13 |
| GLY1 | P37303 | Low specificity L-threonine aldolase | 387 | 3.88 | 1 | 0 | 1 |
| ADE17 | P38009 | Bifunctional purine biosynthesis protein ADE17 | 592 | 21.11 | 8 | 0 | 15 |
| ASC1 | P38011 | Small ribosomal subunit protein RACK1 | 319 | 43.89 | 11 | 0 | 39 |
| AHP1 | P38013 | Peroxiredoxin AHP1 | 176 | 48.86 | 8 | 0 | 30 |
| CDC27 | P38042 | Anaphase-promoting complex subunit CDC27 | 758 | 3.43 | 2 | 0 | 2 |
| RPL32 | P38061 | Large ribosomal subunit protein eL32 | 130 | 42.31 | 6 | 0 | 17 |
| ATP3 | P38077 | ATP synthase subunit gamma, mitochondrial | 311 | 44.05 | 12 | 0 | 35 |
| GRS1 | P38088 | Glycine--tRNA ligase 1, mitochondrial | 690 | 15.07 | 8 | 0 | 12 |
| ATG42 | P38109 | Vacuolar serine-type carboxypeptidase ATG42 | 508 | 5.12 | 3 | 0 | 4 |
| ARA1 | P38115 | D-arabinose dehydrogenase [NAD(P)+] heavy chain | 344 | 4.07 | 1 | 0 | 1 |
| TAF5 | P38129 | Transcription initiation factor TFIID subunit 5 | 798 | 5.26 | 3 | 0 | 6 |
| OLA1 | P38219 | Obg-like ATPase 1 | 394 | 34.01 | 11 | 0 | 19 |
| RPG1 | P38249 | Eukaryotic translation initiation factor 3 subunit A | 964 | 18.67 | 15 | 0 | 37 |
| SIF2 | P38262 | SIR4-interacting protein SIF2 | 535 | 2.06 | 1 | 0 | 1 |
| MCX1 | P38323 | ATP-dependent clpX-like chaperone, mitochondrial | 520 | 5.00 | 2 | 0 | 2 |
| ENP1 | P38333 | Essential nuclear protein 1 | 483 | 6.63 | 2 | 0 | 3 |
| REI1 | P38344 | Cytoplasmic 60S subunit biogenesis factor REI1 | 393 | 2.80 | 1 | 0 | 8 |
| TPS3 | P38426 | Trehalose synthase complex regulatory subunit TPS3 | 1054 | 5.03 | 4 | 0 | 5 |
| TSL1 | P38427 | Trehalose synthase complex regulatory subunit TSL1 | 1098 | 13.93 | 12 | 1 | 30 |
| GUA1 | P38625 | GMP synthase [glutamine-hydrolyzing] | 525 | 8.57 | 4 | 0 | 9 |
| GLO3 | P38682 | ADP-ribosylation factor GTPase-activating protein GLO3 | 493 | 2.03 | 1 | 0 | 1 |
| KSP1 | P38691 | Serine/threonine-protein kinase KSP1 | 1029 | 2.92 | 2 | 0 | 2 |
| MSC7 | P38694 | Putative aldehyde dehydrogenase-like protein YHR039C | 644 | 1.86 | 1 | 0 | 2 |
| RPS20 | P38701 | Small ribosomal subunit protein uS10 | 121 | 45.45 | 6 | 0 | 12 |
| DED81 | P38707 | Asparagine--tRNA ligase, cytoplasmic | 554 | 10.29 | 4 | 0 | 7 |

| Gene | Protein ID | Protein description | Length (aa) | Coverage (%) | Total peptides | Unique peptides | Total spectral count |
|---|---|---|---|---|---|---|---|
| YHR020W | P38708 | Putative proline--tRNA ligase YHR020W | 688 | 12.21 | 6 | 0 | 9 |
| GND1 | P38720 | 6-phosphogluconate dehydrogenase, decarboxylating 1 | 489 | 19.22 | 8 | 0 | 24 |
| OSH7 | P38755 | Oxysterol-binding protein homolog 7 | 437 | 12.13 | 5 | 0 | 10 |
| RPN1 | P38764 | 26S proteasome regulatory subunit RPN1 | 993 | 3.12 | 2 | 0 | 3 |
| SSZ1 | P38788 | Ribosome-associated complex subunit SSZ1 | 538 | 35.87 | 13 | 0 | 28 |
| DYS1 | P38791 | Deoxyhypusine synthase | 387 | 11.89 | 3 | 0 | 9 |
| RTC3 | P38804 | Restriction of telomere capping protein 3 | 111 | 18.02 | 2 | 0 | 4 |
| TRA1 | P38811 | Transcription-associated protein 1 | 3744 | 0.19 | 1 | 0 | 1 |
| BZZ1 | P38822 | Protein BZZ1 | 633 | 7.11 | 3 | 0 | 9 |
| ARO9 | P38840 | Aromatic amino acid aminotransferase 2 | 513 | 6.82 | 2 | 0 | 2 |
| RPN10 | P38886 | 26S proteasome regulatory subunit RPN10 | 268 | 4.10 | 1 | 0 | 2 |
| BAT1 | P38891 | Branched-chain-amino-acid aminotransferase, mitochondrial | 393 | 11.70 | 4 | 0 | 7 |
| HSP10 | P38910 | 10 kDa heat shock protein, mitochondrial | 106 | 53.77 | 4 | 0 | 9 |
| TIF11 | P38912 | Eukaryotic translation initiation factor 1A | 153 | 14.38 | 2 | 0 | 3 |
| BFR1 | P38934 | Nuclear segregation protein BFR1 | 470 | 12.34 | 5 | 0 | 8 |
| SEC31 | P38968 | Protein transport protein SEC31 | 1273 | 5.34 | 5 | 0 | 11 |
| ADE6 | P38972 | Phosphoribosylformylglycinamidine synthase | 1358 | 4.49 | 5 | 0 | 11 |
| HXT6 | P39003 | High-affinity hexose transporter HXT6 | 570 | 2.46 | 1 | 0 | 2 |
| STM1 | P39015 | Suppressor protein STM1 | 273 | 41.76 | 8 | 0 | 23 |
| CCT2 | P39076 | T-complex protein 1 subunit beta | 527 | 25.43 | 9 | 0 | 25 |
| CCT4 | P39078 | T-complex protein 1 subunit delta | 528 | 3.03 | 1 | 0 | 3 |
| CCT6 | P39079 | T-complex protein 1 subunit zeta | 546 | 3.66 | 1 | 0 | 2 |
| CAJ1 | P39101 | Protein CAJ1 | 391 | 13.30 | 4 | 0 | 9 |
| PIK1 | P39104 | Phosphatidylinositol 4-kinase PIK1 | 1066 | 1.41 | 1 | 0 | 4 |
| ILV3 | P39522 | Dihydroxy-acid dehydratase, mitochondrial | 585 | 16.92 | 8 | 0 | 24 |
| ACO2 | P39533 | Homocitrate dehydratase, mitochondrial | 789 | 1.65 | 1 | 0 | 3 |
| YHB1 | P39676 | Flavohemoprotein | 399 | 17.04 | 5 | 0 | 11 |
| NUP60 | P39705 | Nucleoporin NUP60 | 539 | 26.53 | 9 | 0 | 20 |
| ECM1 | P39715 | Shuttling pre-60S factor ECM1 | 212 | 8.96 | 2 | 0 | 6 |
| RVS167 | P39743 | Reduced viability upon starvation protein 167 | 482 | 27.80 | 9 | 0 | 45 |
| TIF4631 | P39935 | Eukaryotic initiation factor 4F subunit p150 | 952 | 21.43 | 17 | 0 | 39 |
| TIF4632 | P39936 | Eukaryotic initiation factor 4F subunit p130 | 914 | 10.28 | 8 | 0 | 14 |
| RPS26A | P39938 | Small ribosomal subunit protein eS26A | 119 | 42.86 | 5 | 0 | 25 |
| SAH1 | P39954 | Adenosylhomocysteinase | 449 | 21.83 | 9 | 0 | 15 |
| BEM2 | P39960 | GTPase-activating protein BEM2/IPL2 | 2167 | 0.42 | 1 | 0 | 1 |
| DLD3 | P39976 | D-2-hydroxyglutarate--pyruvate transhydrogenase DLD3 | 496 | 9.07 | 4 | 0 | 9 |
| NUG1 | P40010 | Nuclear GTP-binding protein NUG1 | 520 | 29.81 | 11 | 0 | 47 |
| ARB1 | P40024 | ABC transporter ATP-binding protein ARB1 | 610 | 7.38 | 3 | 0 | 5 |

| Gene | Protein ID | Protein description | Length (aa) | Coverage (%) | Total peptides | Unique peptides | Total spectral count |
|---|---|---|---|---|---|---|---|
| HMF1 | P40037 | Protein HMF1 | 129 | 23.26 | 2 | 0 | 10 |
| KAP123 | P40069 | Importin subunit beta-4 | 1113 | 11.86 | 11 | 0 | 29 |
| BUR6 | P40096 | Negative cofactor 2 complex subunit alpha | 142 | 7.75 | 1 | 0 | 2 |
| GPP2 | P40106 | Glycerol-1-phosphate phosphohydrolase 2 | 250 | 29.60 | 7 | 0 | 20 |
| SSB2 | P40150 | Ribosome-associated molecular chaperone SSB2 | 613 | 61.99 | 33 | 0 | 255 |
| YNL208W | P40159 | Uncharacterized protein YNL208W | 199 | 17.09 | 2 | 0 | 8 |
| RIO2 | P40160 | Serine/threonine-protein kinase RIO2 | 425 | 3.53 | 1 | 0 | 1 |
| MMF1 | P40185 | Protein MMF1, mitochondrial | 145 | 40.00 | 5 | 0 | 25 |
| CCS1 | P40202 | Superoxide dismutase 1 copper chaperone | 249 | 13.25 | 2 | 0 | 2 |
| RPL13B | P40212 | Large ribosomal subunit protein eL13B | 199 | 44.22 | 8 | 0 | 43 |
| PRE6 | P40303 | Proteasome subunit alpha type-4 | 254 | 9.84 | 2 | 0 | 2 |
| RPT2 | P40327 | 26S proteasome regulatory subunit 4 homolog | 437 | 7.09 | 2 | 0 | 6 |
| SEC9 | P40357 | Protein transport protein SEC9 | 651 | 2.00 | 1 | 0 | 3 |
| LYS12 | P40495 | Homoisocitrate dehydrogenase, mitochondrial | 371 | 19.14 | 6 | 0 | 11 |
| RPL34B | P40525 | Large ribosomal subunit protein eL34B | 121 | 12.40 | 2 | 0 | 8 |
| GVP36 | P40531 | Protein GVP36 | 326 | 23.31 | 7 | 0 | 13 |
| SGN1 | P40561 | RNA-binding protein SGN1 | 250 | 10.00 | 2 | 0 | 6 |
| HYR1 | P40581 | Glutathione peroxidase-like peroxiredoxin HYR1 | 163 | 4.29 | 1 | 0 | 1 |
| ALA1 | P40825 | Alanine--tRNA ligase, mitochondrial | 983 | 5.29 | 5 | 0 | 6 |
| PHB1 | P40961 | Prohibitin-1 | 287 | 12.89 | 3 | 0 | 6 |
| RPL33B | P41056 | Large ribosomal subunit protein eL33B | 107 | 39.25 | 5 | 0 | 16 |
| RPS29A | P41057 | Small ribosomal subunit protein uS14A | 56 | 26.79 | 2 | 0 | 6 |
| GPP1 | P41277 | Glycerol-1-phosphate phosphohydrolase 1 | 250 | 54.00 | 14 | 0 | 46 |
| ERG10 | P41338 | Acetyl-CoA acetyltransferase | 398 | 18.84 | 6 | 0 | 10 |
| BUD6 | P41697 | Bud site selection protein 6 | 788 | 8.50 | 5 | 0 | 12 |
| RPL10 | P41805 | Large ribosomal subunit protein uL16 | 221 | 42.99 | 13 | 0 | 46 |
| VMA13 | P41807 | V-type proton ATPase subunit H | 478 | 6.49 | 2 | 0 | 2 |
| SEC26 | P41810 | Coatomer subunit beta | 973 | 2.26 | 2 | 0 | 3 |
| SEC27 | P41811 | Coatomer subunit beta' | 889 | 5.51 | 4 | 0 | 6 |
| TFG1 | P41895 | Transcription initiation factor IIF subunit alpha | 735 | 10.88 | 7 | 0 | 11 |
| TFG2 | P41896 | Transcription initiation factor IIF subunit beta | 400 | 8.00 | 2 | 0 | 4 |
| YRB1 | P41920 | Ran-specific GTPase-activating protein 1 | 201 | 14.93 | 2 | 0 | 3 |
| PSA1 | P41940 | Mannose-1-phosphate guanyltransferase | 361 | 33.52 | 11 | 0 | 23 |
| CIR1 | P42940 | Probable electron transfer flavoprotein subunit beta | 261 | 36.02 | 9 | 0 | 21 |
| GCN20 | P43535 | Protein GCN20 | 752 | 1.33 | 1 | 0 | 1 |
| RPN11 | P43588 | Ubiquitin carboxyl-terminal hydrolase RPN11 | 306 | 4.25 | 1 | 0 | 2 |
| DUG1 | P43616 | Cys-Gly metallodipeptidase DUG1 | 481 | 19.33 | 7 | 0 | 29 |
| FIP1 | P45976 | Pre-mRNA polyadenylation factor FIP1 | 327 | 9.79 | 2 | 0 | 4 |

| Gene | Protein ID | Protein description | Length (aa) | Coverage (%) | Total peptides | Unique peptides | Total spectral count |
|---|---|---|---|---|---|---|---|
| ALD4 | P46367 | Potassium-activated aldehyde dehydrogenase, mitochondrial | 519 | 2.70 | 1 | 0 | 3 |
| GUS1 | P46655 | Glutamate--tRNA ligase, cytoplasmic | 708 | 32.91 | 23 | 0 | 61 |
| ARC1 | P46672 | tRNA-aminoacylation cofactor ARC1 | 376 | 26.06 | 7 | 0 | 14 |
| AIP1 | P46680 | Actin-interacting protein 1 | 615 | 4.39 | 2 | 0 | 2 |
| RPS10B | P46784 | Small ribosomal subunit protein eS10B | 105 | 30.48 | 2 | 0 | 13 |
| FYV8 | P46949 | Protein FYV8 | 817 | 1.84 | 1 | 0 | 4 |
| RPL17B | P46990 | Large ribosomal subunit protein uL22B | 184 | 47.83 | 8 | 0 | 40 |
| ALY2 | P47029 | Arrestin-related trafficking adapter 3 | 1046 | 1.15 | 1 | 0 | 1 |
| CCT8 | P47079 | T-complex protein 1 subunit theta | 568 | 4.40 | 3 | 0 | 4 |
| MPP10 | P47083 | U3 small nucleolar RNA-associated protein MPP10 | 593 | 6.91 | 3 | 0 | 4 |
| ARP3 | P47117 | Actin-related protein 3 | 449 | 4.90 | 2 | 0 | 2 |
| LIA1 | P47120 | Deoxyhypusine hydroxylase | 325 | 9.85 | 2 | 0 | 2 |
| MOG1 | P47123 | Nuclear import protein MOG1 | 218 | 64.68 | 9 | 0 | 130 |
| ACF4 | P47129 | Assembly-complementing factor 4 | 309 | 10.36 | 2 | 0 | 7 |
| JSN1 | P47135 | Protein JSN1 | 1091 | 9.44 | 7 | 0 | 16 |
| ADO1 | P47143 | Adenosine kinase | 340 | 25.29 | 6 | 0 | 13 |
| RSM7 | P47150 | Small ribosomal subunit protein uS7m | 247 | 10.93 | 2 | 0 | 7 |
| BAT2 | P47176 | Branched-chain-amino-acid aminotransferase, cytosolic | 376 | 18.62 | 5 | 0 | 20 |
| FAA4 | P47912 | Long-chain-fatty-acid--CoA ligase 4 | 694 | 4.76 | 3 | 0 | 5 |
| RPS7B | P48164 | Small ribosomal subunit protein eS7B | 190 | 54.21 | 10 | 0 | 39 |
| SEC16 | P48415 | COPII coat assembly protein SEC16 | 2195 | 14.81 | 24 | 0 | 54 |
| LYS20 | P48570 | Homocitrate synthase, cytosolic isozyme | 428 | 11.92 | 4 | 0 | 10 |
| RPS12 | P48589 | Small ribosomal subunit protein eS12 | 143 | 46.85 | 6 | 0 | 28 |
| RBL2 | P48606 | Tubulin-specific chaperone A | 106 | 7.55 | 1 | 0 | 2 |
| VMA10 | P48836 | V-type proton ATPase subunit G | 114 | 18.42 | 2 | 0 | 5 |
| NUP57 | P48837 | Nucleoporin NUP57 | 541 | 5.73 | 3 | 0 | 5 |
| ASN2 | P49090 | Asparagine synthetase [glutamine-hydrolyzing] 2 | 572 | 7.87 | 4 | 0 | 5 |
| RPL37A | P49166 | Large ribosomal subunit protein eL37A | 88 | 7.95 | 1 | 0 | 1 |
| RPL38 | P49167 | Large ribosomal subunit protein eL38 | 78 | 51.28 | 4 | 0 | 8 |
| APT1 | P49435 | Adenine phosphoribosyltransferase 1 | 187 | 12.83 | 2 | 0 | 5 |
| RPL4B | P49626 | Large ribosomal subunit protein uL4B | 362 | 52.76 | 15 | 0 | 73 |
| NUP145 | P49687 | Nucleoporin NUP145 | 1317 | 0.76 | 1 | 0 | 3 |
| RNR4 | P49723 | Ribonucleoside-diphosphate reductase small chain 2 | 345 | 19.71 | 5 | 0 | 21 |
| PHB2 | P50085 | Prohibitin-2 | 310 | 12.58 | 3 | 0 | 14 |
| IMD4 | P50094 | Inosine-5'-monophosphate dehydrogenase 4 | 524 | 22.33 | 8 | 0 | 26 |
| IMD3 | P50095 | Inosine-5'-monophosphate dehydrogenase 3 | 523 | 37.86 | 14 | 0 | 48 |
| PSP2 | P50109 | Protein PSP2 | 593 | 3.04 | 1 | 0 | 1 |
| PSP1 | P50896 | Protein PSP1 | 841 | 1.78 | 1 | 0 | 1 |

| Gene | Protein ID | Protein description | Length (aa) | Coverage (%) | Total peptides | Unique peptides | Total spectral count |
|---|---|---|---|---|---|---|---|
| YML6 | P51998 | Large ribosomal subunit protein uL4m | 286 | 4.55 | 1 | 0 | 2 |
| YKE2 | P52553 | Prefoldin subunit 6 | 114 | 28.07 | 2 | 0 | 2 |
| ACS2 | P52910 | Acetyl-coenzyme A synthetase 2 | 683 | 9.08 | 5 | 0 | 10 |
| TAF6 | P53040 | Transcription initiation factor TFIID subunit 6 | 516 | 5.43 | 2 | 0 | 3 |
| ARO8 | P53090 | Aromatic/aminoadipate aminotransferase 1 | 500 | 24.20 | 9 | 0 | 20 |
| NUT1 | P53114 | Mediator of RNA polymerase II transcription subunit 5 | 1132 | 1.06 | 1 | 0 | 1 |
| MLC1 | P53141 | Myosin light chain 1 | 149 | 30.20 | 4 | 0 | 8 |
| LSG1 | P53145 | Large subunit GTPase 1 | 640 | 9.53 | 5 | 0 | 10 |
| MNP1 | P53163 | Large ribosomal subunit protein bL12m | 194 | 18.56 | 4 | 0 | 8 |
| DUO1 | P53168 | DASH complex subunit DUO1 | 247 | 10.93 | 2 | 0 | 2 |
| NQM1 | P53228 | Transaldolase NQM1 | 333 | 2.40 | 1 | 0 | 3 |
| SLX9 | P53251 | Ribosome biogenesis protein SLX9 | 210 | 13.33 | 2 | 0 | 3 |
| PIL1 | P53252 | Sphingolipid long chain base-responsive protein PIL1 | 339 | 26.25 | 6 | 0 | 23 |
| DAM1 | P53267 | DASH complex subunit DAM1 | 343 | 10.79 | 3 | 0 | 5 |
| YGR130C | P53278 | Uncharacterized protein YGR130C | 816 | 14.83 | 10 | 0 | 21 |
| RBG2 | P53295 | Ribosome-interacting GTPase 2 | 368 | 3.53 | 1 | 0 | 1 |
| TIM13 | P53299 | Mitochondrial import inner membrane translocase subunit TIM13 | 105 | 31.43 | 2 | 0 | 6 |
| RPT4 | P53549 | 26S proteasome subunit RPT4 | 437 | 2.75 | 1 | 0 | 3 |
| COP1 | P53622 | Coatomer subunit alpha | 1201 | 4.25 | 3 | 0 | 7 |
| SNF12 | P53628 | Transcription regulatory protein SNF12 | 566 | 4.95 | 2 | 0 | 2 |
| CPR6 | P53691 | Peptidyl-prolyl cis-trans isomerase CPR6 | 371 | 9.16 | 3 | 0 | 9 |
| ESF2 | P53743 | Pre-rRNA-processing protein ESF2 | 316 | 3.16 | 1 | 0 | 1 |
| YNL247W | P53852 | Cysteine--tRNA ligase | 767 | 3.39 | 2 | 0 | 2 |
| BNI4 | P53858 | Protein BNI4 | 892 | 3.14 | 2 | 0 | 3 |
| MRPL22 | P53881 | Large ribosomal subunit protein uL22m | 309 | 5.18 | 1 | 0 | 1 |
| APC1 | P53886 | Anaphase-promoting complex subunit 1 | 1748 | 2.40 | 3 | 0 | 4 |
| YNL134C | P53912 | Uncharacterized protein YNL134C | 376 | 12.23 | 3 | 0 | 10 |
| IMP4 | P53941 | U3 small nucleolar ribonucleoprotein protein IMP4 | 290 | 3.45 | 1 | 0 | 2 |
| COG6 | P53959 | Conserved oligomeric Golgi complex subunit 6 | 839 | 1.07 | 1 | 0 | 1 |
| SUB1 | P54000 | RNA polymerase II transcriptional coactivator SUB1 | 292 | 5.14 | 1 | 0 | 1 |
| ALD6 | P54115 | Magnesium-activated aldehyde dehydrogenase, cytosolic | 500 | 35.60 | 15 | 0 | 33 |
| DAK1 | P54838 | Dihydroxyacetone kinase 1 | 584 | 4.62 | 2 | 0 | 5 |
| ERG13 | P54839 | Hydroxymethylglutaryl-CoA synthase | 491 | 8.76 | 3 | 0 | 5 |
| DOS2 | P54858 | Protein DOS2 | 310 | 5.16 | 1 | 0 | 3 |
| PRO2 | P54885 | Gamma-glutamyl phosphate reductase | 456 | 8.11 | 3 | 0 | 5 |
| RPL22B | P56628 | Large ribosomal subunit protein eL22B | 122 | 38.52 | 3 | 0 | 24 |
| ACT1 | P60010 | Actin | 375 | 58.67 | 17 | 0 | 76 |
| LYZ | P61626 | Lysozyme C | 148 | 61.49 | 8 | 8 | 40 |

| Gene | Protein ID | Protein description | Length (aa) | Coverage (%) | Total peptides | Unique peptides | Total spectral count |
|---|---|---|---|---|---|---|---|
| B2M | P61769 | Beta-2-microglobulin | 119 | 8.40 | 1 | 1 | 1 |
| HBB | P68871 | Hemoglobin subunit beta | 147 | 21.77 | 2 | 2 | 5 |
| DAD3 | P69850 | DASH complex subunit DAD3 | 94 | 20.21 | 2 | 0 | 5 |
| KRT85 | P78386 | Keratin, type II cuticular Hb5 | 507 | 2.96 | 2 | 0 | 3 |
| TIM11 | P81449 | ATP synthase subunit e, mitochondrial | 96 | 10.42 | 1 | 0 | 1 |
| GPD1 | Q00055 | Glycerol-3-phosphate dehydrogenase [NAD(+)] 1 | 391 | 2.05 | 1 | 0 | 1 |
| TPS1 | Q00764 | Alpha,alpha-trehalose-phosphate synthase [UDP-forming] 56 kDa subunit | 495 | 3.23 | 1 | 0 | 1 |
| REG1 | Q00816 | Resistance to glucose repression protein 1 | 1014 | 5.82 | 5 | 0 | 5 |
| ACC1 | Q00955 | Acetyl-CoA carboxylase | 2233 | 4.93 | 8 | 0 | 22 |
| UBP3 | Q01477 | Ubiquitin carboxyl-terminal hydrolase 3 | 912 | 25.77 | 21 | 0 | 64 |
| MAP1 | Q01662 | Methionine aminopeptidase 1 | 387 | 2.58 | 1 | 0 | 2 |
| GCR2 | Q01722 | Glycolytic genes transcriptional activator GCR2 | 534 | 1.50 | 1 | 0 | 1 |
| TIM44 | Q01852 | Mitochondrial import inner membrane translocase subunit TIM44 | 431 | 14.85 | 5 | 0 | 6 |
| RPS15 | Q01855 | Small ribosomal subunit protein uS19 | 142 | 30.28 | 3 | 0 | 25 |
| RPT6 | Q01939 | 26S proteasome regulatory subunit 8 homolog | 405 | 11.11 | 3 | 0 | 6 |
| OSH6 | Q02201 | Oxysterol-binding protein homolog 6 | 448 | 6.03 | 2 | 0 | 4 |
| MRPL13 | Q02204 | Large ribosomal subunit protein mL50 | 264 | 47.73 | 7 | 0 | 19 |
| RPL6A | Q02326 | Large ribosomal subunit protein eL6A | 176 | 53.98 | 10 | 0 | 28 |
| TBF1 | Q02457 | Protein TBF1 | 562 | 2.67 | 1 | 0 | 1 |
| ABF2 | Q02486 | ARS-binding factor 2, mitochondrial | 183 | 9.84 | 2 | 0 | 2 |
| YPL014W | Q02606 | Uncharacterized protein YPL014W | 381 | 4.99 | 1 | 0 | 3 |
| RPL21A | Q02753 | Large ribosomal subunit protein eL21A | 160 | 35.00 | 5 | 0 | 23 |
| SRP1 | Q02821 | Importin subunit alpha | 542 | 7.38 | 4 | 0 | 7 |
| MUK1 | Q02866 | Protein MUK1 | 612 | 1.63 | 1 | 0 | 1 |
| VPS30 | Q02948 | Vacuolar protein sorting-associated protein 30 | 557 | 1.26 | 1 | 0 | 3 |
| GIP3 | Q03016 | GLC7-interacting protein 3 | 1276 | 2.66 | 3 | 0 | 6 |
| COF1 | Q03048 | Cofilin | 143 | 38.46 | 4 | 0 | 13 |
| DIG1 | Q03063 | Down-regulator of invasive growth 1 | 452 | 9.51 | 3 | 0 | 8 |
| SVL3 | Q03088 | Styryl dye vacuolar localization protein 3 | 825 | 3.15 | 2 | 0 | 2 |
| YMR099C | Q03161 | Glucose-6-phosphate 1-epimerase | 297 | 7.74 | 2 | 0 | 4 |
| MRPS17 | Q03246 | Small ribosomal subunit protein uS17m | 237 | 10.55 | 2 | 0 | 3 |
| TFB3 | Q03290 | RNA polymerase II transcription factor B subunit 3 | 321 | 24.61 | 6 | 0 | 8 |
| OYE2 | Q03558 | NADPH dehydrogenase 2 | 400 | 29.50 | 9 | 0 | 45 |
| CLU1 | Q03690 | Clustered mitochondria protein 1 | 1277 | 1.17 | 1 | 0 | 2 |
| NAB6 | Q03735 | RNA-binding protein NAB6 | 1134 | 11.73 | 11 | 0 | 27 |
| TAF12 | Q03761 | Transcription initiation factor TFIID subunit 12 | 539 | 6.68 | 3 | 0 | 7 |
| ENT5 | Q03769 | Epsin-5 | 411 | 2.68 | 1 | 0 | 1 |
| ARX1 | Q03862 | Probable metalloprotease ARX1 | 593 | 2.36 | 1 | 0 | 6 |

| Gene | Protein ID | Protein description | Length (aa) | Coverage (%) | Total peptides | Unique peptides | Total spectral count |
|---|---|---|---|---|---|---|---|
| RVB1 | Q03940 | RuvB-like protein 1 | 463 | 19.01 | 6 | 0 | 8 |
| SPC19 | Q03954 | DASH complex subunit SPC19 | 165 | 14.55 | 2 | 0 | 6 |
| SND1 | Q04007 | SRP-independent targeting protein 1 | 877 | 2.96 | 2 | 0 | 4 |
| YHM2 | Q04013 | Citrate/oxoglutarate carrier protein | 314 | 6.69 | 1 | 0 | 1 |
| RPN9 | Q04062 | 26S proteasome regulatory subunit RPN9 | 393 | 2.29 | 1 | 0 | 1 |
| TIF35 | Q04067 | Eukaryotic translation initiation factor 3 subunit G | 274 | 27.74 | 5 | 0 | 20 |
| HNT1 | Q04344 | Adenosine 5'-monophosphoramidase HNT1 | 158 | 20.89 | 2 | 0 | 6 |
| RBA50 | Q04418 | RNA polymerase II-associated protein RBA50 | 439 | 10.25 | 4 | 0 | 7 |
| SEC13 | Q04491 | Protein transport protein SEC13 | 297 | 24.24 | 6 | 0 | 7 |
| MRPL1 | Q04599 | Large ribosomal subunit protein uL1m | 285 | 9.12 | 2 | 0 | 3 |
| YMR315W | Q04869 | Uncharacterized protein YMR315W | 349 | 16.62 | 4 | 0 | 5 |
| ADH6 | Q04894 | NADP-dependent alcohol dehydrogenase 6 | 360 | 5.00 | 1 | 0 | 3 |
| IVY1 | Q04934 | Protein IVY1 | 453 | 6.40 | 2 | 0 | 3 |
| RTN1 | Q04947 | Reticulon-like protein 1 | 295 | 23.05 | 4 | 0 | 9 |
| YMR226C | Q05016 | NADP-dependent 3-hydroxy acid dehydrogenase | 267 | 3.75 | 1 | 0 | 2 |
| RRP5 | Q05022 | rRNA biogenesis protein RRP5 | 1729 | 1.62 | 3 | 0 | 4 |
| TAF9 | Q05027 | Transcription initiation factor TFIID subunit 9 | 157 | 12.74 | 1 | 0 | 3 |
| ARP9 | Q05123 | Actin-like protein ARP9 | 467 | 4.71 | 2 | 0 | 3 |
| ASM4 | Q05166 | Nucleoporin ASM4 | 528 | 3.41 | 1 | 0 | 1 |
| RTT103 | Q05543 | Regulator of Ty1 transposition protein 103 | 409 | 22.74 | 8 | 0 | 23 |
| GLO2 | Q05584 | Hydroxyacylglutathione hydrolase, cytoplasmic isozyme | 274 | 13.14 | 3 | 0 | 4 |
| HCR1 | Q05775 | Eukaryotic translation initiation factor 3 subunit J | 265 | 13.96 | 2 | 0 | 4 |
| ADE13 | Q05911 | Adenylosuccinate lyase | 482 | 25.93 | 10 | 0 | 18 |
| COG4 | Q06096 | Conserved oligomeric Golgi complex subunit 4 | 861 | 1.28 | 1 | 0 | 1 |
| KAP95 | Q06142 | Importin subunit beta-1 | 861 | 10.57 | 8 | 0 | 29 |
| YLR177W | Q06251 | Uncharacterized protein YLR177W | 628 | 2.23 | 1 | 0 | 1 |
| YLR179C | Q06252 | Uncharacterized protein YLR179C | 201 | 38.31 | 6 | 0 | 21 |
| MMR1 | Q06324 | Mitochondrial MYO2 receptor-related protein 1 | 491 | 5.30 | 2 | 0 | 2 |
| RGA2 | Q06407 | Rho-type GTPase-activating protein 2 | 1009 | 0.89 | 1 | 0 | 1 |
| CRN1 | Q06440 | Coronin-like protein | 651 | 3.23 | 2 | 0 | 4 |
| SPN1 | Q06505 | Transcription factor SPN1 | 410 | 6.59 | 2 | 0 | 6 |
| RRP15 | Q06511 | Ribosomal RNA-processing protein 15 | 250 | 4.40 | 1 | 0 | 1 |
| BFR2 | Q06631 | Protein BFR2 | 534 | 25.28 | 11 | 0 | 24 |
| CSR1 | Q06705 | Phosphatidylinositol transfer protein CSR1 | 408 | 21.08 | 7 | 0 | 15 |
| PRDX1 | Q06830 | Peroxiredoxin-1 | 199 | 29.15 | 6 | 6 | 10 |
| YPR097W | Q06839 | PX domain-containing protein YPR097W | 1073 | 0.84 | 1 | 0 | 1 |
| PBP4 | Q07362 | Protein PBP4 | 185 | 7.57 | 1 | 0 | 6 |
| BRE1 | Q07457 | E3 ubiquitin-protein ligase BRE1 | 700 | 1.43 | 1 | 0 | 3 |

| Gene | Protein ID | Protein description | Length (aa) | Coverage (%) | Total peptides | Unique peptides | Total spectral count |
|---|---|---|---|---|---|---|---|
| SHS1 | Q07657 | Seventh homolog of septin 1 | 551 | 24.68 | 11 | 0 | 24 |
| PUF3 | Q07807 | mRNA-binding protein PUF3 | 879 | 13.20 | 9 | 0 | 18 |
| VPS13 | Q07878 | Intermembrane lipid transfer protein VPS13 | 3144 | 0.38 | 1 | 0 | 1 |
| RLP24 | Q07915 | Ribosome biogenesis protein RLP24 | 199 | 5.03 | 1 | 0 | 1 |
| BUD20 | Q08004 | Bud site selection protein 20 | 166 | 14.46 | 2 | 0 | 5 |
| DIS3 | Q08162 | Exosome complex exonuclease DIS3 | 1001 | 1.50 | 1 | 0 | 1 |
| ZEO1 | Q08245 | Protein ZEO1 | 113 | 46.90 | 8 | 0 | 26 |
| NOB1 | Q08444 | 20S-pre-rRNA D-site endonuclease NOB1 | 459 | 2.61 | 1 | 0 | 3 |
| RPN8 | Q08723 | 26S proteasome regulatory subunit RPN8 | 338 | 3.55 | 1 | 0 | 2 |
| RSA1 | Q08932 | Ribosome assembly 1 protein | 381 | 3.67 | 1 | 0 | 1 |
| YPL225W | Q08971 | Protein PBDC1 homolog | 146 | 45.21 | 6 | 0 | 13 |
| NEW1 | Q08972 | [NU+] prion formation protein 1 | 1196 | 2.84 | 2 | 0 | 9 |
| TMA46 | Q12000 | Translation machinery-associated protein 46 | 345 | 6.96 | 2 | 0 | 5 |
| MDN1 | Q12019 | Midasin | 4910 | 0.29 | 1 | 0 | 1 |
| FSF1 | Q12029 | Sideroflexin FSF1 | 327 | 4.28 | 1 | 0 | 1 |
| SLF1 | Q12034 | Protein SLF1 | 447 | 6.04 | 2 | 0 | 4 |
| FCF2 | Q12035 | rRNA-processing protein FCF2 | 217 | 6.91 | 1 | 0 | 1 |
| SPE3 | Q12074 | Spermidine synthase SPE3 | 293 | 13.99 | 3 | 0 | 6 |
| SGT2 | Q12118 | Small glutamine-rich tetratricopeptide repeat-containing protein 2 | 346 | 17.05 | 4 | 0 | 9 |
| SAS10 | Q12136 | Something about silencing protein 10 | 610 | 9.02 | 3 | 0 | 11 |
| YRA1 | Q12159 | RNA annealing protein YRA1 | 226 | 38.94 | 8 | 0 | 24 |
| FCY1 | Q12178 | Cytosine deaminase | 158 | 17.72 | 2 | 0 | 2 |
| RPL7B | Q12213 | Large ribosomal subunit protein uL30B | 244 | 46.31 | 12 | 0 | 28 |
| LSP1 | Q12230 | Sphingolipid long chain base-responsive protein LSP1 | 341 | 24.63 | 6 | 0 | 29 |
| HER1 | Q12276 | HMG2-induced ER-remodeling protein 1 | 1246 | 0.72 | 1 | 0 | 1 |
| RDL1 | Q12305 | Thiosulfate:glutathione sulfurtransferase | 139 | 60.43 | 6 | 0 | 12 |
| HSP42 | Q12329 | Heat shock protein 42 | 375 | 7.47 | 2 | 0 | 3 |
| WTM1 | Q12363 | Transcriptional modulator WTM1 | 437 | 10.53 | 4 | 0 | 5 |
| AHA1 | Q12449 | Hsp90 co-chaperone AHA1 | 350 | 12.00 | 4 | 0 | 9 |
| OSH2 | Q12451 | Oxysterol-binding protein homolog 2 | 1283 | 3.20 | 3 | 0 | 12 |
| CEX1 | Q12453 | Cytoplasmic export protein 1 | 761 | 5.91 | 4 | 0 | 5 |
| YPR1 | Q12458 | Putative reductase 1 | 312 | 12.82 | 3 | 0 | 4 |
| NOP56 | Q12460 | Nucleolar protein 56 | 504 | 6.55 | 2 | 0 | 2 |
| RVB2 | Q12464 | RuvB-like protein 2 | 471 | 19.32 | 7 | 0 | 10 |
| TIF6 | Q12522 | Eukaryotic translation initiation factor 6 | 245 | 11.43 | 2 | 0 | 2 |
| GLT1 | Q12680 | Glutamate synthase [NADH] | 2145 | 3.59 | 6 | 0 | 9 |
| KRT33B | Q14525 | Keratin, type I cuticular Ha3-II | 404 | 11.63 | 5 | 0 | 19 |
| KRT31 | Q15323 | Keratin, type I cuticular Ha1 | 416 | 13.46 | 6 | 0 | 20 |

| Gene | Protein ID | Protein description | Length (aa) | Coverage (%) | Total peptides | Unique peptides | Total spectral count |
|---|---|---|---|---|---|---|---|
| SLO1 | Q3E784 | SCOCO-like protein 1 | 85 | 14.12 | 1 | 0 | 1 |
| YNR034W-A | Q3E841 | Uncharacterized protein YNR034W-A | 98 | 32.65 | 2 | 0 | 3 |
| KRT35 | Q92764 | Keratin, type I cuticular Ha5 | 455 | 6.15 | 3 | 0 | 16 |
| PNO1 | Q99216 | Pre-rRNA-processing protein PNO1 | 274 | 14.23 | 3 | 0 | 4 |
| DUF1 | Q99247 | DUB-associated factor 1 | 1116 | 1.16 | 1 | 0 | 1 |
| MEX67 | Q99257 | mRNA export factor MEX67 | 599 | 4.84 | 2 | 0 | 5 |
| MPD2 | Q99316 | Protein disulfide isomerase MPD2 | 277 | 12.64 | 3 | 0 | 6 |
| KRT84 | Q9NSB2 | Keratin, type II cuticular Hb4 | 600 | 5.83 | 6 | 3 | 18 |
| PFK2 | A0A8H4BY05 | ATP-dependent 6-phosphofructokinase | 959 | 17.10 | 12 | 0 | 22 |
| NOG1 | A0A8H4F6Y2 | Nucleolar GTP-binding protein 1 | 647 | 2.01 | 1 | 0 | 1 |
| RRP6 | A0A8H4F761 | RRP6 isoform 1 | 733 | 2.05 | 1 | 0 | 2 |
| SCY_5346 | A6ZPD1 | Conserved protein | 1289 | 0.70 | 1 | 0 | 4 |
| GRX3 | A6ZY62 | Glutaredoxin | 285 | 5.61 | 1 | 0 | 1 |
| AWRI1631_91530 | B5VKU0 | YIR004Wp-like protein (Fragment) | 451 | 3.77 | 1 | 0 | 6 |
| BRE5 | C7GLK6 | Bre5p | 515 | 29.90 | 11 | 0 | 24 |
| EC1118_1G1_3609g | C8Z8U8 | Eukaryotic translation initiation factor 2A | 642 | 5.30 | 2 | 0 | 4 |
| K7_SRB4 | G2WCQ5 | Mediator of RNA polymerase II transcription subunit 17 | 687 | 1.16 | 1 | 0 | 1 |
|  | P02534 | Keratin, type I microfibrillar 48 kDa, component 8C-1 | 412 | 11.17 | 5 | 0 | 18 |
| HTA1 | P04911 | Histone H2A.1 | 132 | 12.12 | 2 | 0 | 3 |
| RPL31B | P0C2H9 | Large ribosomal subunit protein eL31B | 113 | 61.95 | 8 | 0 | 26 |
| SAM2 | P19358 | S-adenosylmethionine synthase 2 | 384 | 18.49 | 6 | 0 | 12 |
| TIF3 | P34167 | Eukaryotic translation initiation factor 4B | 436 | 1.83 | 1 | 0 | 1 |
| PHO90 | P39535 | Low-affinity phosphate transporter PHO90 | 881 | 1.36 | 1 | 0 | 1 |
| RPS27A | P62979 | Ubiquitin-40S ribosomal protein S27a | 156 | 27.56 | 4 | 1 | 21 |
| HOT1 | Q03213 | High-osmolarity-induced transcription protein 1 | 719 | 2.36 | 1 | 0 | 1 |
| DIG2 | Q03373 | Down-regulator of invasive growth 2 | 323 | 3.41 | 1 | 0 | 1 |
| MRPL35 | Q06678 | Large ribosomal subunit protein mL38 | 367 | 3.00 | 1 | 0 | 1 |
| DNF2 | Q12675 | Phospholipid-transporting ATPase DNF2 | 1612 | 0.56 | 1 | 0 | 1 |
| ADE16 | A7A0T4 | 5-aminoimidazole-4-carboxamide ribonucleotide (AICAR) transformylase/IMP cyclohydrolase | 591 | 9.31 | 4 | 0 | 7 |
| SCRG_00677 | B3LH69 | H(+)-transporting two-sector ATPase | 617 | 27.71 | 15 | 0 | 35 |
| SCRG_00930 | B3LID9 | branched-chain-2-oxoacid decarboxylase | 563 | 11.01 | 4 | 0 | 37 |
| AWRI1631_73850 | B5VJA5 | YGR162Wp-like protein | 842 | 22.68 | 16 | 0 | 38 |
| AWRI1631_111660 | B5VM96 | Fructose-bisphosphate aldolase | 217 | 27.65 | 5 | 1 | 24 |
| EC1118_1J19_0188g | C8ZBM1 | Mog1p | 188 | 65.96 | 7 | 0 | 115 |

| Gene | Protein ID | Protein description | Length (aa) | Coverage (%) | Total peptides | Unique peptides | Total spectral count |
|---|---|---|---|---|---|---|---|
| K7_RPL13A | G2WC81 | 60S ribosomal protein L13 | 199 | 37.19 | 6 | 1 | 15 |
| K7_LAT1 | G2WM33 | Acetyltransferase component of pyruvate dehydrogenase complex | 482 | 34.85 | 13 | 1 | 47 |
| HSP150 | A6ZQH3 | Cell wall mannoprotein HSP150 | 389 | 4.63 | 2 | 0 | 3 |
| TSA2 | Q04120 | Peroxiredoxin TSA2 | 196 | 4.08 | 1 | 0 | 4 |
| RPL13A | Q12690 | Large ribosomal subunit protein eL13A | 199 | 44.22 | 8 | 0 | 26 |
| HTZ1 | Q12692 | Histone H2A.Z | 134 | 11.94 | 2 | 0 | 6 |
| RPS21B | Q3E754 | Small ribosomal subunit protein eS21B | 87 | 68.97 | 6 | 0 | 23 |
| CIS3 | A6ZQH4 | Cell wall mannoprotein CIS3 | 225 | 3.56 | 1 | 0 | 2 |
| PYK2 | P52489 | Pyruvate kinase 2 | 506 | 3.95 | 3 | 0 | 6 |
| RPL26B | P53221 | Large ribosomal subunit protein uL24B | 127 | 48.82 | 10 | 0 | 28 |
| FAA1 | P30624 | Long-chain-fatty-acid--CoA ligase 1 | 700 | 1.71 | 1 | 0 | 1 |
| TY1B-JR2 | P47100 | Transposon Ty1-JR2 Gag-Pol polyprotein | 1755 | 11.45 | 14 | 0 | 35 |
| IMD2 | P38697 | Inosine-5'-monophosphate dehydrogenase 2 | 523 | 23.71 | 8 | 0 | 31 |
| TY1B-ML2 | Q03434 | Transposon Ty1-ML2 Gag-Pol polyprotein | 1755 | 12.36 | 15 | 0 | 37 |
| SHE2 | A6ZZJ0 | SWI5-dependent HO expression protein 2 | 246 | 5.69 | 1 | 0 | 1 |
| LSB3 | A7A261 | LAS seventeen-binding protein 3 | 459 | 3.92 | 1 | 0 | 1 |
| SEM1 | O94742 | 26S proteasome complex subunit SEM1 | 89 | 13.48 | 1 | 0 | 1 |
| SOD1 | P00441 | Superoxide dismutase [Cu-Zn] | 154 | 9.09 | 1 | 1 | 2 |
| HIS1 | P00498 | ATP phosphoribosyltransferase | 297 | 4.04 | 1 | 0 | 1 |
| MSW1 | P04803 | Tryptophan--tRNA ligase, mitochondrial | 379 | 2.90 | 1 | 0 | 5 |
| FUM1 | P08417 | Fumarate hydratase, mitochondrial | 488 | 2.66 | 1 | 0 | 1 |
| GSTP1 | P09211 | Glutathione S-transferase P | 210 | 7.62 | 1 | 1 | 2 |
| RPP1B | P10622 | Large ribosomal subunit protein P1B | 106 | 15.09 | 1 | 0 | 6 |
| HEM15 | P16622 | Ferrochelatase, mitochondrial | 393 | 5.34 | 1 | 0 | 1 |
| KGD2 | P19262 | Dihydrolipoyllysine-residue succinyltransferase component of 2-oxoglutarate dehydrogenase complex, mitochondrial | 463 | 3.46 | 1 | 0 | 2 |
| SCL1 | P21243 | Proteasome subunit alpha type-1 | 252 | 5.16 | 1 | 0 | 1 |
| RRP43 | P25359 | Exosome complex component RRP43 | 394 | 2.79 | 1 | 0 | 2 |
| SBA1 | P28707 | Co-chaperone protein SBA1 | 216 | 5.09 | 1 | 0 | 6 |
| DPH5 | P32469 | Diphthine methyl ester synthase | 300 | 4.33 | 1 | 0 | 1 |
| SUI1 | P32911 | Eukaryotic translation initiation factor eIF-1 | 108 | 20.37 | 1 | 0 | 3 |
| YPT7 | P32939 | Ypt/Rab-type GTPase YPT7 | 208 | 6.25 | 1 | 0 | 1 |
| SSS1 | P35179 | Protein transport protein SSS1 | 80 | 12.50 | 1 | 0 | 2 |
| MUD2 | P36084 | Splicing factor MUD2 | 527 | 3.23 | 1 | 0 | 1 |
| MRPL17 | P36528 | Large ribosomal subunit protein mL46 | 281 | 4.98 | 1 | 0 | 1 |
| YGP1 | P38616 | Protein YGP1 | 354 | 3.39 | 1 | 0 | 2 |
| VMA7 | P39111 | V-type proton ATPase subunit F | 118 | 14.41 | 1 | 0 | 5 |
| PRE5 | P40302 | Proteasome subunit alpha type-6 | 234 | 5.98 | 1 | 0 | 6 |

| Gene | Protein ID | Protein description | Length (aa) | Coverage (%) | Total peptides | Unique peptides | Total spectral count |
|---|---|---|---|---|---|---|---|
| RLP7 | P40693 | Ribosome biogenesis protein RLP7 | 322 | 4.04 | 1 | 0 | 3 |
| GLR1 | P41921 | Glutathione reductase | 483 | 2.48 | 1 | 0 | 2 |
| NNF1 | P47149 | Kinetochore-associated protein NNF1 | 201 | 5.97 | 1 | 0 | 1 |
| FOL2 | P51601 | GTP cyclohydrolase 1 | 243 | 4.94 | 1 | 0 | 2 |
| HSK3 | P69852 | DASH complex subunit HSK3 | 69 | 15.94 | 1 | 0 | 2 |
| NAM8 | Q00539 | Protein NAM8 | 523 | 2.49 | 1 | 0 | 3 |
| YSA1 | Q01976 | ADP-ribose pyrophosphatase | 231 | 6.49 | 1 | 0 | 1 |
| HPT1 | Q04178 | Hypoxanthine-guanine phosphoribosyltransferase | 221 | 5.43 | 1 | 0 | 2 |
| ARC18 | Q05933 | Actin-related protein 2/3 complex subunit 3 | 178 | 7.30 | 1 | 0 | 1 |
| MEU1 | Q07938 | S-methyl-5'-thioadenosine phosphorylase | 337 | 3.26 | 1 | 0 | 3 |
| TAF10 | Q12030 | Transcription initiation factor TFIID subunit 10 | 206 | 9.22 | 1 | 0 | 1 |
| APC9 | Q12107 | Anaphase-promoting complex subunit 9 | 265 | 6.42 | 1 | 0 | 2 |
| TAF3 | Q12297 | Transcription initiation factor TFIID subunit 3 | 353 | 4.82 | 1 | 0 | 4 |
| DAD2 | P36162 | DASH complex subunit DAD2 | 133 | 8.27 | 1 | 0 | 1 |
| NPT1 | P39683 | Nicotinate phosphoribosyltransferase | 429 | 3.96 | 1 | 0 | 1 |
| RSM23 | Q01163 | Small ribosomal subunit protein mS29 | 450 | 2.22 | 1 | 0 | 1 |
| GPD2 | P41911 | Glycerol-3-phosphate dehydrogenase [NAD(+)] 2, mitochondrial | 440 | 2.95 | 1 | 0 | 2 |
| PNC1 | P53184 | Nicotinamidase | 216 | 4.17 | 1 | 0 | 1 |
| SRPP | O82803 | Small rubber particle protein | 204 | 4.90 | 1 | 1 | 1 |
| GDA1 | P32621 | Guanosine-diphosphatase | 518 | 4.05 | 1 | 0 | 3 |
| YBR226C | P38322 | Putative uncharacterized protein YBR226C | 136 | 13.24 | 1 | 1 | 37 |
| PPT1 | P53043 | Serine/threonine-protein phosphatase T | 513 | 1.75 | 1 | 0 | 1 |
| WRS1 | Q12109 | Tryptophan--tRNA ligase, cytoplasmic | 432 | 4.17 | 1 | 0 | 1 |
| GAS1 | P22146 | 1,3-beta-glucanosyltransferase GAS1 | 559 | 2.33 | 1 | 0 | 1 |
| TMA17 | Q12513 | Translation machinery-associated protein 17 | 150 | 7.33 | 1 | 0 | 1 |
| RPC10 | P40422 | DNA-directed RNA polymerases I, II, and III subunit RPABC4 | 70 | 28.57 | 1 | 0 | 2 |
| SBP1 | P10080 | Single-stranded nucleic acid-binding protein | 294 | 8.16 | 1 | 0 | 1 |
| SLC1 | P33333 | 1-acyl-sn-glycerol-3-phosphate acyltransferase | 303 | 3.30 | 1 | 0 | 2 |
| YDL124W | Q07551 | NADPH-dependent alpha-keto amide reductase | 312 | 3.85 | 1 | 0 | 1 |
| TF | P02787 | Serotransferrin | 698 | 1.72 | 1 | 1 | 1 |
| REE1 | P40893 | Regulation of enolase protein 1 | 198 | 7.07 | 1 | 0 | 1 |
| FPR2 | P32472 | Peptidyl-prolyl cis-trans isomerase FPR2 | 135 | 10.37 | 1 | 0 | 1 |
| RPP1A | P05318 | Large ribosomal subunit protein P1A | 106 | 7.55 | 1 | 0 | 2 |
| WHI2 | P12611 | Growth regulation protein | 486 | 2.47 | 1 | 0 | 1 |
| RTT102 | P53330 | Regulator of Ty1 transposition protein 102 | 157 | 12.74 | 1 | 0 | 1 |
| SCW4 | P53334 | Probable family 17 glucosidase SCW4 | 386 | 3.63 | 1 | 0 | 1 |
| PEX19 | A6ZXR1 | Peroxisomal membrane protein import receptor PEX19 | 341 | 2.93 | 1 | 0 | 1 |
| HHT1 | P61830 | Histone H3 | 136 | 5.15 | 1 | 0 | 6 |

| Gene | Protein ID | Protein description | Length (aa) | Coverage (%) | Total peptides | Unique peptides | Total spectral count |
|---|---|---|---|---|---|---|---|
| ADK2 | E9P974 | GTP:AMP phosphotransferase, mitochondrial | 234 | 4.70 | 1 | 0 | 1 |
| EXG1 | P23776 | Glucan 1,3-beta-glucosidase I/II | 448 | 2.46 | 1 | 0 | 1 |
| GLC3 | P32775 | 1,4-alpha-glucan-branching enzyme | 704 | 1.28 | 1 | 0 | 1 |
| CKA2 | P19454 | Casein kinase II subunit alpha' | 339 | 2.65 | 1 | 0 | 1 |
| HMO1 | Q03973 | High mobility group protein 1 | 246 | 4.88 | 1 | 0 | 1 |
| SSP120 | P39931 | Protein SSP120 | 234 | 3.85 | 1 | 0 | 1 |
| VBA4 | Q04602 | Vacuolar basic amino acid transporter 4 | 768 | 4.82 | 1 | 0 | 1 |
| PHO88 | P38264 | SRP-independent targeting protein 3 | 188 | 7.45 | 1 | 0 | 1 |
| RSM25 | A6ZVI4 | Small ribosomal subunit protein mS23 | 264 | 2.65 | 1 | 0 | 1 |
| MCP2 | Q06567 | ABC1 family protein MCP2 | 569 | 1.76 | 1 | 0 | 1 |
| EMC1 | P25574 | ER membrane protein complex subunit 1 | 760 | 1.32 | 1 | 0 | 1 |
| SKS1 | Q12505 | Serine/threonine-protein kinase SKS1 | 502 | 2.79 | 1 | 0 | 2 |
| TOM22 | P49334 | Mitochondrial import receptor subunit TOM22 | 152 | 11.18 | 1 | 0 | 1 |
| PRP24 | P49960 | U4/U6 snRNA-associated-splicing factor PRP24 | 444 | 3.38 | 1 | 0 | 1 |
| GDI1 | P39958 | Rab GDP-dissociation inhibitor | 451 | 1.77 | 1 | 0 | 1 |
| YMR31 | P19955 | 37S ribosomal protein YMR-31, mitochondrial | 123 | 9.76 | 1 | 0 | 1 |
| RAD26 | P40352 | DNA repair and recombination protein RAD26 | 1085 | 0.92 | 1 | 0 | 1 |
| SWC3 | P31376 | SWR1-complex protein 3 | 625 | 1.76 | 1 | 0 | 2 |
| BYE1 | P36106 | Transcription factor BYE1 | 594 | 2.02 | 1 | 0 | 3 |
| LYS9 | P38999 | Saccharopine dehydrogenase [NADP(+), L-glutamate-forming] | 446 | 4.93 | 1 | 0 | 1 |
| PRP5 | A6ZLH6 | Pre-mRNA-processing ATP-dependent RNA helicase PRP5 | 849 | 0.82 | 1 | 0 | 4 |

#### Sheet 2. Mog1 band (88 proteins)

| Gene | Protein ID | Protein description | Length (aa) | Coverage (%) | Total peptides | Unique peptides | Total spectral count |
|---|---|---|---|---|---|---|---|
| RPS1B | A6ZM02 | Small ribosomal subunit protein eS1B | 255 | 15.69 | 4 | 0 | 19 |
| RKI1 | A6ZNU5 | Ribose-5-phosphate isomerase | 258 | 9.30 | 2 | 0 | 6 |
| RPS0A | A6ZUM5 | Small ribosomal subunit protein uS2A | 252 | 30.16 | 5 | 0 | 77 |
| RPS0B | A7A0V3 | Small ribosomal subunit protein uS2B | 252 | 30.16 | 5 | 0 | 77 |
| RPS1A | A7A1W1 | Small ribosomal subunit protein eS1A | 255 | 10.59 | 3 | 0 | 21 |
| KRT86 | O43790 | Keratin, type II cuticular Hb6 | 486 | 4.94 | 5 | 0 | 4 |
| KRT33A | O76009 | Keratin, type I cuticular Ha3-I | 404 | 2.23 | 2 | 0 | 3 |
| KRT34 | O76011 | Keratin, type I cuticular Ha4 | 436 | 2.06 | 2 | 1 | 4 |
| KRT15 | O77727 | Keratin, type I cytoskeletal 15 | 453 | 6.18 | 5 | 1 | 9 |
| ADH1 | P00330 | Alcohol dehydrogenase 1 | 348 | 6.61 | 4 | 0 | 3 |
| TDH2 | P00358 | Glyceraldehyde-3-phosphate dehydrogenase 2 | 332 | 11.14 | 3 | 0 | 10 |
| TDH3 | P00359 | Glyceraldehyde-3-phosphate dehydrogenase 3 | 332 | 11.14 | 3 | 0 | 10 |
| TDH1 | P00360 | Glyceraldehyde-3-phosphate dehydrogenase 1 | 332 | 6.93 | 2 | 0 | 7 |
| CDC19 | P00549 | Pyruvate kinase 1 | 500 | 7.40 | 4 | 0 | 8 |
|  | P00761 | Trypsin | 231 | 25.11 | 4 | 4 | 338 |
| ALDOA | P00883 | Fructose-bisphosphate aldolase A | 364 | 8.24 | 1 | 1 | 11 |
| TPI1 | P00942 | Triosephosphate isomerase | 248 | 2.82 | 2 | 0 | 4 |
| GPM1 | P00950 | Phosphoglycerate mutase 1 | 247 | 11.74 | 4 | 0 | 6 |
| CSN2 | P02666 | Beta-casein | 224 | 3.12 | 2 | 2 | 1 |
| CSN3 | P02668 | Kappa-casein | 190 | 5.26 | 1 | 1 | 7 |
| ALB | P02768 | Albumin | 609 | 1.15 | 2 | 1 | 2 |
| ALB | P02769 | Albumin | 607 | 1.15 | 2 | 1 | 2 |
| TEF1 | P02994 | Elongation factor 1-alpha | 458 | 1.97 | 2 | 0 | 2 |
| CAT | P04040 | Catalase | 527 | 2.47 | 2 | 2 | 1 |
| KRT1 | P04264 | Keratin, type II cytoskeletal 1 | 644 | 40.84 | 29 | 27 | 179 |
| POR1 | P04840 | Mitochondrial outer membrane protein porin 1 | 283 | 23.32 | 6 | 0 | 24 |
| RPP0 | P05317 | Large ribosomal subunit protein uL10 | 312 | 3.85 | 1 | 0 | 3 |
| RPS3 | P05750 | Small ribosomal subunit protein uS3 | 240 | 42.92 | 9 | 0 | 51 |
| RPS13 | P05756 | Small ribosomal subunit protein uS15 | 151 | 7.28 | 1 | 0 | 1 |
| PRB1 | P09232 | Cerevisin | 635 | 3.94 | 2 | 0 | 8 |
| RPL11A | P0C0W9 | Large ribosomal subunit protein uL5A | 174 | 8.05 | 2 | 0 | 7 |
| RPS4A | P0CX35 | Small ribosomal subunit protein eS4A | 261 | 21.07 | 6 | 0 | 33 |
| RPS6A | P0CX37 | Small ribosomal subunit protein eS6A | 236 | 13.98 | 4 | 0 | 12 |
| RPL1A | P0CX43 | Large ribosomal subunit protein uL1A | 217 | 3.69 | 1 | 0 | 1 |
| RPL2A | P0CX45 | Large ribosomal subunit protein uL2A | 254 | 4.33 | 1 | 0 | 1 |
| RPL12A | P0CX53 | Large ribosomal subunit protein uL11A | 165 | 9.09 | 1 | 0 | 2 |
| RPL19A | P0CX82 | Large ribosomal subunit protein eL19A | 189 | 8.99 | 3 | 0 | 6 |

| Gene | Protein ID | Protein description | Length (aa) | Coverage (%) | Total peptides | Unique peptides | Total spectral count |
|---|---|---|---|---|---|---|---|
| SSA1 | P10591 | Heat shock protein SSA1 | 642 | 1.87 | 3 | 0 | 4 |
| SSA2 | P10592 | Heat shock protein SSA2 | 639 | 1.88 | 2 | 0 | 4 |
| SSB1 | P11484 | Ribosome-associated molecular chaperone SSB1 | 613 | 4.89 | 4 | 0 | 7 |
| KRT10 | P13645 | Keratin, type I cytoskeletal 10 | 584 | 34.08 | 21 | 16 | 93 |
| DPM1 | P14020 | Dolichol-phosphate mannosyltransferase | 267 | 4.12 | 1 | 0 | 1 |
| BGL2 | P15703 | Glucan 1,3-beta-glucosidase | 313 | 3.83 | 1 | 0 | 1 |
| KAR2 | P16474 | Endoplasmic reticulum chaperone BiP | 682 | 1.76 | 1 | 0 | 4 |
| HSP60 | P19882 | Heat shock protein 60, mitochondrial | 572 | 5.77 | 5 | 0 | 7 |
| SSA4 | P22202 | Heat shock protein SSA4 | 642 | 1.87 | 1 | 0 | 4 |
| VMA4 | P22203 | V-type proton ATPase subunit E | 233 | 12.02 | 4 | 0 | 10 |
| RPS2 | P25443 | Small ribosomal subunit protein uS5 | 254 | 9.06 | 2 | 0 | 2 |
| BMH1 | P29311 | Protein BMH1 | 267 | 8.61 | 3 | 0 | 5 |
| RPL8B | P29453 | Large ribosomal subunit protein eL8B | 256 | 14.45 | 4 | 0 | 23 |
| EFT1 | P32324 | Elongation factor 2 | 842 | 1.07 | 2 | 0 | 2 |
| MRP20 | P32387 | Large ribosomal subunit protein uL23m | 263 | 3.04 | 1 | 0 | 6 |
| EFB1 | P32471 | Elongation factor 1-beta | 206 | 18.93 | 4 | 0 | 7 |
| RPN12 | P32496 | 26S proteasome regulatory subunit RPN12 | 274 | 2.92 | 1 | 0 | 5 |
| VMA8 | P32610 | V-type proton ATPase subunit D | 256 | 9.38 | 3 | 0 | 9 |
| GSP1 | P32835 | GTP-binding nuclear protein GSP1/CNR1 | 219 | 10.96 | 2 | 0 | 5 |
| BMH2 | P34730 | Protein BMH2 | 273 | 4.03 | 1 | 0 | 1 |
| TSA1 | P34760 | Peroxiredoxin TSA1 | 196 | 5.10 | 1 | 0 | 1 |
| KRT9 | P35527 | Keratin, type I cytoskeletal 9 | 623 | 27.93 | 18 | 17 | 64 |
| KRT2 | P35908 | Keratin, type II cytoskeletal 2 epidermal | 639 | 43.51 | 27 | 24 | 127 |
| ASC1 | P38011 | Small ribosomal subunit protein RACK1 | 319 | 2.82 | 1 | 0 | 2 |
| ATP3 | P38077 | ATP synthase subunit gamma, mitochondrial | 311 | 6.43 | 2 | 0 | 5 |
| RVS167 | P39743 | Reduced viability upon starvation protein 167 | 482 | 1.45 | 1 | 0 | 7 |
| GPP2 | P40106 | Glycerol-1-phosphate phosphohydrolase 2 | 250 | 20.00 | 5 | 0 | 14 |
| SSB2 | P40150 | Ribosome-associated molecular chaperone SSB2 | 613 | 4.89 | 4 | 0 | 7 |
| RPL13B | P40212 | Large ribosomal subunit protein eL13B | 199 | 5.03 | 1 | 0 | 1 |
| GPP1 | P41277 | Glycerol-1-phosphate phosphohydrolase 1 | 250 | 22.00 | 5 | 0 | 23 |
| CIR1 | P42940 | Probable electron transfer flavoprotein subunit beta | 261 | 16.48 | 4 | 0 | 17 |
| MOG1 | P47123 | Nuclear import protein MOG1 | 218 | 24.31 | 5 | 0 | 143 |
| MRPL22 | P53881 | Large ribosomal subunit protein uL22m | 309 | 2.59 | 1 | 0 | 1 |
| ACT1 | P60010 | Actin | 375 | 1.87 | 2 | 0 | 1 |
| LYZ | P61626 | Lysozyme C | 148 | 13.51 | 2 | 2 | 2 |
| KRT85 | P78386 | Keratin, type II cuticular Hb5 | 507 | 2.96 | 3 | 1 | 3 |
| SEC13 | Q04491 | Protein transport protein SEC13 | 297 | 7.07 | 2 | 0 | 8 |
| KRT33B | Q14525 | Keratin, type I cuticular Ha3-II | 404 | 3.96 | 3 | 0 | 6 |

| Gene | Protein ID | Protein description | Length (aa) | Coverage (%) | Total peptides | Unique peptides | Total spectral count |
|---|---|---|---|---|---|---|---|
| KRT31 | Q15323 | Keratin, type I cuticular Ha1 | 416 | 3.85 | 4 | 1 | 6 |
| KRT35 | Q92764 | Keratin, type I cuticular Ha5 | 455 | 1.54 | 2 | 0 | 3 |
| KRT84 | Q9NSB2 | Keratin, type II cuticular Hb4 | 600 | 2.33 | 2 | 0 | 3 |
| AWRI1631_162870 | B5VTH7 | DNA replication licensing factor MCM4 (Fragment) | 883 | 1.47 | 1 | 1 | 1 |
|  | P02534 | Keratin, type I microfibrillar 48 kDa, component 8C-1 | 412 | 3.88 | 3 | 0 | 6 |
| EC1118_1J19_0188g | C8ZBM1 | Mog1p | 188 | 43.09 | 6 | 1 | 148 |
| K7_RPL13A | G2WC81 | 60S ribosomal protein L13 | 199 | 5.03 | 1 | 0 | 1 |
| RPL13A | Q12690 | Large ribosomal subunit protein eL13A | 199 | 5.03 | 1 | 0 | 1 |
| PYK2 | P52489 | Pyruvate kinase 2 | 506 | 1.98 | 1 | 0 | 2 |
|  | P02441 | Keratin, high-sulfur matrix protein, IIIA3 | 131 | 7.63 | 1 | 0 | 1 |
| YJR056C | P47115 | Uncharacterized protein YJR056C | 236 | 4.24 | 1 | 0 | 3 |
| MGM101 | P32787 | Mitochondrial genome maintenance protein MGM101 | 269 | 3.35 | 1 | 0 | 2 |
| PRO3 | P32263 | Pyrroline-5-carboxylate reductase | 286 | 4.55 | 1 | 0 | 2 |

#### Sheet 3. Upper Spt5 band (86 proteins)

| Gene | Protein ID | Protein description | Length (aa) | Coverage (%) | Total peptides | Unique peptides | Total spectral count |
|---|---|---|---|---|---|---|---|
| DED1 | A6ZP47 | ATP-dependent RNA helicase DED1 | 604 | 2.98 | 2 | 0 | 5 |
| RPS0A | A6ZUM5 | Small ribosomal subunit protein uS2A | 252 | 3.17 | 1 | 0 | 3 |
| RPS0B | A7A0V3 | Small ribosomal subunit protein uS2B | 252 | 3.17 | 1 | 0 | 3 |
| KRT86 | O43790 | Keratin, type II cuticular Hb6 | 486 | 1.44 | 1 | 0 | 1 |
| KRT15 | O77727 | Keratin, type I cytoskeletal 15 | 453 | 10.38 | 5 | 2 | 18 |
| ADH1 | P00330 | Alcohol dehydrogenase 1 | 348 | 4.31 | 2 | 0 | 7 |
| TDH2 | P00358 | Glyceraldehyde-3-phosphate dehydrogenase 2 | 332 | 6.93 | 2 | 0 | 9 |
| TDH3 | P00359 | Glyceraldehyde-3-phosphate dehydrogenase 3 | 332 | 6.93 | 2 | 0 | 9 |
| TDH1 | P00360 | Glyceraldehyde-3-phosphate dehydrogenase 1 | 332 | 6.93 | 2 | 0 | 9 |
| CDC19 | P00549 | Pyruvate kinase 1 | 500 | 5.40 | 3 | 0 | 6 |
| PGK1 | P00560 | Phosphoglycerate kinase | 416 | 2.40 | 1 | 0 | 5 |
|  | P00761 | Trypsin | 231 | 32.90 | 5 | 5 | 566 |
| ALDOA | P00883 | Fructose-bisphosphate aldolase A | 364 | 8.24 | 1 | 1 | 16 |
| CSN1S2 | P02663 | Alpha-S2-casein | 222 | 8.56 | 2 | 2 | 2 |
| ALB | P02768 | Albumin | 609 | 10.34 | 7 | 5 | 17 |
| ALB | P02769 | Albumin | 607 | 2.31 | 2 | 0 | 5 |
| LTF | P02788 | Lactotransferrin | 710 | 2.54 | 2 | 2 | 6 |
| TEF1 | P02994 | Elongation factor 1-alpha | 458 | 9.17 | 6 | 0 | 20 |
| CAT | P04040 | Catalase | 527 | 1.71 | 1 | 1 | 1 |
| PAB1 | P04147 | Polyadenylate-binding protein, cytoplasmic and nuclear | 577 | 1.73 | 1 | 0 | 4 |
| KRT1 | P04264 | Keratin, type II cytoskeletal 1 | 644 | 46.89 | 34 | 31 | 187 |
| RPS31 | P05759 | Ubiquitin-ribosomal protein eS31 fusion protein | 152 | 5.92 | 1 | 0 | 3 |
| PDC1 | P06169 | Pyruvate decarboxylase isozyme 1 | 563 | 1.78 | 2 | 0 | 5 |
| RAD2 | P07276 | DNA repair protein RAD2 | 1031 | 3.49 | 4 | 0 | 22 |
| RPB2 | P08518 | DNA-directed RNA polymerase II subunit RPB2 | 1224 | 4.66 | 6 | 0 | 18 |
| RPL19A | P0CX82 | Large ribosomal subunit protein eL19A | 189 | 10.05 | 3 | 0 | 5 |
| SSA1 | P10591 | Heat shock protein SSA1 | 642 | 1.25 | 1 | 0 | 5 |
| SSA2 | P10592 | Heat shock protein SSA2 | 639 | 1.25 | 1 | 0 | 5 |
| PYC1 | P11154 | Pyruvate carboxylase 1 | 1178 | 2.12 | 3 | 0 | 3 |
| SSB1 | P11484 | Ribosome-associated molecular chaperone SSB1 | 613 | 6.85 | 4 | 0 | 6 |
| KRT10 | P13645 | Keratin, type I cytoskeletal 10 | 584 | 39.21 | 22 | 18 | 109 |
| FBA1 | P14540 | Fructose-bisphosphate aldolase | 359 | 3.34 | 1 | 0 | 1 |
| YEF3 | P16521 | Elongation factor 3A | 1044 | 1.15 | 1 | 0 | 2 |
| GAL11 | P19659 | Mediator of RNA polymerase II transcription subunit 15 | 1081 | 1.20 | 1 | 0 | 2 |
| HSP60 | P19882 | Heat shock protein 60, mitochondrial | 572 | 2.97 | 1 | 0 | 1 |
| RPA135 | P22138 | DNA-directed RNA polymerase I subunit RPA135 | 1203 | 3.41 | 4 | 0 | 16 |
| SYP1 | P25623 | Suppressor of yeast profilin deletion | 870 | 1.03 | 1 | 0 | 1 |

| Gene | Protein ID | Protein description | Length (aa) | Coverage (%) | Total peptides | Unique peptides | Total spectral count |
|---|---|---|---|---|---|---|---|
| CDC48 | P25694 | Cell division control protein 48 | 835 | 1.56 | 1 | 0 | 1 |
| CDC60 | P26637 | Leucine--tRNA ligase, cytoplasmic | 1090 | 4.86 | 5 | 0 | 18 |
| SPT5 | P27692 | Transcription elongation factor SPT5 | 1063 | 41.58 | 49 | 0 | 240 |
| FUN30 | P31380 | ATP-dependent helicase FUN30 | 1131 | 0.88 | 1 | 0 | 1 |
| EFT1 | P32324 | Elongation factor 2 | 842 | 5.11 | 4 | 0 | 9 |
| SSE1 | P32589 | Heat shock protein homolog SSE1 | 693 | 2.74 | 2 | 0 | 3 |
| BEM3 | P32873 | GTPase-activating protein BEM3 | 1128 | 4.26 | 5 | 0 | 24 |
| KRT9 | P35527 | Keratin, type I cytoskeletal 9 | 623 | 38.52 | 21 | 20 | 124 |
| KRT2 | P35908 | Keratin, type II cytoskeletal 2 epidermal | 639 | 47.89 | 28 | 24 | 115 |
| PIM1 | P36775 | Lon protease homolog, mitochondrial | 1133 | 7.59 | 10 | 0 | 32 |
| ATP3 | P38077 | ATP synthase subunit gamma, mitochondrial | 311 | 4.50 | 1 | 0 | 1 |
| AKL1 | P38080 | Serine/threonine-protein kinase AKL1 | 1108 | 2.35 | 3 | 0 | 6 |
| ROT2 | P38138 | Glucosidase 2 subunit alpha | 954 | 1.78 | 2 | 0 | 2 |
| TSL1 | P38427 | Trehalose synthase complex regulatory subunit TSL1 | 1098 | 1.37 | 2 | 0 | 6 |
| TIF4631 | P39935 | Eukaryotic initiation factor 4F subunit p150 | 952 | 29.83 | 32 | 0 | 177 |
| TIF4632 | P39936 | Eukaryotic initiation factor 4F subunit p130 | 914 | 1.53 | 2 | 0 | 8 |
| RPS26A | P39938 | Small ribosomal subunit protein eS26A | 119 | 7.56 | 1 | 0 | 1 |
| BOI2 | P39969 | Protein BOI2 | 1040 | 2.02 | 2 | 0 | 12 |
| SSB2 | P40150 | Ribosome-associated molecular chaperone SSB2 | 613 | 6.85 | 4 | 0 | 6 |
| MKT1 | P40850 | Post-transcriptional regulator MKT1 | 830 | 2.89 | 1 | 0 | 1 |
| ALY2 | P47029 | Arrestin-related trafficking adapter 3 | 1046 | 1.05 | 1 | 0 | 3 |
| JSN1 | P47135 | Protein JSN1 | 1091 | 4.12 | 5 | 0 | 8 |
| SEC16 | P48415 | COPII coat assembly protein SEC16 | 2195 | 0.73 | 1 | 0 | 1 |
| NUP145 | P49687 | Nucleoporin NUP145 | 1317 | 0.76 | 1 | 0 | 1 |
| COP1 | P53622 | Coatomer subunit alpha | 1201 | 1.33 | 2 | 0 | 7 |
| ACT1 | P60010 | Actin | 375 | 5.87 | 2 | 0 | 8 |
| LYZ | P61626 | Lysozyme C | 148 | 24.32 | 4 | 4 | 23 |
| KRT85 | P78386 | Keratin, type II cuticular Hb5 | 507 | 1.38 | 1 | 0 | 1 |
| NAB6 | Q03735 | RNA-binding protein NAB6 | 1134 | 11.11 | 13 | 0 | 42 |
| UTP14 | Q04500 | U3 small nucleolar RNA-associated protein 14 | 899 | 1.33 | 1 | 0 | 1 |
| KAP95 | Q06142 | Importin subunit beta-1 | 861 | 1.16 | 1 | 0 | 1 |
| RGA2 | Q06407 | Rho-type GTPase-activating protein 2 | 1009 | 0.89 | 1 | 0 | 4 |
| SYT1 | Q06836 | Arf guanine nucleotide exchange factor SYT1 | 1226 | 1.22 | 1 | 0 | 2 |
| YPR097W | Q06839 | PX domain-containing protein YPR097W | 1073 | 0.84 | 1 | 0 | 1 |
| NEW1 | Q08972 | [NU+] prion formation protein 1 | 1196 | 9.45 | 12 | 0 | 48 |
| PUF2 | Q12221 | mRNA-binding protein PUF2 | 1075 | 0.84 | 1 | 0 | 1 |
| KRT33B | Q14525 | Keratin, type I cuticular Ha3-II | 404 | 1.73 | 1 | 0 | 8 |
| KRT31 | Q15323 | Keratin, type I cuticular Ha1 | 416 | 1.68 | 1 | 0 | 8 |

| Gene | Protein ID | Protein description | Length (aa) | Coverage (%) | Total peptides | Unique peptides | Total spectral count |
|---|---|---|---|---|---|---|---|
| KRT35 | Q92764 | Keratin, type I cuticular Ha5 | 455 | 1.54 | 1 | 0 | 8 |
| DUF1 | Q99247 | DUB-associated factor 1 | 1116 | 1.79 | 2 | 0 | 3 |
| KRT84 | Q9NSB2 | Keratin, type II cuticular Hb4 | 600 | 2.67 | 2 | 0 | 4 |
| | P02534 | Keratin, type I microfibrillar 48 kDa, component 8C-1 | 412 | 1.70 | 1 | 0 | 8 |
| RPS27A | P62979 | Ubiquitin-40S ribosomal protein S27a | 156 | 5.77 | 1 | 0 | 3 |
| SCRG_00930 | B3LID9 | branched-chain-2-oxoacid decarboxylase | 563 | 1.78 | 1 | 0 | 5 |
| AWRI1631_73850 | B5VJA5 | YGR162Wp-like protein | 842 | 31.59 | 30 | 1 | 152 |
| ECM30 | Q06673 | Protein ECM30 | 1274 | 0.94 | 1 | 0 | 4 |
| KIN1 | P13185 | Serine/threonine protein kinase KIN1 | 1064 | 0.85 | 1 | 0 | 2 |
| ECM21 | P38167 | Protein ECM21 | 1117 | 0.98 | 1 | 0 | 1 |
| LEO1 | P38439 | RNA polymerase-associated protein LEO1 | 464 | 1.72 | 1 | 0 | 6 |

#### Sheet 4. Lower Spt5 band (120 proteins)

| Gene | Protein ID | Protein description | Length (aa) | Coverage (%) | Total peptides | Unique peptides | Total spectral count |
|---|---|---|---|---|---|---|---|
| DED1 | A6ZP47 | ATP-dependent RNA helicase DED1 | 604 | 5.63 | 7 | 0 | 7 |
| DCP2 | A6ZRW5 | mRNA-decapping enzyme subunit 2 | 970 | 3.30 | 3 | 0 | 15 |
| RPS0A | A6ZUM5 | Small ribosomal subunit protein uS2A | 252 | 7.94 | 2 | 0 | 4 |
| RPS0B | A7A0V3 | Small ribosomal subunit protein uS2B | 252 | 7.94 | 2 | 0 | 4 |
| EIS1 | C7GL88 | Eisosome protein 1 | 843 | 3.32 | 4 | 0 | 13 |
| KRT86 | O43790 | Keratin, type II cuticular Hb6 | 486 | 1.44 | 1 | 0 | 2 |
| KRT15 | O77727 | Keratin, type I cytoskeletal 15 | 453 | 9.27 | 6 | 1 | 14 |
| ADH1 | P00330 | Alcohol dehydrogenase 1 | 348 | 7.18 | 5 | 0 | 9 |
| TDH2 | P00358 | Glyceraldehyde-3-phosphate dehydrogenase 2 | 332 | 12.35 | 5 | 0 | 13 |
| TDH3 | P00359 | Glyceraldehyde-3-phosphate dehydrogenase 3 | 332 | 12.35 | 5 | 0 | 13 |
| TDH1 | P00360 | Glyceraldehyde-3-phosphate dehydrogenase 1 | 332 | 12.35 | 4 | 0 | 13 |
| CDC19 | P00549 | Pyruvate kinase 1 | 500 | 12.00 | 7 | 0 | 27 |
|  | P00761 | Trypsin | 231 | 38.96 | 6 | 6 | 661 |
| ALDOA | P00883 | Fructose-bisphosphate aldolase A | 364 | 8.24 | 1 | 1 | 7 |
| ENO1 | P00924 | Enolase 1 | 437 | 2.75 | 1 | 0 | 2 |
| ENO2 | P00925 | Enolase 2 | 437 | 2.75 | 1 | 0 | 2 |
| CSN1S1 | P02662 | Alpha-S1-casein | 214 | 5.61 | 1 | 1 | 2 |
| CSN1S2 | P02663 | Alpha-S2-casein | 222 | 17.12 | 5 | 5 | 9 |
| CSN2 | P02666 | Beta-casein | 224 | 3.12 | 1 | 1 | 6 |
| CSN3 | P02668 | Kappa-casein | 190 | 9.47 | 2 | 2 | 11 |
| ALB | P02768 | Albumin | 609 | 15.76 | 11 | 8 | 32 |
| ALB | P02769 | Albumin | 607 | 2.31 | 3 | 0 | 6 |
| LTF | P02788 | Lactotransferrin | 710 | 5.63 | 4 | 4 | 16 |
| HSP82 | P02829 | ATP-dependent molecular chaperone HSP82 | 709 | 1.27 | 1 | 0 | 1 |
| TEF1 | P02994 | Elongation factor 1-alpha | 458 | 12.66 | 8 | 0 | 22 |
| CPA2 | P03965 | Carbamoyl-phosphate synthase arginine-specific large chain | 1118 | 2.77 | 3 | 0 | 8 |
| PAB1 | P04147 | Polyadenylate-binding protein, cytoplasmic and nuclear | 577 | 6.76 | 4 | 0 | 7 |
| KRT1 | P04264 | Keratin, type II cytoskeletal 1 | 644 | 40.99 | 30 | 27 | 163 |
| SUP35 | P05453 | Eukaryotic peptide chain release factor GTP-binding subunit | 685 | 1.46 | 2 | 0 | 1 |
| RPS31 | P05759 | Ubiquitin-ribosomal protein eS31 fusion protein | 152 | 5.92 | 1 | 0 | 2 |
| PDC1 | P06169 | Pyruvate decarboxylase isozyme 1 | 563 | 7.82 | 4 | 0 | 14 |
| URA2 | P07259 | Protein URA2 | 2214 | 0.99 | 2 | 0 | 4 |
| VAS1 | P07806 | Valine--tRNA ligase, mitochondrial | 1104 | 4.98 | 6 | 0 | 32 |
| ILS1 | P09436 | Isoleucine--tRNA ligase, cytoplasmic | 1072 | 5.78 | 7 | 0 | 29 |
| RPL18A | P0CX49 | Large ribosomal subunit protein eL18A | 186 | 4.30 | 1 | 0 | 2 |
| RPL19A | P0CX82 | Large ribosomal subunit protein eL19A | 189 | 8.99 | 2 | 0 | 7 |
| SSA1 | P10591 | Heat shock protein SSA1 | 642 | 1.25 | 3 | 0 | 4 |

| Gene | Protein ID | Protein description | Length (aa) | Coverage (%) | Total peptides | Unique peptides | Total spectral count |
|---|---|---|---|---|---|---|---|
| SSA2 | P10592 | Heat shock protein SSA2 | 639 | 1.25 | 3 | 0 | 4 |
| SSB1 | P11484 | Ribosome-associated molecular chaperone SSB1 | 613 | 17.46 | 11 | 0 | 31 |
| PDR1 | P12383 | Transcription factor PDR1 | 1068 | 2.81 | 3 | 0 | 5 |
| LAT1 | P12695 | Dihydrolipoyllysine-residue acetyltransferase component of pyruvate dehydrogenase complex, mitochondrial | 482 | 1.66 | 1 | 0 | 3 |
| KRT10 | P13645 | Keratin, type I cytoskeletal 10 | 584 | 41.10 | 26 | 20 | 94 |
| HSC82 | P15108 | ATP-dependent molecular chaperone HSC82 | 705 | 1.28 | 1 | 0 | 1 |
| STI1 | P15705 | Heat shock protein STI1 | 589 | 2.21 | 1 | 0 | 1 |
| YEF3 | P16521 | Elongation factor 3A | 1044 | 19.16 | 20 | 0 | 49 |
| PFK1 | P16861 | ATP-dependent 6-phosphofructokinase subunit alpha | 987 | 1.01 | 1 | 0 | 3 |
| VMA1 | P17255 | V-type proton ATPase catalytic subunit A | 1071 | 1.40 | 2 | 0 | 3 |
| SNF5 | P18480 | SWI/SNF chromatin-remodeling complex subunit SNF5 | 905 | 1.44 | 1 | 0 | 3 |
| RGR1 | P19263 | Mediator of RNA polymerase II transcription subunit 14 | 1082 | 2.68 | 3 | 0 | 4 |
| GAL11 | P19659 | Mediator of RNA polymerase II transcription subunit 15 | 1081 | 2.50 | 2 | 0 | 10 |
| HSP60 | P19882 | Heat shock protein 60, mitochondrial | 572 | 5.94 | 3 | 0 | 10 |
| UBA1 | P22515 | Ubiquitin-activating enzyme E1 1 | 1024 | 4.88 | 5 | 0 | 11 |
| SPB1 | P25582 | 27S pre-rRNA (guanosine(2922)-2'-O)-methyltransferase | 841 | 1.07 | 1 | 0 | 2 |
| SYP1 | P25623 | Suppressor of yeast profilin deletion | 870 | 2.41 | 2 | 0 | 2 |
| SPT5 | P27692 | Transcription elongation factor SPT5 | 1063 | 32.55 | 33 | 0 | 118 |
| EFT1 | P32324 | Elongation factor 2 | 842 | 6.18 | 6 | 0 | 20 |
| PAN1 | P32521 | Actin cytoskeleton-regulatory complex protein PAN1 | 1480 | 0.74 | 1 | 0 | 2 |
| SPT16 | P32558 | FACT complex subunit SPT16 | 1035 | 1.45 | 2 | 0 | 2 |
| SSE1 | P32589 | Heat shock protein homolog SSE1 | 693 | 2.74 | 2 | 0 | 11 |
| SWI3 | P32591 | SWI/SNF complex subunit SWI3 | 825 | 7.03 | 6 | 0 | 17 |
| KRT9 | P35527 | Keratin, type I cytoskeletal 9 | 623 | 20.87 | 13 | 12 | 53 |
| DEF1 | P35732 | RNA polymerase II degradation factor 1 | 738 | 3.93 | 3 | 0 | 5 |
| KRT2 | P35908 | Keratin, type II cytoskeletal 2 epidermal | 639 | 45.07 | 30 | 26 | 114 |
| PIM1 | P36775 | Lon protease homolog, mitochondrial | 1133 | 2.47 | 4 | 0 | 12 |
| APE3 | P37302 | Aminopeptidase Y | 537 | 1.30 | 1 | 0 | 1 |
| ROT2 | P38138 | Glucosidase 2 subunit alpha | 954 | 6.60 | 7 | 0 | 11 |
| RPG1 | P38249 | Eukaryotic translation initiation factor 3 subunit A | 964 | 19.61 | 20 | 0 | 62 |
| TPS3 | P38426 | Trehalose synthase complex regulatory subunit TPS3 | 1054 | 9.68 | 10 | 0 | 27 |
| TSL1 | P38427 | Trehalose synthase complex regulatory subunit TSL1 | 1098 | 1.37 | 3 | 0 | 2 |
| KSP1 | P38691 | Serine/threonine-protein kinase KSP1 | 1029 | 0.97 | 1 | 0 | 1 |
| RPN1 | P38764 | 26S proteasome regulatory subunit RPN1 | 993 | 1.61 | 2 | 0 | 6 |
| PIK1 | P39104 | Phosphatidylinositol 4-kinase PIK1 | 1066 | 3.66 | 4 | 0 | 14 |
| TIF4631 | P39935 | Eukaryotic initiation factor 4F subunit p150 | 952 | 16.70 | 20 | 0 | 86 |
| TIF4632 | P39936 | Eukaryotic initiation factor 4F subunit p130 | 914 | 12.91 | 12 | 0 | 44 |

| Gene | Protein ID | Protein description | Length (aa) | Coverage (%) | Total peptides | Unique peptides | Total spectral count |
|---|---|---|---|---|---|---|---|
| RPS26A | P39938 | Small ribosomal subunit protein eS26A | 119 | 7.56 | 1 | 0 | 1 |
| POL5 | P39985 | rDNA transcriptional regulator POL5 | 1022 | 6.07 | 6 | 0 | 26 |
| KAP123 | P40069 | Importin subunit beta-4 | 1113 | 2.07 | 2 | 0 | 4 |
| SSB2 | P40150 | Ribosome-associated molecular chaperone SSB2 | 613 | 15.33 | 10 | 0 | 30 |
| GUS1 | P46655 | Glutamate--tRNA ligase, cytoplasmic | 708 | 2.54 | 2 | 0 | 7 |
| FYV8 | P46949 | Protein FYV8 | 817 | 2.45 | 2 | 0 | 6 |
| URB2 | P47108 | Nucleolar pre-ribosomal-associated protein 2 | 1174 | 2.73 | 3 | 0 | 3 |
| JSN1 | P47135 | Protein JSN1 | 1091 | 9.35 | 9 | 0 | 24 |
| SEC16 | P48415 | COPII coat assembly protein SEC16 | 2195 | 3.74 | 8 | 0 | 17 |
| MCM6 | P53091 | DNA replication licensing factor MCM6 | 1017 | 1.28 | 1 | 0 | 5 |
| YGR130C | P53278 | Uncharacterized protein YGR130C | 816 | 4.29 | 4 | 0 | 16 |
| BNI4 | P53858 | Protein BNI4 | 892 | 6.17 | 6 | 0 | 20 |
| ACT1 | P60010 | Actin | 375 | 8.53 | 3 | 0 | 7 |
| LYZ | P61626 | Lysozyme C | 148 | 24.32 | 4 | 4 | 12 |
| KRT85 | P78386 | Keratin, type II cuticular Hb5 | 507 | 1.38 | 1 | 0 | 2 |
| ULP1 | Q02724 | Ubiquitin-like-specific protease 1 | 621 | 1.13 | 1 | 0 | 1 |
| SVL3 | Q03088 | Styryl dye vacuolar localization protein 3 | 825 | 1.21 | 1 | 0 | 1 |
| STE20 | Q03497 | Serine/threonine-protein kinase STE20 | 939 | 4.58 | 3 | 0 | 11 |
| NAB6 | Q03735 | RNA-binding protein NAB6 | 1134 | 7.76 | 11 | 0 | 19 |
| MRD1 | Q06106 | Multiple RNA-binding domain-containing protein 1 | 887 | 6.99 | 7 | 0 | 19 |
| KAP95 | Q06142 | Importin subunit beta-1 | 861 | 2.32 | 2 | 0 | 4 |
| IMH1 | Q06704 | Golgin IMH1 | 911 | 2.09 | 2 | 0 | 3 |
| TSR1 | Q07381 | Ribosome biogenesis protein TSR1 | 788 | 1.02 | 1 | 0 | 1 |
| DIS3 | Q08162 | Exosome complex exonuclease DIS3 | 1001 | 2.80 | 4 | 0 | 9 |
| PUF2 | Q12221 | mRNA-binding protein PUF2 | 1075 | 1.86 | 2 | 0 | 4 |
| KRT33B | Q14525 | Keratin, type I cuticular Ha3-II | 404 | 1.73 | 1 | 0 | 4 |
| KRT31 | Q15323 | Keratin, type I cuticular Ha1 | 416 | 1.68 | 1 | 0 | 4 |
| KRT35 | Q92764 | Keratin, type I cuticular Ha5 | 455 | 1.54 | 1 | 0 | 4 |
| KRT84 | Q9NSB2 | Keratin, type II cuticular Hb4 | 600 | 2.33 | 2 | 0 | 17 |
| NOG1 | A0A8H4F6Y2 | Nucleolar GTP-binding protein 1 | 647 | 1.39 | 1 | 1 | 1 |
|  | P02534 | Keratin, type I microfibrillar 48 kDa, component 8C-1 | 412 | 1.70 | 1 | 0 | 4 |
| RPS27A | P62979 | Ubiquitin-40S ribosomal protein S27a | 156 | 5.77 | 1 | 0 | 2 |
| SCRG_00677 | B3LH69 | H(+)-transporting two-sector ATPase | 617 | 2.43 | 2 | 0 | 3 |
| SCRG_00930 | B3LID9 | branched-chain-2-oxoacid decarboxylase | 563 | 1.78 | 1 | 0 | 6 |
| AWRI1631_73850 | B5VJA5 | YGR162Wp-like protein | 842 | 18.29 | 19 | 1 | 79 |
| K7_LAT1 | G2WM33 | Acetyltransferase component of pyruvate dehydrogenase complex | 482 | 1.66 | 1 | 0 | 3 |
| HTZ1 | Q12692 | Histone H2A.Z | 134 | 6.72 | 1 | 0 | 4 |

| Gene | Protein ID | Protein description | Length (aa) | Coverage (%) | Total peptides | Unique peptides | Total spectral count |
|---|---|---|---|---|---|---|---|
| PYK2 | P52489 | Pyruvate kinase 2 | 506 | 1.98 | 1 | 0 | 3 |
| YOL098C | Q12496 | Uncharacterized protein YOL098C | 1037 | 0.96 | 1 | 0 | 4 |
| PRP9 | A6ZXU1 | RNA splicing factor | 530 | 1.32 | 1 | 0 | 1 |
| NNK1 | P36003 | Nitrogen network kinase 1 | 928 | 0.97 | 1 | 0 | 1 |
| CTSG | P08311 | Cathepsin G | 255 | 3.53 | 1 | 1 | 2 |
| BOI1 | P38041 | Protein BOB1 | 980 | 0.92 | 1 | 0 | 2 |
| NUP133 | P36161 | Nucleoporin NUP133 | 1157 | 1.04 | 1 | 0 | 1 |
| VRP1 | P37370 | Verprolin | 817 | 1.10 | 1 | 0 | 1 |
| DAL81 | P21657 | Transcriptional activator protein DAL81 | 970 | 0.72 | 1 | 0 | 1 |
